# Molecular basis of mitogen-activated protein kinase ERK2 activation by its upstream kinase MEK1

**DOI:** 10.64898/2026.01.19.700303

**Authors:** Jill von Velsen, Pauline Juyoux, Nicola Piasentin, Hayden Fisher, Karine Lapouge, Oscar Vadas, Francesco Luigi Gervasio, Matthew W. Bowler

## Abstract

The RAS-RAF-MEK-ERK mitogen-activated protein kinase (MAPK) pathway relays extracellular signals into a cellular response and its dysregulation leads to many pathologies, particularly cancer. Much is known about the individual components of the pathway, however, details of their interactions and how specificity is maintained is still poorly understood. Here, we determined cryo-EM structures of the MAP2K MEK1 activating its substrate MAPK ERK2, the final event in the cascade. We define the molecular details of specificity and activation of ERK2 and examine the mechanism of substrate recognition using solution techniques and molecular dynamics. Binding of the substrate MAPK leads to release of the MAP2K catalytic machinery, a mechanism enhanced in many gain of function disease mutations. Further, we observe that ERK2 release is not required for nucleotide exchange, suggesting how a processive mechanism could proceed. Our data advance the understanding of MAPK signalling specificity and provide a starting point for drug development.

**One-Sentence Summary:** Cryo-EM structures of the MEK1-ERK2 complex reveal details of cellular signal transmission.

## Main text

Cells need to respond to their environment in order to adapt and react (*1*). Numerous signalling cascades have evolved in eukaryotes to transmit signals or external conditions, from the cell surface into the cell, thereby eliciting a response (*2*). The mitogen-activated protein kinases (MAPKs) are a family of highly conserved kinases (*3*) responsible for cellular reactions elicited by extracellular signals from stress, mitogens, cytokines or growth factors and include the extracellular signal-regulated kinases (ERK1 and 2), the final serine/threonine kinases in the RAS-RAF-MEK-ERK branch of the pathways (*4, 5*). The ERKs have a wide range of substrates that they can activate, including transcription factors and protein kinases, leading to responses such as cell growth, differentiation, and proliferation. The key role of the pathway in regulating cell behavior leads to its dysfunction being implicated in many diseases, with around 30% of cancers carrying mutations in components of the pathway (*6, 7*).

The cascades involve multiple kinases, allowing signal amplification as well as positive and negative feedback loops (*8, 9*). Signals are propagated through the pathways via sequential phosphorylation of activation loops (A-loops) that lead to structural rearrangements, shifting conformational equilibria from from inactive to active states (*2, 10*). In the case of the MAP kinases, a TxY motif on the A-loop is doubly phosphorylated by their upstream kinases, the MAP2Ks or MEKs, unusual dual-specificity kinases able to phosphorylate both threonine and tyrosine residues. While the MAPKs are promiscuous, the MAP2Ks are highly selective, with ERK1 and 2 being the only known substrates of MEK1 (*11*). What determines signal specificity among closely related MAPK pathways has been a longstanding question in the field. In the last twenty years, studies identified short linear motifs (Kinase Interaction Motifs (KIMs or D-motifs)) that ensure some specificity (*12*). The KIMs consist of 12-17 residues, are located in the disordered N-terminal region of MAP2Ks, and dock to an allosteric site, the common docking (CD) site, on the MAPK (*13*). The KIM prepares the MAPK A-loop for phosphorylation (*14*), while the MAP2K N-terminal linker region, located between the KIM and core kinase domain, guides the MAP2K to the correct docking sites, aligning the kinases for catalysis (*15*). Once activated, the MAPK moves to cellular compartments and the nucleus, where cell-specific responses are induced via the activation of further protein kinases and transcription factors (*5*).

There is a wealth of data on the individual components of these pathways (*16–20*); however, until recently, details of the binding interactions and mechanisms of phosphorylation have been lacking. Cryogenic electron microscopy (cryo-EM) is now revealing many of the interactions between components of the pathways, including scaffolds and modulators, leading to a better understanding of signal transmission (*21–24*). Our work on the MKK6-p38α branch of the pathway (*15*) provided the first glimpse of a MAPK (p38α) being activated by its MAP2K (MKK6). It demonstrated how specificity could be maintained between structurally similar signalling components, and showed that substrate recognition was distal from the A-loop, a feature recently also observed in the CDK-CAK complex (*25*). However, the limited resolution did not allow a full description of activation. Here, we have determined cryo-EM structures of the MAP2K MEK1 (MAP2K1) in complex with its substrate MAPK ERK2 (MAPK1) in three states, to overall resolutions between 2.99 Å and 3.63 Å. Integrating these data with X-ray crystallography, hydrogen-deuterium exchange mass spectrometry (HDX-MS), molecular dynamics (MD) simulations and small-angle X-ray scattering (SAXS), we demonstrate that the interaction between the kinases is transient, with a high dissociation constant, but that multiple sites of interaction drive specificity and catalysis. Substrate recognition leads to unfolding of the regulatory A-helix on the MAP2K linker, a hotspot for disease mutations (*26, 27*), allowing MEK1 to move to an active conformation, explaining the mechanism of gain of function in some cancers. The determination of a second MAP2K-MAPK structure from a different pathway now allows the structural basis of MAPK specificity to be described. Remarkably, specificity resides on multiple sites remote from the active site, frequently on variable loop regions, allowing specific interactions between structurally similar components. Our data reveal the molecular details of MAP kinase signalling and pave the way for potential new drug development in cancer therapy.

### Engineering a stable MEK1-ERK2 complex

The interactions between MAPK pathway components must be transient in order to maintain signal transmission, posing a challenge for structural studies. Despite this, AlphaFold3 (AF3) (*28*) predicts a complex between the kinases with high confidence (Fig. 1A and S1) but specific interactions remain unclear. In order to stabilise the complex, we employed the same strategy that we applied to the MKK6-p38α complex by using the KIM sequence from the *Toxoplasma gondii* effector protein GRA24 (*29, 30*). We found that the GRA24 KIM also binds ERK1 and 2 with high affinity (K_d_ = 0.31 μM for both), at least 100-fold higher than the MEK1 KIM peptide (Fig. S2A). To obtain high-resolution information on the binding mode, we determined the crystal structure of human ERK1 (MAPK3) bound to a peptide corresponding to the sequence of the GRA24KIM1 at 2.17 Å resolution (Fig. S2 and Table S1). Human ERK1 was used as we were unable to obtain crystals with ERK2 (88% similarity between the two proteins). The binding of the GRA24KIM1 induces the same conformational change as the binding of the KIM sequence from MEK2 on ERK2 (4h3q (*31*), RMSD between the Cα of the kinases 0.79 Å, Fig. S2C and D). Further, we introduced phosphomimetic mutations in the MEK1 A-loop (S218D and S222D) to produce a constitutively active MEK1 chimera with the GRA24 KIM, named MEK1^DD^GRA (Fig. 1B and Fig. S3). The addition of the GRA24 KIM increases residue numbering by 6; for clarity, we use the wild-type MEK1 numbering (uniprot ID Q02750) throughout the manuscript. The activity of MEK1^DD^GRA was confirmed via native ESI-QTOF and LC-MS/MS showing phosphorylation of the ERK2^WT^ A-loop residues (Fig. S4). To limit complex heterogeneity through two possible phosphorylation events, we further stabilised the complex by mutating the ERK2 A-loop threonine to a valine, leaving only the tyrosine available for phosphorylation (ERK2^T185V^, Fig. 1B). Finally, we aimed to trap the transition state conformation with ADP and the transition state analogue aluminium tetrafluoride (AlF_4_ ^−^ ) (*32*). We were also able to purify a complex with the native KIM, the MEK1^DD^-ERK2^T185V^ complex (Fig. S3). However, structural studies led to poor 2D class averages and we pursued the MEK1^DD^GRA-ERK2^T185V^ complex for further high-resolution studies (Fig. S3E).

**Fig. 1.**
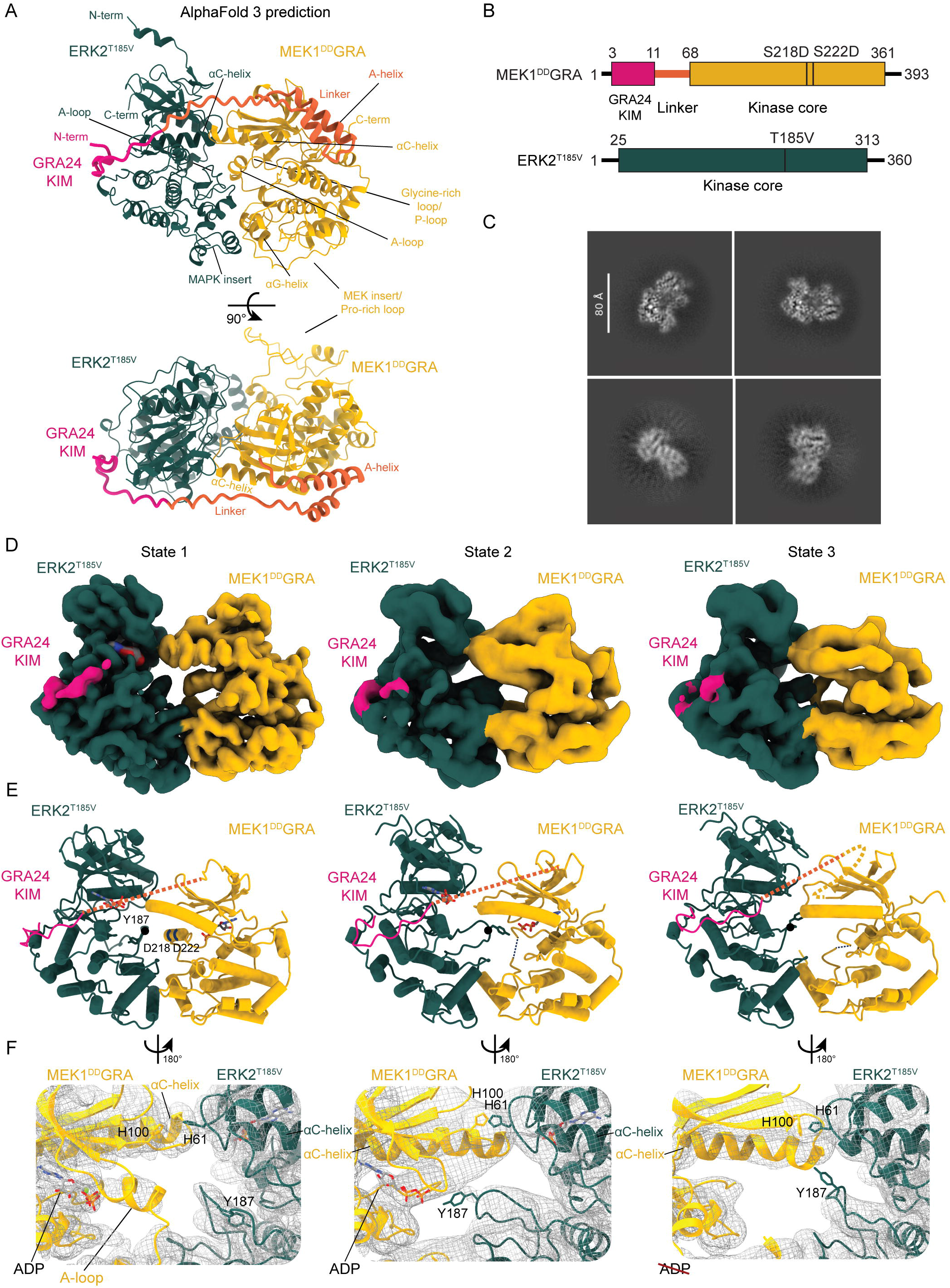
Structures of the MEK1^DD^GRA-ERK2^T185V^ complex. **A** AlphaFold 3 prediction of the MEK1^DD^GRA-ERK2^T185V^ complex. Important structural elements are labelled. The kinase core domain of MEK1 is coloured yellow, the ERK2 kinase core domain in dark green. The introduced GRA24 KIM-motif on MEK1 is coloured magenta, followed by the linker region (residues 12 to 67), in orange **B** Schematic representation of protein constructs. Numbers indicate domain limits and locations of mutants following Uniprot numbering, colour scheme as for A. **C** Representative 2D class averages from the dataset collected on HexAuFoil grids. **D** Cryo-EM reconstructions of MEK1^DD^GRA-ERK2^T185V^ complexes coloured according to A (state 1: inactive; state 2: active; state 3: nucleotide free). **E** Models of MEK1^DD^GRA-ERK2^T185V^ structures, with the flexible linker indicated by a dashed line. Side chains are displayed on residues of interest and labeled: The A-loop residue Y187 on ERK2 is the phosphate acceptor. D218 and D222 mutant sites render MEK1^DD^GRA constitutively active. **F** The LocScale sharpened Coulomb potential maps (grey mesh) showing the differences in the interface regions between the kinases. H100 on MEK1 and H61 on ERK2^T185V^ were identified as a novel site of contact.

We imaged the 86 kDa MEK1^DD^GRA-ERK2^T185V^ complex using cryo-EM. Initial micrographs recorded on UltraAuFoil grids showed promising results and reasonable class averages, but use of HexAuFoil grids (*33*) yielded improved two-dimensional (2D) class averages with clear secondary-structure features and low background (Fig. 1C), allowing the reconstruction of three different states of the complex from a single data set of 88,517 movies. Multiple rounds of picking and training using Topaz (*34*) were necessary to generate a stack of good-quality particles. A combination of 3D classification and heterogeneous *ab initio* reconstruction (*35*) including high-resolution information (HR-HAIR (*36*)) improved particle alignments and allowed the separation of three distinct states (Fig. 1D, and Fig. S5, S6). The most abundant state, resolved to an average resolution of 2.99 Å, is a non-productive complex where the A-loop of MEK1 is in the inactive conformation, and the A-loop of ERK2 cannot access the active site. In a second state, resolved to an average resolution of 3.47 Å, the A-loop of MEK1 is disordered, and the A-loop of ERK2 extends into the active site of MEK1 with the tyrosine (Y187 of the TxY motif) approaching the catalytic aspartate (MEK1 D190) and nucleotide. In the final state, resolved to an average resolution of 3.63 Å, the kinases are still complexed together, but no nucleotide is bound in MEK1, and the A-loop of ERK2 retracts from the active site with the tyrosine facing away (Fig. 1E and F, and S7). In all states ERK2 remains in a similar conformation with the only differences observed in the A-loop (Fig. S8A).

### Inactive MEK1 can bind its substrate with face-to-face architecture at multiple sites

As the inactive complex had the highest overall resolution, with the maps showing details of side chains, we used this state (state 1, inactive) to build the initial model and compare it to the subsequent states (Fig. 1D-F). The kinase domains are clearly resolved and are in very similar positions to the AF3 prediction (overall RMSD 2.0 Å), that also predicts the inactive conformation of the MEK1 A-loop (Fig S8B and E). Neither the disordered MEK insert (also known as the proline-rich loop), nor the N-terminal linker, were visible in the reconstruction, despite the latter having two confidently predicted helices (Fig. 1 and S1) one of which, the A-helix, is often resolved in crystal structures of MEK1 and is thought to have a regulatory role by stabilising the inactive conformation (*37*). Surprisingly, the A-loop of MEK1 is in the inactive conformation, despite phosphomimetic mutations. In this inactive state, a short helix in the A-loop of MEK1 prevents the A-loop of ERK2 from approaching the nucleotide in the MEK1 active site and the MEK1 αC-helix is in the inactive “out” position, in a similar conformation to the crystal structure of inactive MEK1 (*17*) (Fig. S8B). The overall interactions of the complex are comparable to the MKK6-p38α complex (*15*): the KIM interacts at the CD-site in a similar manner to the isolated KIM peptide (Fig. S9), and the αG-helix of MEK1 binds through mainly hydrophobic interactions, led by a MEK1 phenylalanine (F311), at the MAPK-specific insert of ERK2 (Fig. 2A). In the MEK1-ERK2 complex, additional interactions were identified: two histidine residues on the N-terminal lobes of the kinases, H100 on MEK1 and H61 on ERK2, and a triad of histidines coordinating an asparagine, MEK1 H239 and ERK2 H232 and H239, coordinating ERK2 D235 (Fig. 2A).

**Fig. 2.**
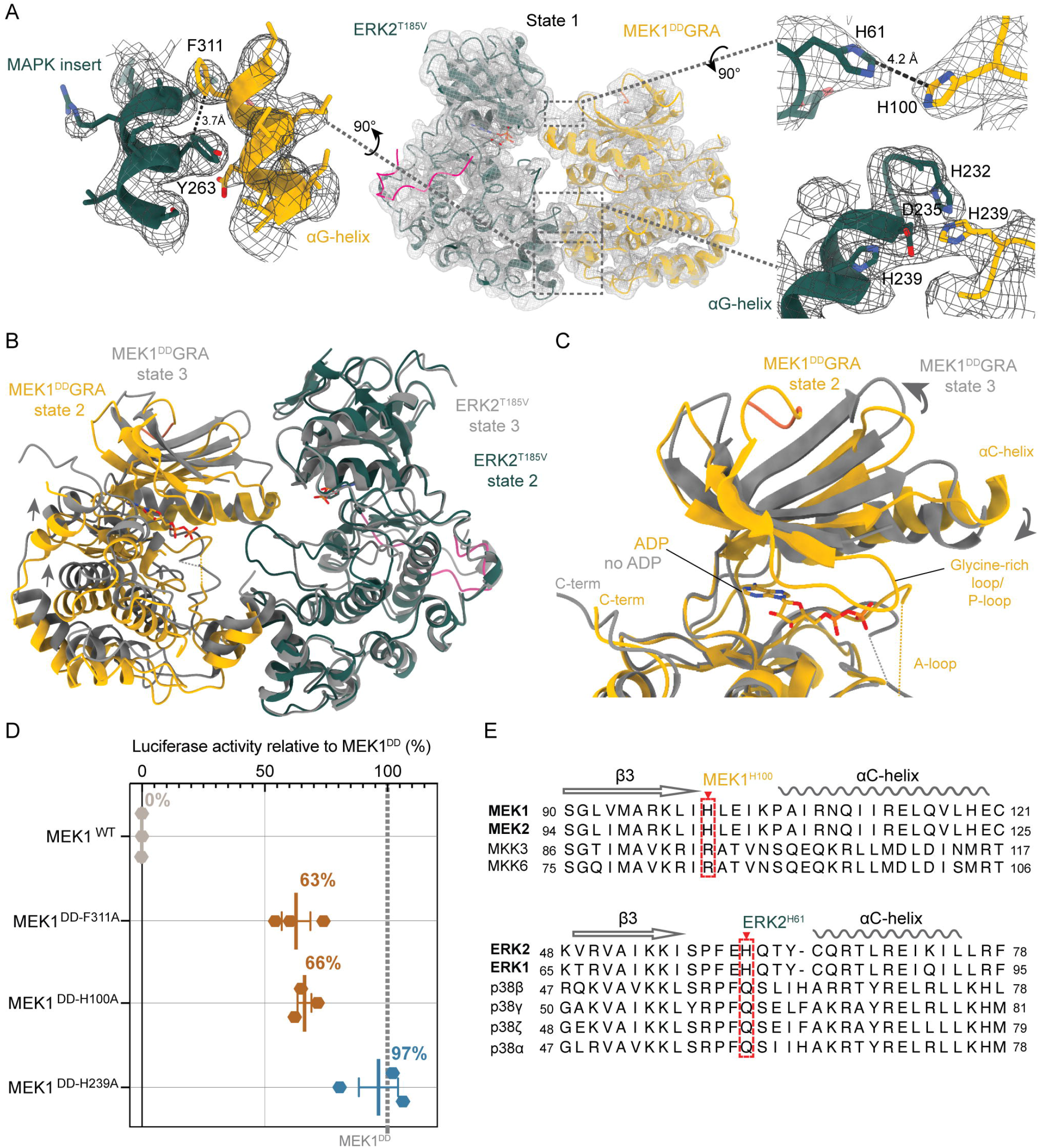
Interaction sites in the MEK1^DD^GRA-ERK2^T185V^ complex. **A** Beyond the KIM, MEK1 interacts with ERK2 at the hydrophobic pocket formed by the MAPK specific insert via the αG-helix (zoom on the left), a histidine triad coordinating a glutamate between the αG-helix of ERK2 and the loop between the αF-helix and MEK1 A-loop, and a permissive His-His interaction in the N-lobes (zooms on the right). Sharpened Coulomb potential maps from the State 1 reconstruction are displayed as a grey mesh with the state 1 model. **B** State 2 and 3 aligned on the substrate ERK2. Nucleotide free MEK1 of state 3 is rotated 5.6° and shifted up by 2 Å to slightly detach from its substrate. **C** The N-lobe of state 3 has opened 12.5° relative to the C-lobe of active MEK1, and the P-loop and β1-β2 are disordered, releasing nucleotide **D** Luciferase reporter assay of MEK1 mutants presented as activity relative to MEK1^DD^. MEK1^WT^ displays no activity in absence of stimulus, while mutations at contact sites show varying effects on signalling activity (mutant details in Fig. S10, RL and relative activity in Table S3, and S4, respectively) **E** The histidines are conserved in the MEK-ERK branch, but not the other MAPK pathways, and are conserved in metazoans (Fig S12).

### The active state binds its substrate at multiple sites distal from the A-loop

A second resolved state of the complex showed features for the A-loop of ERK2 entering the active site of MEK1 and no signal for the A-loop of MEK1 (Fig. 1F and S7A). This state (state 2, active) was determined at an overall resolution of 3.47 Å, but the main features of the kinases are well resolved. The relative orientation of the two kinases is the same as the state 1 complex, maintaining the same sites of contact, including the N-lobe histidine interactions, despite the movement of the MEK1 αC-helix and A-loop (Fig. S7B). In an active kinase conformation, once the αC-helix moves to the “in” position a conserved glutamate residue (MEK1-E114) forms a salt bridge with a lysine from the β3 strand (MEK1-K97), positioning it to coordinate the phosphates of the nucleotide, and allowing the DFG motif to move in to align a magnesium ion, thereby providing the correct environment for phosphoryl transfer. In state 2 the catalytic MEK1 αC-helix has moved towards the “in” conformation, when compared to the crystal structure of the inactive MEK1 (Fig. S8F), and the A-loop of ERK2 extends towards the active site of MEK1. The map in the region of the MEK1 αC-helix is blurred, implying movement (Fig S7C), and there is no clear position for MEK1 K97, making it difficult to see if the catalytic spine is in place. We were unable to further separate a class of particles with a better-defined position. A predicted model of active MEK1 (AF-Q02750-K1A, available from KinCore (https://dunbrack.fccc.edu/kincore/activemodels), (*38*)), including the K97 salt bridge, is not inconsistent with our maps (Fig. S7C), and our data suggest that the MEK1 αC-helix is likely in a continuum of positions. Refinement of the model placed the MEK1 αC-helix in an intermediate position that is consistent with the data and reflects an average position of the MEK1 αC-helix. We refer to this state as “active”, as the maps are consistent with the active conformation of the MEK1 αC-helix even if we cannot definitively place all the classical markers of an active kinase conformation. Although the complex was supplemented with ADP.AlF ^−^, we do not observe the metal fluoride, possibly due to the dynamic nature of the state. In our model, the A-loop tyrosine (Y187) of ERK2 is 6.2 Å from the MEK1 catalytic aspartate (D190) and 4.7 Å from the ADP β-phosphate. As with the MKK6-p38α complex, no contact between the MAPK A-loop and the MAP2K is observed, further supporting our hypothesis that substrate recognition occurs at sites remote from the active site and the substrate (ERK2) A-loop is relatively free to move within the confines of the MAP2K active site. This would allow either the tyrosine or threonine to access the active site for nucleophilic attack of the γ-phosphate.

### The nucleotide free state suggests a route for a processive mechanism

The third state, where nucleotide is absent from MEK1, was determined at an average resolution of 3.63 Å (state 3, nucleotide free). The same contact sites are maintained as in the other states and the MEK1 A-loop is disordered (Fig. 1F); however, MEK1 has rotated 5° relative to ERK2, but still maintains the N-lobe connection, and the MEK1 αG-helix moves away from the MAPK insert (Fig. 2B, C) but remains in the hydrophobic pocket. The most significant difference is the absence of nucleotide, along with complete disorder of the MEK1 P-loop (or glycine-rich loop) and the retraction of the ERK2 A-loop (Fig. 2B). The P-loop has an important role in coordinating the phosphates of ATP in the active site and positioning them for catalysis (*39*). In this state, the loop itself, and most of the β1-β2 strands, are disordered, and the nucleotide binding site is empty. However, the histidine interactions observed in the previous states are maintained (Fig. 1 and S7D, E) and the overall conformation is close to the active form (Fig. S7F). The MEK1 K_d_ for ADP is 2 μM (*40*), well below the buffer condition of 250 μM, implying that this conformation has significantly reduced affinity for nucleotides. This structure shows that a MAP2K can remain bound to its MAPK substrate while the nucleotide pocket opens, consistent with a processive mechanism. In a processive mechanism phosphorylation of both A-loop sites occurs without kinase dissociation, as opposed to a distributive mechanism, where the kinases completely dissociate between subsequent phosphorylations (*41*).

### A complex network of sites ensures specificity and primes the kinases for catalysis

To characterise our structural observations, we measured the effect of mutations in MEK1 on ERK signalling in cells using a luciferase reporter assay (Fig. S10, 11). We transiently transfected HEK293T cells with MEK1^DD^ mutants and a plasmid bearing the firefly luciferase under the control of the Serum Response Element (SRE) promoter, that is activated through substrates of ERK2 such as ETS Like-1 protein (Elk1). Luminescence read-outs were normalised through the use of a constitutively active renilla luciferase and signalling activity calculated relative to MEK1^DD^. Mutation of the MEK1 phenylalanine (F311), that interacts with the ERK2 C-lobe, to alanine, or the MEK1 histidine (H100) in the N-lobe, reduces signalling by 37% and 34% respectively, consistent with a role for these contacts in correctly positioning enzyme and substrate (Fig. 2D). Interestingly, mutation of MEK1-H239 to alanine did not significantly reduce signalling, although it is only one of many interactions in the C-lobe (Fig 2A and D). These histidines are conserved in MEK1 and 2 and ERK1 and 2 but are absent in the other branches of the MAPK cascades (Fig. 2E and S12).

We then used HDX-MS analysis of the MEK1^DD^-ERK2^WT^ complex (Fig. S3), containing the native MEK1 KIM motif and ADP.AlF_4_ ^−^, to validate interactions observed in the cryo-EM structure with a non-native KIM (Figs. 3A-D, S13) (*42*). Several regions of either MEK1^DD^ or ERK2^WT^ were protected from hydrogen-deuterium exchange when in each other’s presence, correlating with the cryo-EM models of the MEK1^DD^GRA-ERK2^T185V^ complex. Protected regions correspond to the MEK1 αC-helix and the β3-αC connecting loop -including MEK1 H100, that interacts with ERK2 H61 (Fig. 2A and 3A, B), and the MAPK insert of ERK2, where the αG-helix of MEK1 binds (Fig. 3C, D). This confirms that the interactions observed in our stabilised complex are the same in the presence of the native KIM.

**Fig. 3.**
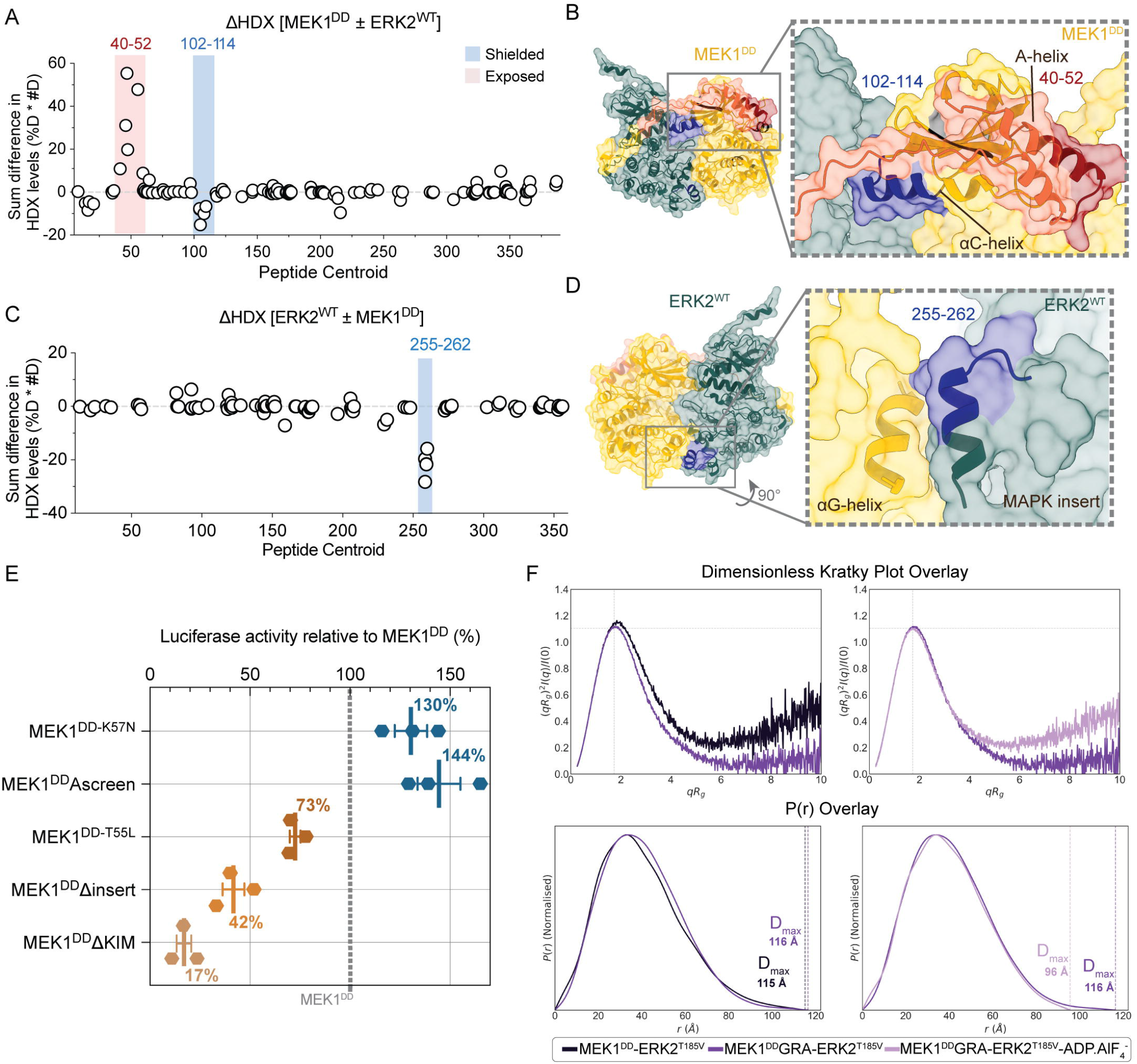
Dynamics of the MEK1-ERK2 complex. **A** Differences in deuteration levels (ΔHDX) for MEK1^DD^ alone in solution or in the presence of a 1.5 fold excess of ERK2^WT^. The x-axis corresponds to residue numbering from the N- to C-terminus of MEK1. Each dot represents a peptide, plotted according to its central residue (peptide centroid), calculated as a sum of differences for all measured time points (3s, 30s, 300s). The y-axis presents the product of the sum in differences, calculated both as percentage deuteration (%D) and number of deuterons (#D) to increase signal-to-noise ratio. Regions highlighted in blue indicate protection from exchange upon addition of ERK2^WT^, whereas regions highlighted in red show increases in H/D exchange rate, suggesting conformational change. Peptide map and deuteration level plot are shown in Fig. S13A and B and relevant uptake plots in Figure S13C. **B** Mapping of MEK1^DD^ regions undergoing change upon ERK2^WT^ binding identified in HDX-MS measurement onto a model of the MEK1^DD^-ERK2^WT^ complex displayed with surface and cartoon representation. The colouring is as Fig. 1A but including blue for shielded and red for exposed areas. **C** ΔHDX for ERK2^WT^ alone in solution or in presence of a 1.5 fold excess of MEK1^DD^. Graph presentation and colouring as for A. Peptide map and deuteration level plot in Fig. S13A and B. Uptake plot for shielded ERK2 peptide shown in Figure S13D. **D** Mapping of ERK2^WT^ site undergoing change upon MEK1^DD^ binding identified in HDX-MS measurement onto a model of the MEK1^DD^-ERK2^WT^ complex displayed with surface and cartoon representation. The colour scheme is as before but including blue for the shielded area. **E** Luciferase reporter assay of MEK1 mutants presented as activity relative to MEK1^DD^. Mutations in the A-helix show varying effects on signalling activity (mutant details in Fig. S10, relative luminescence (RL) and relative activity in Table S3, and S4, respectively). Removal of the MEK insert or KIM leads to signal reduction. **F** Top row: Dimensionless Kratky plot from SEC-SAXS experiments for MEK1^DD^-ERK2^T185V^ (dark purple) and MEK1^DD^GRA-ERK2^T185V^ (purple) complexes overlaid for comparison, the introduction of the GRA KIM yields a more globular particle with reduced flexibility. The second dimensionless Kratky plot shows overlay of MEK1^DD^GRA-ERK2^T185V^ (purple) and MEK1^DD^GRA-ERK2^T185V^-ADP.AlF ^-^ (light purple); the addition of nucleotide and transition state analogue further compacts the complex but increases local flexibility. The bottom row shows P(r) distribution for MEK1^DD^-ERK2^T185V^ (dark purple) and MEK1^DD^GRA-ERK2^T185V^ (purple) which have similar d_max_ values. In the second panel, the P(r) distribution for MEK1^DD^GRA-ERK2^T185V^ (purple) and MEK1^DD^GRA-ERK2^T185V^-ADP.AlF_4_^-^ (light purple) are seen, where a lower D_max_ for sample containing ADP.AlF_4_^-^ addition indicates complex compaction.

In addition to the areas showing protection, the A-helix in the N-terminal linker of MEK1 showed significant deprotection upon binding of ERK2, suggesting an unfolding of the helix when bound to ERK2 (Fig. 3A and B). This helix has been assigned a regulatory or inhibitory role, and is thought to stabilise the MEK1 αC-helix in the inactive “out” position (*37*). The MEK1 A-helix is a hotspot for mutations found in cancer and other diseases. These mutations have been demonstrated to either destabilise the A-helix, leading to partial unfolding and higher P-loop flexibility (*43, 44*) or remove interactions with the MEK1 kinase core that stabilise the inactive conformation (*37, 44*). We returned to the luciferase reporter assay in order to test the connection of the A-helix with substrate recognition and catalytic activity (Fig. S10E). One of the most common cancer mutations found in MEK1 is K57N (*45*). The removal of the MEK1 A-helix lysine breaks an interaction with the MEK1 αC-helix that stabilises the inactive conformation (*37*). We tested MEK1^DD-K57N^ as a control for increased signalling and observed an increase of 130% relative to MEK1^DD^ (Fig. 3E). We then tested two constructs, one designed to disrupt the helix and all interactions between the A-helix and the kinase core by replacing the A-helix (residues 49 to 60) with alanines (MEK1^DD^Ascreen); and another mutant designed to increase the stability of the A-helix (MEK1^DD-T55L^). Using well established helical propensity properties of amino acids (*46*) we examined the residues on the solvent exposed surface of the A-helix with no interaction with the kinase core (Fig. S10E). We selected T55L as a point mutation likely to increase the helical propensity of the A-helix, replacing a residue with very weak helix properties with one of the strongest, while also increasing hydrophobic packing of the A-helix and maintaining minimal impact on overall protein interactions. Disrupting the A-helix (MEK1^DD^Ascreen) led to an increase in signalling of 144%, whereas stabilising the A-helix (MEK1^DD-T55L^) reduced signalling by 27% compared to MEK1^DD^ (Fig. 3E). These data support the premise that the A-helix unfolds in response to correct substrate binding, leading to greater conformational freedom, ultimately affecting activity.

The proline-rich loop, specific to MEK1 and 2, shows high flexibility in the HDX-MS data (Fig. S13). The insert has been implicated in binding to scaffolds and upstream components of the pathway, and enhances ERK activation (*47*). We observed residual features in the maps of three states of the MEK1^DD^GRA-ERK2^T185V^ complex near the ERK2 A-loop (Fig. S14). The signal is within reach of the proline-rich loop; however, it is not sufficiently clear to place any residues. Despite the indications of an interaction with ERK2 in the cryo-EM reconstructions, no increased protection of this loop was observed in the HDX-MS data. It should be noted that the KIM sequence of MEK1 does not show increased protection upon addition of ERK2 either, even though this is a well-characterised and defined binding site, highlighting the highly dynamic nature of the interactions. Upon inspection of the uptake plots for the KIM peptides, a trend of protection can be observed (Fig. S13C), but remains not significant in triplicate, likely also due to the very fast exchange rate already at 3s. We therefore tested a mutant of MEK1^DD^ with the proline rich loop removed (MEK1^DD^Δinsert) in the luciferase reporter assay. Removal of the loop leads to a 58% reduction in signalling relative to MEK1^DD^ (Fig. 3E). We also tested a construct with the N-terminal KIM removed (MEK1^DD^ΔKIM) and found signalling was reduced by 83% (Fig. 3E) demonstrating the importance of the KIM in initial substrate recruitment. These data imply that a transient interaction between the proline rich loop and ERK2 is required for recognition and catalysis and is a further specificity checkpoint unique to the MEK-ERK branch of the MAP kinases.

### Solution studies of the MEK1^DD^GRA-ERK2^T185V^ and MEK1^DD^-ERK2^WT^ complexes demonstrate a highly dynamic interchange between the kinases

We used isothermal titration calorimetry (ITC) to measure the interaction between the MEK1^DD^GRA-ERK2^WT^, MEK1^DD^-ERK2^WT^ and MEK1^WT^-ERK2^WT^ kinases. While we were able to determine dissociation constants for MEK1^DD^GRA and ERK2^WT^ (1.94 μM) we were unable to measure dissociation constants between either MEK1^DD^ and ERK2^WT^ or MEK1^WT^ and ERK2^WT^ (Fig. S15 and Table S5), implying that the K_d_s are higher than 10 μM, consistent with the value of 29 μM determined using fluorescent labeled proteins in cells (*48*). This is compatible with the requirement of the kinases to transmit signals via phosphorylation, but not monopolise either the MAPK or MAP2K, so they are free to continue to activate targets. Despite this low affinity, we were able to purify the MEK1^DD^-ERK2^WT^ and MEK1^DD^-ERK2^T185V^ complexes at a starting concentration of 50 μM, where they eluted in a single peak (Fig. S3) during SEC. This is in contrast to our studies of the MKK6-p38α complex, where we were unable to purify the native complex. The ability to purify the native complex opened the possibility to study the complexes by SAXS.

We studied both the MEK1^DD^GRA-ERK2^T185V^ and MEK1^DD^-ERK2^T185V^ complexes by size-exclusion chromatography-coupled small-angle X-ray scattering (SEC-SAXS) in the absence of nucleotide and in the ADP.AlF_4_^−^ supplemented state (Fig. S16 to S18 and Table S6). Analysis of the elution traces revealed conformational heterogeneity in the non-stabilised complex (MEK1^DD^-ERK2^T185V^). While the molecular weight (MW) estimates across the peak remain relatively consistent with a 1:1 heterodimer, the Radius of Gyration (R_g_) exhibited a continuous decrease (34 Å to 28 Å), indicative of a polydisperse ensemble of extended and compact conformations resolving on the column (Fig. S16). Introduction of the GRA KIM (MEK1^DD^GRA-ERK2^T185V^) stabilises the complex: the R_g_ across the elution peak is constant (c. 32 Å), indicative of a monodisperse sample with uniform shape. This was further supported by the dimensionless Kratky plot, which showed a shift in peak position and curve shape reflective of a more compact particle for MEK1^DD^GRA-ERK2^T185V^ when compared to MEK1^DD^-ERK2^T185V^ (Fig. 3E and S18).

To assess the agreement between the cryo-EM structure and the conformation in solution, we generated theoretical scattering profiles from the composite model of the inactive state using PEPSI-SAXS (*49*). This yielded reasonable goodness-of-fit values (χ^2^) to the experimental data of 2.70 for MEK1^DD^GRA-ERK2^T185V^ and 3.61 for MEK1^DD^GRA-ERK2^T185V^-ADP.AlF ^−^ after refinement of the model using non-linear normal mode analysis. The dimensionless Kratky plot of the ligand-bound species showed a characteristic upward deviation at higher qR_g_ values (qR_g_ > 6) compared to the apo form, reflective of increased local flexibility, while the pair distance distribution function, *P*(r), shows a smaller D_max_ for the ligand-bound form (Fig. 3F). The formation of the active conformation of the MEK1^DD^GRA-ERK2^T185V^ complex shows a global compaction of the core, but an increase in local flexibility as well. Collectively, this suggests that in presence of ADP.AlF ^−^ the complex locks into a compact architecture, while simultaneously inducing local plasticity or elongation of linker or loop segments (Fig. 3F). This heterogeneity in the catalytically competent heterodimer is reflected in the cryo-EM dataset, particularly in states 2 and 3, suggesting multiple conformations in the active site, allowing fast turnover of nucleotide and substrate.

### Molecular dynamics simulations show that competent catalytic states can be formed in the active state and highlight the dynamic nature of the interaction

To investigate the dynamics of the complex, and integrate the observations collected from our structural and solution data, we also performed MD simulations starting from a set of structures derived from the cryo-EM models. For the state 1 and 2 models the following modifications were made: First we reverted the A-loop of ERK2 to the wild type and combined the model with AF3 predictions to produce a composite model of MEK1^DD^GRA-ERK2^WT^ (Fig. 4A). Additionally, we produced a third model with the catalytic salt bridge in MEK1 formed, that is consistent with the maps of state 2. For each of these three starting MEK1^DD^GRA-ERK2^WT^ states, we also created models where the KIM of MEK1 was reverted to the wild type sequence to produce a composite model of the MEK1^DD^-ERK2^WT^ complex (Fig. 4B). We then ran MD simulations of these six models to explore active and inactive states of both MEK1^DD^-ERK2^WT^ and MEK1^DD^GRA-ERK2^WT^ complexes, for a set of 48 replicas (8 per system) of 2 μs each (Table S7).

**Fig 4.**
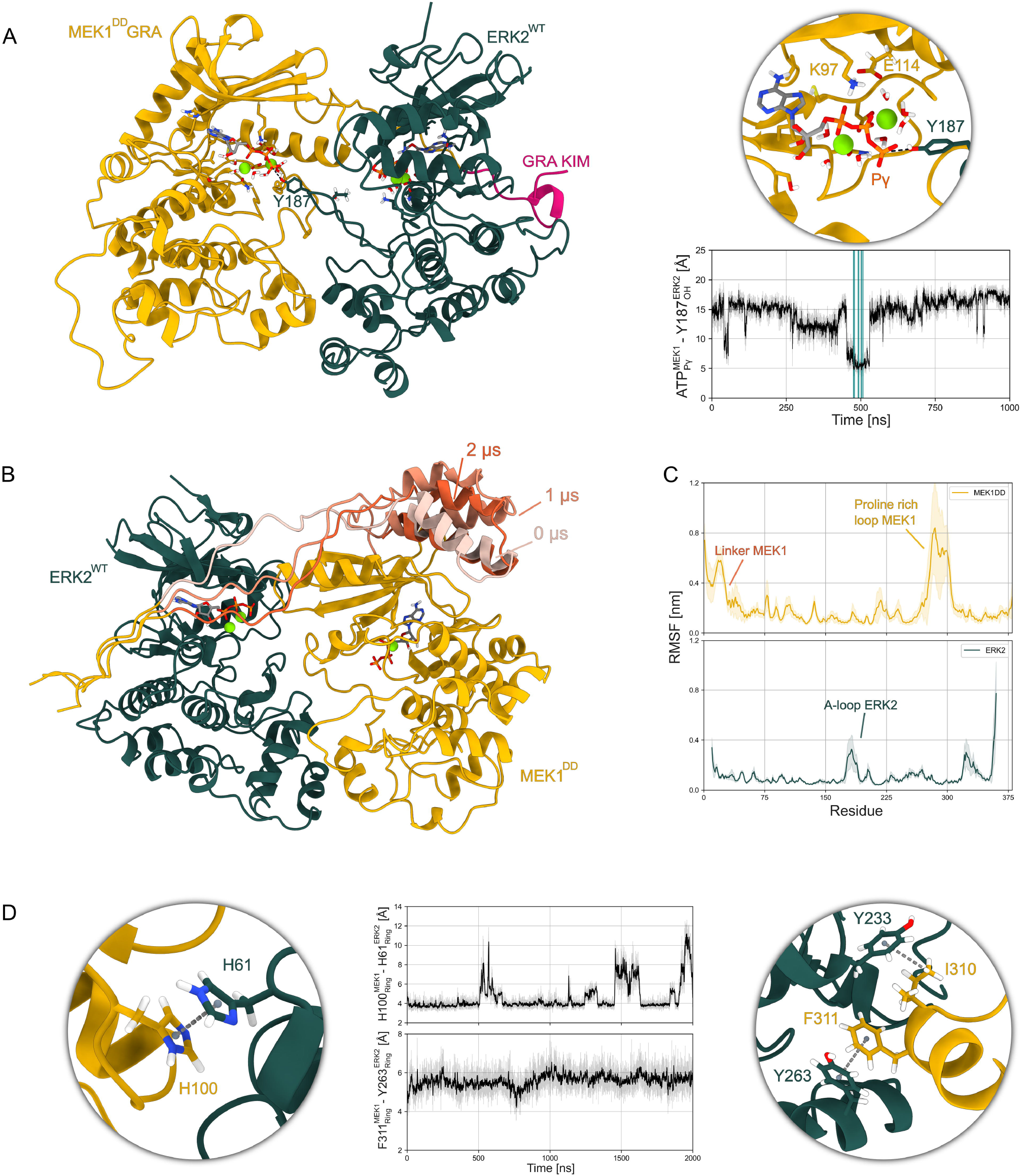
MD simulations show that ERK2 Y187 can reach MEK1 ATP and that the flexibility of the system is consistent with HDX-MS and cryo-EM data. A Frame extracted from an MD simulation (see Movie S1) of the fully active state in which Y187 of ERK2^WT^ approaches the γ-phosphate of MEK1^DD^GRA ATP at a catalytically compatible distance (∼3.8 Å). The inset shows K97 interacting with the ATP α-phosphate as well as with E114 to form the salt bridge that stabilizes the αC-in active state. In the distances over time plot, the frames where ERK2^WT^ Y187 reaches MEK1^DD^GRA ATP at a catalytically compatible distance are highlighted in blue. B Three frames extracted from an MD simulation of the inactive state of the MEK1^DD^-ERK2^WT^ at different times where the α-helices of the N-terminal linker of MEK1^DD^ show large rearrangements. C Average root mean square fluctuation (RMSF) for all the MEK1^DD^GRA-ERK2^WT^ MD simulations (24 replicas, 48 μs). D Distances between MEK1^DD^GRA H100 and ERK2^WT^ H61 at the N-lobe interface and between the MEK1^DD^GRA F311 and ERK2^WT^ Y263 at the MAPK-specific insert αG-helix interface. The insets show frames extracted from the corresponding time series.

The simulations explored several states of the active and inactive conformational ensembles. The synthetic HDX-MS profiles computed from these simulations show reasonable agreement with the experimental ones (Fig. S19), suggesting that the ensembles explored capture key features of the dynamic behavior observed in solution. Occasionally, the simulations of the active complexes explored catalytically competent states in which Y187 of the ERK2 A-loop approached the γ-phosphate of the ATP in the active site of MEK1 at distances compatible with phosphate transfer (Fig. 4A). The formation of this state seems to be driven by both an approach of the ERK2 A-loop to the active site of MEK1 and an outward rotation of Y187 (Movie S1). Interestingly, in no simulation did T185, the other phosphorylation site of ERK2, approach the γ-phosphate of ATP at the timescales investigated.

The analysis of the flexibility of the complex revealed characteristics consistent with the cryo-EM and HDX-MS data (Fig. 4B, C and S19 and 20). In ERK2, the most flexible domain is the A-loop, even in the inactive state (Fig.4D and S21), demonstrating that in the crowded environment the kinase still retains partial freedom of movement. In MEK1 the most flexible domains are the A-helix / N-terminal linker region and the proline-rich loop, the latter exhibiting considerable conformational freedom (Fig. S20). Remarkably, in some simulations the A-helix of the N-terminal linker of MEK1 detached and showed increased flexibility (Fig. 4C), consistent with both the absence of this domain in the cryo-EM maps and its increased deuterium exposure in the HDX-MS measurements on binding ERK2.

In all of the simulations, the kinases stayed at least partially in contact and did not fully dissociate. As for the MKK6-p38α complex, the C-lobe proved to be the critical interface for the dimer stability (Fig. 4D and S22). The N-lobe interface, however, is much more ephemeral, and can be lost and re-formed while the complex is held together by the C-lobe interface and the linker (Fig. S23). In a number of simulations MEK1 H100 and ERK2 H61 were close at the N-lobe interface (Fig. 4D), supporting the experimental findings that this interaction contributes to substrate specificity but is not so tight as to prevent substrate release.

Overall, the two coordinating magnesium ions in each of the binding sites of the active complexes show high mobility (Fig. S24). While the ions do not leave the binding site and remain coordinated to the ATP phosphates, in most of the simulations at least one of the two Mg atoms leaves the proximal oxygen of the aspartate of the DFG motif. This mobility is in line with the difficulty to resolve the ion positions in the cryo-EM maps, and might suggest that the second coordinating metal ion is only transiently present in the binding site at the moment of phosphorylation. Conversely, the single coordinating ion in all the inactive simulations (totalling 32 μs) is extremely stable (Fig. S24). In this configuration, the partially folded A-loop of MEK1 limits the available space for the aspartate of the DFG domain, hinting that the inactive loop might hinder down-stream phosphorylation both by obstructing the access to the binding site and by limiting the aspartate freedom to host a second ion at the binding site. These simulations confirm the binding sites observed in the cryo-EM structures and highlight the overall flexibility of the complex, particularly the active site of MEK1, that is in constant turnover even in the active, substrate bound form.

## Discussion

We have determined snapshots of various stages of the activation of the MAPK ERK2 by its upstream MAP2K MEK1. Combining these structures with biophysical studies in solution, as well as molecular dynamics simulations, we provide insight into the mechanisms of substrate recognition and catalysis in this important signalling pathway.

Our study has revealed a complex network of interactions between the MAP2K and its substrate MAPK. A surprising feature is the ability of MEK1 to bind its substrate while being in the inactive conformation (state 1). In solution, the MEK1 A-loop is probably in equilibrium between active and inactive states, with phosphorylation driving it to the active side (*23*). The equilibrium between conformations would also explain why the wild type MAP2Ks exhibit basal activity (*50*). The aspartate mutations applied in this study appear not to be as effective as phosphorylation, biasing the equilibrium enough for enzymatic activity but not completely to the structurally active state. In crystal structures of MAP2Ks with the same mutations, the A-loop is not visible (*16, 18*), implying high flexibility, averaging the A-loop active and inactive conformations that we were able to partially separate computationally within our cryo-EM data. The use of the constitutively active mutations replacing the serines of MAP2K A-loops with either aspartate or glutamate residues clearly biases the A-loop towards the active conformation, but is probably an imperfect mimic of the phosphorylated conformation.

Long-timescale MD and experimentally validated adaptive Markov state model simulations have suggested that MAP2K-MAPK complex disassembly proceeds sequentially, with the N-lobes dissociating first, followed by the αG-helix and finally the KIM (*15*). The observed dissociation hierarchy implies a reverse assembly pathway in which KIM docking precedes αG-helix engagement and, finally, the formation of weaker N-lobe contacts, assuming that the association proceeds along the reverse of the dominant dissociation pathway. While alternative association pathways are possible, this interpretation is consistent with the MD and in line with the main association and dissociation pathway observed for the MKK6-p38α complex. This is additionally supported by the observation that there is almost no catalytic activity if the KIM is removed (Fig. 3E). The N-terminal kinase interaction motif (KIM) on the MAP2K is therefore likely responsible for initial docking and preparation of the MAPK for phosphorylation by exposing the A-loop (*14*). However, it also transmits instructions to the MAP2K via its linker. We previously demonstrated that the linker between the KIM and kinase core is an important feature of specificity (*15*). We hypothesised that the length was important in guiding the αG-helix to the MAPK insert. We find that in MEK1, the helical elements in the linker play an additional role in recognising substrate binding. The A-helix acts as a hand break, only released by unfolding on substrate binding, thereby freeing the αC-helix, allowing the catalytically competent conformation to be formed. This helix is the site of many disease linked mutations (Fig. S25A) that also lead to helix unfolding and P-loop destabilisation (*44*), likely causing aberrant signalling by releasing the break on the αC-helix normally linked to substrate binding. Our assay data show that by stiffening the helix signalling is reduced, consistent with locking the break on the αC-helix. The observation of A-helix unfolding on substrate binding (Fig 3 and B), combined with our structural and assay data (Fig 3E), suggests a mechanism for the gain-of-function of mutations in this region: These mutants would be primed for activity before substrate engagement, and may have increased flexibility in the P-loop region allowing faster nucleotide exchange, leading to higher activity, or indeed increased activity in the absence of A-loop activation by phosphorylation, causing aberrant signalling.

Once the KIM and αG-helix are engaged, the N-lobe interaction via MEK1-H100 and ERK2-H60, position the two kinases correctly for catalysis. Histidines are advantageous for transient interactions as they are geometrically permissive and slightly weaker than classical salt bridges (*51, 52*) as well as being able to respond to pH differences (*53*) that could have implications for signalling in different sub-cellular compartments. The identification of a specific interaction between residues on hyper-variable loop regions suggests a potential common site for kinase-kinase interactions in addition to the inserts, and extensions to the N and C termini, common to many protein kinases. These variable loops are free from the constraints of the kinase fold and can therefore shift according to pathway requirements (Fig. 2E and S12). It is interesting to note that the interface with MEK1 and its substrate ERK2 is quite different to the MEK1 activating kinase CRAF (Fig. S25B) demonstrating the number of different specific interfaces that must be maintained. An additional check on substrate recognition is the proline-rich loop - specific to MEK1 and 2 - that likely engages with the ERK2 A-loop (Fig. 3E and S14). A threonine residue in this loop (T292) is also a site of negative feedback phosphorylation (*54, 55*). This could prevent the interaction with ERK, regulating signalling. This area is therefore a potential additional region of interaction between the kinases, but like the other sites, it is low-affinity and transient, contributing to overall complex formation and specificity. At this stage, even when all the specific interactions are fulfilled, if the MEK1 A-loop remains in the inactive conformation, it would not be able to activate ERK2, as the short helix blocks access to the nucleotide (Fig. 1F). If the MEK1 A-loop is in the active conformation, the A-loop of ERK2 can approach the catalytic site of MEK1 with a large amount of freedom. Our structure of an active conformation (state 2) shows that there is no interaction between the MAPK A-loop and the MAP2K beyond the nucleophilic oxygen atom approaching the ATP. Our solution and MD simulation data demonstrate considerable dynamics in the active site and loops of the complex when bound to each other, implying that the site is in constant turnover between aligned and unaligned active states (Fig. 3, 4, S21). This would allow the first phosphoryl transfer to occur while avoiding any low-energy wells that could stall the second step. Our structure of the nucleotide free state (state 3) demonstrates that the MAP2K can efficiently exchange nucleotides while being engaged with its substrate, as the P-loop is disordered, allowing ADP release. It has long been debated whether phosphorylation of the MAPKs by their MAP2Ks occurs via a distributive mechanism, where the MAP2K dissociates from the MAPK between phosphorylation events, or a processive mechanism, where nucleotide is released and rebound to allow the second phosphorylation, without dissociation of the kinases. There is strong experimental evidence for both mechanisms, with a dissociative mechanism being more likely *in vitro* (*50, 56*) and a processive mechanism becoming more likely in crowded conditions or in the cell (*57, 58*). In the state 3 structure the A-loop of the MAPK has sufficient conformational freedom so that a phosphorylated residue should not hamper the approach of the second residue to the active site. This shows how a processive mechanism could proceed as the complex does not restrict either A-loop flexibility or nucleotide exchange. While we should bear in mind that this is a stabilised complex, it shows a bound, nucleotide-free conformation compatible with nucleotide exchange, consistent with a processive mechanism. Together, these data demonstrate information relay between enzyme and substrate, remote from the active site, leading to efficient and highly specific activation, a hallmark of this signalling cascade.

The determination of a second MAP2K-MAPK complex structure allows the basis of pathway specificity to be described (Fig. 5A, B). Comparison of the MKK6-p38α complex with the MEK1-ERK2 complex shows similarities in global interactions, but significant differences that would prevent signalling crosstalk. Even though the KIM and αG-helix bind in a very similar manner, and the face-to-face orientation is conserved, the relative position of the enzyme to the substrate is significantly different (Fig. 5C). Superposition of the substrate MAPKs (p38α and ERK2) shows that MKK6 is rotated 36°, relative to MEK1, towards the MAPK substrate with a 5.7 translation along the axis, such that the MKK6 N-lobe is higher relative to its substrate than MEK1 (Fig. S25C). The rotation axis is centred around the point just after the KIM, at which the N-terminus of the MAP2K departs towards the N-terminal lobe, highlighting the linker length as an important feature for correct substrate engagement (*15*) (Fig. S25C). The N-terminal linker of MEK1 serves as another check on the substrate by releasing the catalytic machinery only upon correct substrate binding. The p38α branch does not contain helices in the N-terminal linker, but MEK1 and 2, as well as MKK7, do. Swapping the MKK6 linker for those from MEK1 and 2 abolishes p38α signalling in cell assays (*15*). Therefore, in addition to linker length, the regulatory helices also play an important role in substrate discrimination. Once the αG-helix is docked, and additional C-lobe interactions are in place, the N-lobe interactions must then be fulfilled. The major difference in the interfaces is between the N-lobes of the kinases: a His-His interaction (Fig. 2A), maintained only in the MEK-ERK branch, is the final site that likely determines specificity. If the substrate MAPK is incorrect, the N-lobe interaction will not be stabilised, preventing alignment of the MAPK A-loop with the MAP2K active site. The moderate resolution of the MKK6-p38α structure did not allow confident residue assignment in the N-lobe interaction, but residues in this area are also conserved in this branch, indicating another possible site determining specificity (Fig. 2E and 5). The MEK branch may also have the proline-rich loop as an additional site of interaction at the ERK A-loop (Fig. S14), adding a further verification site.

**Fig. 5.**
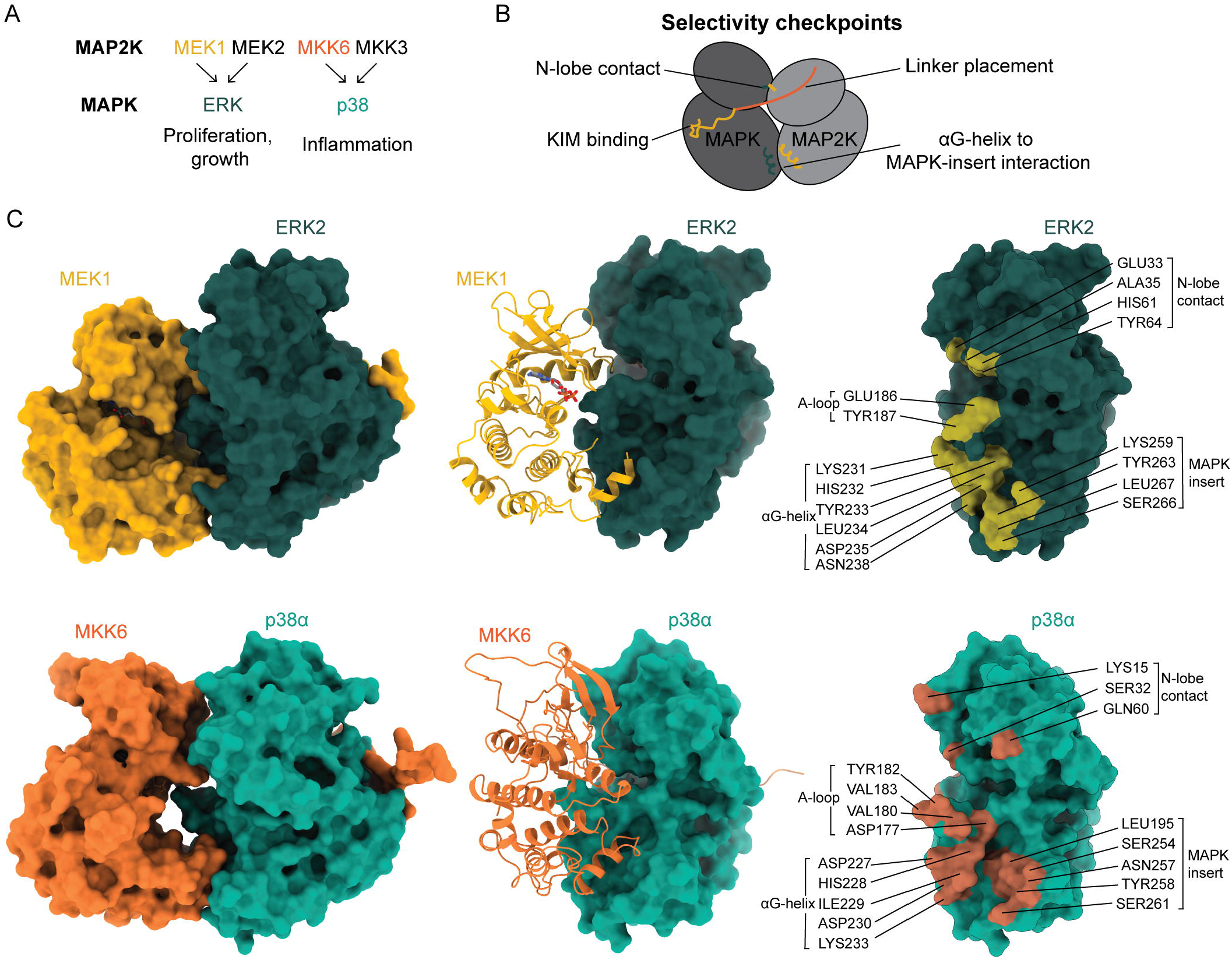
Molecular details of MAPK pathway specificity. **A** Schematic showing the specificity of MAP2Ks for MAPKs and the consequences of signalling nodes with components with structural data on the interaction coloured. **B** Diagram describing the selectivity checkpoints in the MAP2K-MAPK interactions of the MAP kinase pathways. **C** Comparison of the MEK1^DD^GRA-ERK2^T185V^ complex with the MKK6^DD^GRA-p38α^T180V^ complex. While the overall face-to-face conformation is maintained between pathways, significant differences exist in the relative positioning of the kinases (a 36° rotation of MKK6 relative to ERK2 leads to a higher N-lobe interaction) and the residues that interact. From left to right the complexes are displayed as space filling surfaces, the substrate MAPK as a surface and the MAP2K as a ribbon, and the surface of the substrate MAPK with the interface residues highlighted.

Our structural data delineate several snapshots in the activation of ERK2 by MEK1. As with the MKK6-p38α complex, substrate recognition is at sites remote from the active site, allowing considerable flexibility in the residues that can be phosphorylated while maintaining high specificity. In combination with structural studies in solution, and molecular dynamics, our data point to a complex that is highly dynamic during catalysis, allowing fast turnover, essential in signalling, and avoiding low energy states, with an efficient processive mechanism likely *in vivo* (*57*). Together, we describe the structural basis of ERK activation, a fundamental step in many cellular processes and a potential checkpoint for cancer and other diseases. While there are several drugs in clinical use that target the MEKs, ERK1 and 2 do not have drugs in clinical use. Our structural data could facilitate the development of new therapeutics.

## Supporting information

MD simulation trajectory of MEK1DDGRA-ERK2WT fully active complex as Y187 on the activation loop of ERK2 forms a catalytical competent state with ATP

HDX-MS data tables for MEK1DD

HDX-MS data tables for ERK2

## Acknowledgments

Cryo-EM data were collected at CM02, the purchase of this microscope was funded by the EquipEx+ France Cryo-EM project (ANR-21-ESRE-0046) and we thank Guy Schoehn (IBS, Grenoble) for generous support on the microscope and Felix Weis (IBS, Grenoble) for helpful discussions. We thank K. Djinovic-Carugo and A. McCarthy (EMBL Grenoble) for encouragement and M.-A. Hakimi (IAB, Grenoble) for introducing us to the *T. gondii* protein GRA24. We acknowledge the European Synchrotron Radiation Facility (ESRF) for the provision of synchrotron radiation facilities, and we would like to thank the staff of the ESRF and EMBL Grenoble for assistance and support in using beamlines BM29 and MASSIF-1 (ID30A-1). The Swiss National Supercomputing Centre (CSCS) is acknowledged for its generous allocation of supercomputer time on ALPS (project IDs: lp84 and lp66). We thank Martin Pelosse (EMBL Grenoble) for support in eukaryotic protein expression, Jennifer Schwarz at the proteomic core facility at EMBL Heidelberg for mass spectrometry, and Iskander Khusainov and Romain Linares for excellent support in using the EM Facility at EMBL Grenoble. We thank Rémy Visentin and Alexandre Hainard (University of Geneva) for assistance with HDX-MS data acquisition and analysis. NP thanks Yiannis Galdadas (IIT, Italy) and Eleonora Gianquinto (University of Turin, Italy) for useful discussions. This work used the platforms of the Grenoble Instruct-ERIC centre (ISBG ; UAR 3518 CNRS-CEA-UGA-EMBL) within the Grenoble Partnership for Structural Biology (PSB), supported by FRISBI (ANR-10-INBS-0005-02) and GRAL, financed within the University Grenoble Alpes graduate school (Ecoles Universitaires de Recherche) CBH-EUR-GS (ANR-17-EURE-0003). Molecular graphics and analyses were performed with UCSF ChimeraX, developed by the Resource for Biocomputing, Visualization, and Informatics at the University of California, San Francisco, with support from National Institutes of Health R01-GM129325 and the Office of Cyber Infrastructure and Computational Biology, National Institute of Allergy and Infectious Diseases.

## Author contributions

Conceptualization: JvV and MWB

Methodology: JvV, PJ, NP, HF, KL and OV

Investigation: JvV, PJ, NP, HF, KL, OV, FLG and MWB

Visualization: JvV, PJ, NP, HF, KL and OV

Funding acquisition: MWB, FLG

Supervision: MWB, FLG

Writing - original draft– JvV and MWB

Writing – review & editing: JvV, PJ, NP, HF, KL, OV, FLG and MWB

## Data and materials availability

Accession codes: Structure of the ERK1-GRA24KIM1 complex: https://doi.org/10.15151/esrf-dc-2310714581 (raw diffraction data); PDB-9TU0 (coordinates and structure factors; Protein Data Bank). Structures of the MEK1DDGRA-ERK2T185V complex: EMPIAR-13965 (micrographs and particle stack) and https://doi.esrf.fr/10.15151/ESRF-ES-2012234353 (raw micrographs), EMD-56418 (state 1), EMD-56419 (state 2) and EMD-56420 (state 3) (maps; Electron Microscopy Data Bank) and PDB-9TYG (state 1), PDB-9TYH (state 2) and PDB-9TYI (state 3) (coordinates; Protein Data Bank). The SAXS curves and pair distribution functions are deposited in the Small Angle Scattering Biological Data Bank (access codes: SASDY56, SASDY66, SASDY76 and SASDY86). The mass spectrometry proteomics and HDX-MS data have been deposited to the ProteomeXchange Consortium via the PRIDE partner repository with the dataset identifiers PXD073457 and PXD073252 respectively.

## Supplementary Materials

Materials and Methods Figs. S1 to S25

Tables S1 to S7 References (*59–111*) Data S1 and S2 Movie S1

