## Supplementary material for "Molecular basis of mitogen-activated protein kinase ERK2 activation by its upstream kinase MEK1": Materials and methods and Supplementary Figures

**The PDF file includes:**

Materials and Methods
Figs. S1 to S25
Tables S1 to S7
References 59-111

**Other Supplementary Materials for this manuscript include the following:**

Data S1 to S2 and Movie S1

### Materials and Methods

Plasmids

Plasmids were ordered from GenScript (gene synthesis, cloning and mutagenesis). For recombinant protein expression, human ERK2 (MAPK1; WT and the variant T185V) and ERK1 (MAPK3) sequences were fused to a His6 tag with a 3C protease cleavage site and cloned into a pET-28b(+) vector. The MEK1 constructs (MAP2K1; $MEK1^{WT}$, $MEK1^{DD}$ (constitutively active S218D S222D mutant of human MEK1), $MEK1^{DD}$GRA (constitutively active S218D T222D mutant of human MEK1 with GRA24 KIM)) were fused to a twin StrepII tag with a 3C protease cleavage site and cloned into pFastBac1 (Fig. S3). For cellular reporter assays, $MEK1^{DD}$ mutant sequences were cloned into pcDNA3.1 plasmids.

Protein expression and purification

ERK1 and ERK2 construct vectors were transformed into Rosetta(DE3)pLysS *E. coli* competent cells (Novagen) with appropriate antibiotics. Cells were grown in LB at 37°C until $OD_{600}$ = 0.6-0.8, induced with 0.25 mM IPTG, expressed at 37°C for 3h, and harvested by centrifugation for 30 min at 4000 g. Cell pellets were frozen and stored at -75 °C.

MEK1 constructs were transformed into DH10 EMBacY *E. coli* cells (Thermo Fisher Scientific) to produce recombinant baculoviruses, subsequently used for protein expression in Hi5 insect cells (*59*). Cells were harvested 72 h after proliferation arrest by centrifugation and frozen.

Cell pellets of protein constructs were thawed on ice and resuspended in lysis buffer (50 mM HEPES pH 7.5, 200 mM NaCl, 10 mM $MgCl_2$, 5% glycerol, 0.5 mM TCEP, with a Pierce protease inhibitor EDTA-free tablet (Thermo Fisher Scientific) and a trace of DNaseI (Sigma)), before lysis by sonication on ice (40% amplitude, 5 s ON, 10 s OFF pulses for 2:30 min process time). The lysate was cleared by 1 h centrifugation at 45 000 g.

For ERK1 and ERK2 purification, the cleared supernatant was loaded onto a pre-packed 5 ml HisPur Cobalt (Thermo Fisher Scientific) affinity column, previously equilibrated in wash buffer (50 mM HEPES pH 7.5, 200 mM NaCl, 10 mM $MgCl_2$, 5% glycerol, 0.5 mM TCEP). Column bound protein was eluted by a step gradient of elution buffer containing 300 mM imidazole. Protein-containing samples were verified by SDS PAGE analysis, and fractions were mixed with 3C protease. The protein pool with 3C protease was dialysed into wash buffer at 4 °C overnight.

For MEK1 constructs, the supernatant was loaded onto a pre-packed 5 ml StrepTactin XT column (IBA), equilibrated in wash buffer (same as above). Bound protein was eluted with elution buffer containing 50 mM Biotin (IBA). Fractions were collected, analysed by SDS PAGE and pooled with 3C protease, followed by dialysis into wash buffer at 4 °C overnight.

Depending on the further use of the protein, affinity-purified and cleaved protein samples were concentrated in 10 kDa Amicon Ultra membrane concentrators (Millipore) and loaded onto a Superdex S200 Increase 10/300 GL size-exclusion column (GE Healthcare) equilibrated in wash buffer (as before). Peak fractions were analysed by SDS PAGE, pooled, and concentrated for immediate use or flash-freezing and storage at -75 °C (Fig. S3).

Mass spectrometry

*Intact mass*

Measurements were performed by the ISBG platform for MS (PSB, Grenoble). Samples containing 6 µg ERK2 and 1.5 µg MEK1 were diluted to a concentration of 1 µM using 0.1% formic acid (FA). Experiments were performed using a liquid chromatography (LC) electrospray (ESI) Quadrupole-Time of flight (Q-TOF) instrument (AdvanceBio 6545 XT, Agilent Technologies). The HPLC system was an Agilent 1260 Infinity II series. A PLRP-S column for biomolecules was used (PLRP-S 100Å, 1.0 x 50 mm, 3 µm, Agilent Technologies). HPLC mobile phases were prepared using HPLC-grade protonated solvents. HPLC solvents consisted of 0.1% FA in water (Solvent A); 99.9% acetonitrile and 0.1% FA (Solvent B). A gradient of HPLC solvents for a total time of 6 min was used. The solvent flow rate was 0.25 ml/min, and the column temperature was 60 °C. MS acquisition was carried out in positive ion mode. ESI source temperature was set at 325°C, nitrogen was used as drying gas (5.0 L/min) and as nebulizer gas (20 psi). The capillary needle voltage was set at 4000 V. Spectra acquisition rate was 1 spectra/s. The instrument was calibrated in a mass-to-charge (m/z) range 600–3000 using Myoglobin and BSA as standard calibrants (ESI-L, low concentration tuning mix, Agilent Technologies). The MS spectra were acquired and the data processed with Bioconfirm workstation software (v. B.3.4.242, Agilent Technologies) and with GPMAW software (v. 7.00b2, Lighthouse Data, Denmark). The displayed scores were normalized to the sum of intensities between 40 and 43 kDa measured for each run and plotted as a function of molecular weight (normalised intensity in arbitrary units (au), Fig. S4).

*Peptidic digest mass*

Measurements were performed by the protein core facility at EMBL Heidelberg. For sample preparation, protein bands of interest were excised from SDS PAGE gels and cut into approximately 1 mm$^3$ pieces to prepare for in-gel digestion, as previously described (*60*). All reagents were freshly prepared in 100 mM ammonium bicarbonate buffer. Gel pieces were first dehydrated by incubating in 100% acetonitrile. Subsequently, reduction was performed using 10 mM 1,4-dithiothreitol (DTT) for 30 minutes at 56°C. After reduction, the gel pieces were again dehydrated with acetonitrile and then alkylated in 55 mM 2-chloroacetamide (CAA) for 20 minutes at room temperature in the dark. A final dehydration with 100% acetonitrile was performed prior to digestion. Proteins were enzymatically digested overnight at 37°C using trypsin at a concentration of 2 ng/µL in 50 mM ammonium bicarbonate. Peptides were extracted by sonication for 15 minutes, followed by centrifugation and collection of the supernatant. For optimal peptide recovery, gel pieces were subjected to a second extraction step using a solution composed of 50% acetonitrile and 1% formic acid in ultrapure water. The volume of extraction solution added corresponded to twice the gel volume. Samples were again sonicated for 15 minutes, centrifuged, and the supernatants combined with the initial extracts. The pooled peptide extracts were dried using vacuum centrifugation. Dried peptides were reconstituted in a solution containing 4% acetonitrile and 1% formic acid prior to LC-MS/MS analysis.

Following MS measurements were performed with an UltiMate 3000 RSLCnano LC system (Thermo Fisher Scientific) equipped with a trapping cartridge (µ-Precolumn C18 PepMap™ 100, 300 µm i.d. × 5 mm, 5 µm particle size, 100 Å pore size; Thermo Fisher Scientific) and an analytical column (nanoEase™ M/Z HSS T3, 75 µm i.d. × 250 mm, 1.8 µm particle size, 100 Å pore size; Waters). Samples were trapped at a constant flow rate of 30 µL/min using 0.05% trifluoroacetic acid (TFA) in water for 6 minutes. After switching in-line with the analytical column, which was pre-equilibrated with solvent A (3% dimethyl sulfoxide [DMSO], 0.1% formic acid in water), the peptides were eluted at a constant flow rate of 0.3 µL/min using a gradient of increasing solvent B concentration (3% DMSO, 0.1% formic acid in acetonitrile).

Peptides were introduced into the Orbitrap Fusion™ Lumos™ Tribrid™ mass spectrometer (Thermo Fisher Scientific) via a Pico-Tip emitter (360 µm OD × 20 µm ID; 10 µm tip, CoAnn Technologies) using an applied spray voltage of 2.4 kV. The instrument was operated in positive ion mode, and the capillary temperature was set to 275 °C. Full MS scans were acquired in profile mode over an m/z range of 350–2000, with a resolution of 120,000 at m/z 200 in the Orbitrap. The maximum injection time was set to 50 ms, and the AGC target was set to ‘standard’. The instrument was operated in data-dependent acquisition (DDA) mode, with MS/MS scans acquired in the Orbitrap at a resolution of 15,000. The maximum injection time was set to 54 ms, and the AGC target was set to 400%. Fragmentation was performed using higher-energy collisional dissociation (HCD) with a normalized collision energy of 34%. MS2 data were acquired in profile mode, and dynamic exclusion was set to 30 seconds.

For database search, raw files were converted to mzML format using MSConvert from ProteoWizard, using peak picking, 64-bit encoding and zlib compression, and filtering for the 1000 most intense peaks. Files were then searched using MSFragger in FragPipe v.23 against the Uniprot database UP000000625 *E. coli* (4402entries; Oct 2022) containing common contaminants and reversed sequences as well as the amino acid sequences of ERK2wt; ERK2$^{T185V}$; MEK1$^{DD}$; MEK1$^{DD}$GRA constructs. The following modifications were included into the search parameters: Carbamidomethylation (C, 57.0215) as fixed modification; Oxidation (M, 15.9949), Acetylation (protein N-terminus, 42.0106), phosphorylation (STY, 79.96633) as variable modifications. For the full scan (MS1) a mass error tolerance of 20 PPM and for MS/MS (MS2) spectra of 20 PPM was set. For protein digestion, 'trypsin' was used as protease with an allowance of maximum 2 missed cleavages requiring a minimum peptide length of 7 amino acids. The false discovery rate on peptide and protein level was set to 0.01. The output files of the database search were loaded into Skyline v.25.1.0.237 (*61*) and phosphopeptides of interest as well as their matching counterpart peptides were quantified manually using MS1 extracted ion chromatograms.

### X-ray Crystallography

#### *Crystallisation, data collection and structure determination of the ERK1-GRA24KIM1complex*

In order to determine if the GRA24 KIM has similar effects on ERK2 as the native MEK1 KIM, we crystallised ERK1 with a peptide corresponding to the KIM1 sequence of GRA24. Human ERK1 was used instead of ERK2 as we were unable to obtain crystals with human ERK2. ERK1 and ERK2 share 84% sequence identity and 88% similarity. Crystallization conditions were established by testing several commercial screens at the EMBL High Throughput Crystallisation Laboratory (HTX). Crystals of ERK1 in complex with the peptide GLLERRGVSELPPLYI (Lifetein) were obtained at 21ºC in sitting drops from solutions containing 10 mg/ml ERK1 with a 3-fold molar excess of peptide, mixed with an equal volume of 0.1 M MES pH 7, 10% dioxane and 1.8 M ammonium sulphate. Crystals were harvested automatically by laser photoablation using the CrystalDirect robot (*62*) and data were collected at MASSIF-1 at the ESRF (*63, 64*) using automatic protocols for the location and optimal centring of crystals (*65*) on a Pilatus3 2M detector (Dectris). Strategy calculations accounted for flux and crystal volume in the parameter prediction for complete datasets. Crystals were hexagonal in morphology and the crystal from which the best data set was collected had the dimensions: 0.107 x 0.079 x 0.101 mm$^3$. The complex crystallised in the hexagonal space group *P*6522, with one molecule in the asymmetric unit, and diffracted to a resolution of 2.17 Å. Data were processed with XDS (*66*) within the Autoproc pipeline (*67*) and phases were calculated by molecular replacement with Phaser (*68*) using the ERK1 structure PDB 4QTB (*69*) as a starting model and refined with a combination of Refmac (*70*), Phenix (*71*), and Coot (*72*) (Fig S2). Data processing and refinement statistics can be found in Table S1.

### AlphaFold predictions

For predictions of MEK1-ERK2 complexes the sequences of the full length proteins and variants were submitted to the AlphaFold3 (*28*) server (https://alphafoldserver.com/) and used for initial model building and composite models.

Cryo-EM sample preparation, data collection and processing

*Cryo-EM specimen preparation and data collection*

SEC co-purified MEK1$^{DD}$GRA-ERK2$^{T185V}$ was diluted to 4 µM, supplemented with 250 µM ADP, 250 µM $NH_4F$ and 25 µM $AlCl_3$, and incubated on ice for 30 minutes (*32*). For grid preparation before sample application, HexAuFoil grids (Quantifoil) were hydrophilized with a NanoClean 1070 plasma cleaner (Fischione) at 100% power, 20:80 oxygen:argon gas mix, for 3 min. Grid vitrification was performed on a Vitrobot Mark IV (FEI), applying 3.5 µl of sample per grid, and blotting for 4-6 s at blot force 0. Grids were then clipped and screened on a FEI Talos Glacios electron microscope (EMBL Grenoble) operating at 200 kV. Selected grids were taken for data collection on a 300 kV operating TFS Titan Krios electron microscope (CM02, CRG beam line at ESRF, Grenoble). Micrographs were collected using EPU from a Falcon 4i direct electron detector with a SelectrisX energy filter. A total of 88512 movies of 58 raw frames were collected in TIFF format at 270,000 magnification, a pixel size of 0.46 Å with a total exposure of 50 $e^-$/Å$^2$.

*Cryo-EM processing*

Collected movies were imported into cryoSPARC (*35*). All movie frames were aligned and motion-corrected with Patch Motion correction and CTF estimated with Patch CTF. Data were manually curated to eliminate micrographs with large motions, poor resolution, and high ice thickness. Picking was most successful through rounds of Topaz model training and particle picking (*34*). Originally, particles were extracted with a box size of 600 x 600 pixels and four times binned. 2D classification was used to eliminate junk and the noisiest particles. Once a set of aligning particles was identified, a box size of 440 x 440 was used for extraction and twice binned. Rounds of *ab initio* and heterogeneous refinement, requesting 3-6 models, were started, and particles corresponding to the hetero kinase complex were selected for further rounds of 2D and 3D classification. Multiple rounds of classifications were necessary to select good particles, showing secondary structure features and low background noise, leading to a stack of ~800,000 particles. Variability analysis helped to select more homogenous sets of particles and heterogeneous refinement and 3D classification, applying a focused mask on the region between the kinases (Fig. S4), allowed the separation of different states. Initially, non-uniform refinement was used to obtain ~ 3.5 Å resolution maps of different subsets of particles. Applying the HR-HAIR (*36*) strategy immensely improved the overall quality of maps and helped to achieve better alignment of particle stacks. The workflow is shown in Figure S4. The combination of these specific *ab initio* runs with homogeneous reconstruction and local refinement yielded the final three volumes presented. The final volumes were obtained to an overall resolution between 2.99 Å and 3.63 Å (Table S2 and Fig. S6).

*Model building and refinement*

For postprocessing, B-factor sharpening was applied, which helped side chain visualization for model building. LocScale feature-enhanced (*73*) mode was used for creating figures with maps. Refinement programs ISOLDE (*74*), phenix.realspace refine (*75*) and Servalcat (*76*) were used for model refinement. Initial models were based on a combination of AlphaFold3 predictions and crystal structures of MEK1, ERK2 and ERK1. Particularly, the manually curated active form of MEK1 (*77*) was helpful, downloaded from KinCore (https://dunbrack.fccc.edu/kincore/). Data processing and refinement statistics are shown in Table S2.

Luciferase reporter assay

HEK293T cells were maintained in DMEM medium supplemented with 10% (v/v) FBS and 1% Pen-Strep (all Thermo Fisher Scientific) at 37 °C and 5% $CO_2$. The SRE Reporter Kit (BPS Bioscience #60511) was used according to the manufacturer's protocol. In short, white 96-well cell-culture plates (greiner) were seeded on day 1 with 300,000 cells in 100 µl per well after media exchange into BPS media (BPS Bioscience). After 24 h, a mix of 60 ng of the SRE firefly luciferase reporter vector pre-mixed with the constitutively expressing renilla luciferase vector were transiently co-transfected, together with 50 ng MEK1$^{DD}$ mutants, in pcDNA3.1 plasmids (or 50 ng empty vector and 60 ng non-inducible reporter for controls), with 0.35 µl Lipofectamin2000 and 15 µl Opti-MEM (Thermo Scientific) per well. Cell media was exchanged to 50 µl Opti-MEM with 0.5% FBS 6 h after transfection. Firefly and renilla luciferase activities were measured 24 h post-transfection using the Dual Luciferase assay system (BPS Bioscience #60683) with a Clariostar microplate reader (BMG Germany). Focal point and gain were determined per plate using positive and negative controls, measurements were recorded using end-point kinetics settings and measuring in spiral per well. Each experimental plate included following controls: cell media only, seeded cells without transfection, cells transfected with empty DNA only, cells transfected with empty DNA and non-inducible reporter and cells transfected with empty DNA and inducible reporter; the last condition was used as negative control, accounting for unspecific background firefly signal. For each experiment, three transfections per condition were performed on the same plate (technical replicates), and each experiment was repeated three times (experimental replicates), for a total of nine measurements per construct. The relative luminesence (RL) of a condition was obtained by normalising each firefly luciferase value by its corresponding renilla luciferase measurement from the same well, to correct for cell quantity and transfection efficiency, and by then averaging the firefly/renilla ratios from the three technical replicates. To compare the RL of mutants to the positive control (MEK1$^{DD}$), a Shapiro-Wilk test was performed on the mutant RL triplicates to test for normal distribution, and if successful, a paired t test was used to test significance (tests performed with GraphPad Prism v11, RLs and statistical values shown in Table S3). To present the data, the mutant RL was expressed relative to the average RL of the positive control (MEK1$^{DD}$) activity using a relative response rate calculation: (condition RL - negative control RL)/(positive control RL - negative control RL), where MEK1$^{DD}$ and the inducible promoter with empty DNA condition were used as respective controls and therefore set as 100% and 0% (Table S4, Fig. S10).

Western blots

For Western blots, technical triplicate samples from the luciferase reporter assay were pooled and spun down for 15 min at 15 000 g, and the supernatant recovered. Renilla luminescence readings from the reporter assay were used to normalise protein content per sample. Each sample was mixed with 5x loading buffer, loaded on a Tris-Glycine 4–20 % gradient gel (Invitrogen) and run at 125 V for 1:40 h. Transfer onto a nitrocellulose membrane was performed using a semi-dry transfer system (#1704158, BioRad). For blot staining, mouse anti-myc (#2276, Cell signaling) at 1:5000, and rabbit anti-ERK1/2 (#137F5, Cell signaling) at 1:1000 dilution were used for detection of transfected myc-MEK1$^{DD}$ mutants and native ERK1/2 respectively, whilst mouse anti-β-tubulin antibody (MA5-16308, Thermo Fisher Scientific) at 1:5000 dilution was used as loading control. Secondary antibodies linked to fluorophores, goat anti-rabbit Alexa 488 and goat anti-mouse Alexa 647 (A-11008 and A32728, Thermo Fisher Scientific) were used at 1:10000 dilution. Blots were visualised using a BioRad ChemiDoc system (Fig. S11).

Hydrogen-deuterium exchange mass spectrometry

HDX-MS experiments were performed at the UniGe Protein Biochemistry Platform (University of Geneva, Switzerland) following a well-established protocol with minimal modifications (*78*). The supplementary data S1 and S2 (for each protein) detail all the reaction conditions as well as all results (raw; HDX levels and; HDX diff worksheets). The following conditions were tested and compared: i) MEK1, ii) ERK2, iii) MEK1:ERK2 (ratio 1:1.5) and iv) ERK2:MEK1 (ratio 1:1.5), all supplemented with 10 mM ADP, 1 mM $AlCl_3$, 10 mM $NH_4F$. Briefly, a first concentrated mix was prepared for each of the conditions, preincubated for 5 min at the reaction temperature, before initiating deuteration reaction. Deuterium exchange reaction was initiated by adding deuterated buffer at 22 °C to a final volume of 50 µl. Reactions were carried-out for 3 sec, 30 sec and 5 min and terminated by the sequential addition of ice-cold quench buffer. Samples were immediately frozen in liquid nitrogen and stored at -80 °C for up to two weeks. Each condition was performed in triplicate.

To quantify deuterium uptake into the protein, samples were thawed and injected in a UPLC system immersed in ice with 0.1 % FA as liquid phase. The protein was digested via an immobilized Nepenthesin-2/Pepsin mixed column (AffiPro AP-PC-006), and peptides were collected onto a Nucleodur 300-5 C18 metal-beads pre-column filter (Macherey-Nagel). The trap was subsequently eluted, and peptides separated with a C18, 175 Å, 1.9 µm particle size Thermo Hypersil Gold Vanquish 100 x 2.1 mm column over a gradient of 8 – 30 % buffer C over 20 min at 150 l/min (Buffer B: 0.1% formic acid; buffer C: 100% acetonitrile). Mass spectra were acquired on an Orbitrap Velos Pro (Thermo), for ions from 400 to 2200 m/z using an electrospray ionization source operated at 300 °C, 5 kV of ion spray voltage. Peptides were identified by data-dependent acquisition of a non-deuterated sample after MS/MS and data were analyzed by Mascot 2.6 using a database composed of purified proteins and known contaminants. Precursor mass tolerance was set to 10 ppm and fragment mass tolerance to 0.6 Da. Protein digestion was set as nonspecific. All peptides analysed are shown in HDX supplementary tables. Deuterium incorporation levels were quantified using HD examiner version 3.4.2 software (Sierra Analytics), and the quality of every peptide was checked manually. Results are presented as a percentage of maximal deuteration compared to theoretical maximal deuteration. For each protein, criteria to define a change in deuteration level between two states as significant are indicated in the table.

Isothermal Titration Calorimetry

ITC measurements were performed using the Microcal PEAQ-ITC (Malvern Panalytical GmbH) at 20°C with the following settings: 13 injections of 3.0 µL spaced by 150 seconds using a stirring speed of 750 rpm. Before the experiments, samples were purified using SEC in ITC buffer: 50 mM HEPES pH 7.5, 200 mM NaCl, 10 mM MgCl2, 5% glycerol and 0.5 mM TCEP. The GRA24 KIM peptide was ordered from SBpeptide, and the MEK1 KIM peptide from LifeTein. The peptides were resuspended directly in the ITC buffer, and the concentration was determined according to the volume added to the 1 mg peptide powder, and the pH adjusted where necessary. For protein-peptide experiments, 30 µM $ERK2^{WT}$ was titrated by 280 µM or 600 µM of MEK1 KIM peptide, or titrated by 280 µM of GRA24 KIM peptide. 20 µM $ERK1^{WT}$ was titrated by 200 µM of GRA24 KIM peptide. For protein-protein experiments, 20 µM $MEK1^{WT}$, $MEK1^{DD}$, or $MEK1^{DD}$GRA was titrated by 280 µM $ERK2^{WT}$. All ITC measurements were carried out in triplicate. Control experiments (buffer into protein and peptide into buffer) were also performed. The fitted offset option, in PEAQ-ITC analysis software, was used to correct for the heat of dilution. The data were fitted with a one set of sites binding model using the MicroCal PEAQ-ITC analysis software. See Table S5 for details.

Small Angle X-ray scattering

Size-exclusion chromatography-coupled small-angle x-ray scattering (SEC-SAXS) data were collected at the BioSAXS beamline BM29, ESRF, Grenoble, France (*79*). To promote sample monodispersity, samples were eluted from a GE Superdex 200 Increase 10/300 column connected to a Shimadzu HPLC system. The mobile phase consisted of 50 mM HEPES pH 7.5, 200 mM NaCl, 10 mM $MgCl_2$, 2.5% glycerol, 0.5 mM TCEP at a flow rate of 0.5 ml/min. The eluate passed through a 1 mm quartz capillary at 20 ℃, where it was exposed to X-rays ($\lambda$ = 0.99 Å). Scattering was recorded as 1500 continuous 2-second frames on an in-vacuum Dectris Pilatus3 X 2M detector, at a sample to detector distance of 2.827 m (full details in Table S6 and Fig. S16-18).

FreeSAS (*80*) was used for automated data reduction, integration, and buffer subtraction. Final scattering profiles were generated by averaging frames selected based on summed intensity across the elution peak (Supplementary Table 4. Initial Rg and forward scattering intensity (I(0)) values were determined via Guinier analysis in BioXTAS RAW (*81*). The pair-distance distribution function, P(r), and maximum particle dimension (Dmax) were calculated using the Bayesian Indirect Fourier Transform (BIFT) method (*82*). Molecular weight was estimated using multiple methods: Bayesian (*83*), Volume of Correlation (Vc) (*84*), and Porod Volume (Vp) (*85*). Electron density was reconstructed using DENSS (*86*). Atomic models were generated from the cryo-EM structure of $MEK1^{DD}$GRA-$ERK2^{T185V}$, with missing residues modelled via ColabFold (*87*). Theoretical scattering profiles were calculated and fitted to the experimental data using PEPSI-SAXS (*49*), assessing quality of fit via reduced $\chi^2$ and residual analysis. Model refinement was conducted using the PEPSI-SAXS optimisation function with non-linear normal modes (NOLB) (*88*).

Protein structure preparation for MD simulations

Three models of $MEK1^{DD}$GRA in complex with $ERK2^{T185V,}$ as obtained from cryo-EM maps, were used as starting structures for the molecular dynamics simulations. These differ in the $MEK1^{DD}$GRA conformation and are named as follows:

1. *Fully active:* The starting structure was derived from the state 2 model. The A-loop is on the side of MEK1, leaving the binding site open and accessible to the activation loop of $ERK2^{T185V}$, the MEK1 αC is in an *in* position and the catalytic spine was put in place.
2. *Active:* As for the *fully active* conformation, the starting structure was derived from the state 2 model, with MEK1 A-loop unfolded on the side leaving the binding site open and accessible to the activation loop of $ERK2^{T185V}$. Here, however, the αC has moved towards the *in* position but the MEK1 catalytic spine is not in place.
3. *Inactive:* The starting structure was derived from the state 1 model. The A-loop of MEK1 is organized as a short alpha helix at the interface between MEK1 and ERK2, effectively blocking the binding site from the activation loop of $ERK2^{T185V}$. MEK1 αC is in an *out* position.

In all cases, the ADP nucleotides are resolved but not the corresponding coordinating magnesium ions. Therefore, the coordinates of ATP and Mg atoms were taken from similar kinase structures by aligning the DFG motifs and the nucleotides of the previously published structures on those here resolved by cryo-EM. In all model complexes, the coordinates of ATP and the two Mg atoms for $ERK2^{WT}$ were taken from PDB ID 5V60 (*89*). The coordinates of the nucleotide and the Mg atoms for the *fully active* and *active* structures of $MEK1^{DD}$GRA were taken from PDB ID 3FJQ (*90*), while those for the *inactive* structure from PDB ID 6U2G (*91*). These structures were chosen with the rationale of being as similar as possible in sequence, while also containing a bound nucleotide, the two coordinating metals, and no other ligand in the binding site. Notably, in all these PDB structures, the nucleotide resolved is AMP-PCP, an ATP analogue. Thus, the 3D coordinates of AMP-PCP were retained and the molecule was simply transformed into ATP by manually reverting to a phosphoanhydride oxygen. Similarly, in PDB ID 3FJQ the resolved divalent ions are manganese atoms, which have been converted here to Mg. The structures were then refined and optimized with Schrödinger's Maestro (v.2025), and the ATP molecules were redocked using the Glide software (*92*), taking care that the geometry and main interactions between the binding site, nucleotides, and coordination ions were maintained and consistent with the cryo-EM models. When relevant, co-crystallized water molecules coordinates were also taken from the corresponding deposited PDB structures.

The phosphomimetic S218D and S222D mutations in MEK1$^{DD}$GRA were reintroduced if not present in the fitted structure, while T185V in ERK2 was reverted to the corresponding WT threonine. Point mutations were introduced with PyMOL (https://pymol.org). The GRA KIM of the three MEK1$^{DD}$GRA-ERK2$^{WT}$ models was then reverted to WT with MODELLER *automodel* function (*93*). The coordinates of nucleotides and Mg ions were the same for both GRA and WT KIM models, as the mutations are not part of the binding sites. The protonation state of the residues at pH 7.4 was calculated with PROPKA3.1 (*94*) via the PlayMolecule web application (*95*). The residues were left in their usual charge state. Throughout the setup procedure, the correctness of the final models sequences was verified via the *EMBOSS stretcher* tool (https://www.ebi.ac.uk/jdispatcher/psa/emboss_stretcher).

The C-terminus of MEK1 and the N-terminus of ERK2 were shortened by 13 residues (after G380 onwards, in WT numbering) and 9 residues (up to G10), respectively, and were capped with NME and ACE groups to avoid a terminal net charge. These terminal residues are extremely disordered regions that are not resolved by the cryo-EM map and are not part of the MEK1-ERK2 interaction interface. Thus, they were removed to avoid PBC self-interaction artefacts and restrict the simulation box volume to speed up the MD simulations.

Finally, six models of the kinase heterodimers were obtained, that is MEK1$^{DD}$GRA-ERK2$^{WT}$ and MEK1$^{DD}$-ERK2$^{WT}$ complexes in inactive, active, and fully active states (see also Table S7).

Molecular dynamics simulations

All the systems were simulated with the DES-Amber force field (*96, 97*). For ATP and the coordinating $Mg^{2+}$ ions, the parametrization from Meagher *et al.* (*98*) and Grotz and Schwierz (*99*) were used, respectively. These parameters have been optimized to reproduce the coordination modes of ATP and $Mg^{2+}$, and have been already successfully combined in similar kinase MD simulations, e.g. MKK6-p38α (*15*).

Each system was enclosed in a dodecahedron box, ensuring at least 1.1 nm of space between all atoms of the dimer and the box edges. The box was solvated with TIP4PD water molecules (*100*), as suggested for the DES-Amber force field. $Na^+$ and $Cl^-$ ions were added to neutralize the systems' charge and reach the NaCl concentration of 150 mM. Notably, also the parameters for $Cl^-$ were rescaled following Grotz and Schwierz (*99*) to optimize the interaction with the Mg ions. Eventually, all systems comprised ca. 140k atoms.

The MD simulations were run with GROMACS 2025.2 (*101*). The simulation details and the equilibration procedure was the same for all the systems. First, the box's potential energy was minimized via a steepest descent integrator. Then, the system was thermalized at 310 K in the canonical ensemble (NVT) for 3 ns followed by a first round of 70 ns equilibration in the isothermal-isobaric ensemble (NpT). A set of position restraints were applied throughout the equilibration phases to gradually relax the starting frames and avoid introducing structural artifacts. The restraints were applied on both side chains and backbone atoms of the kinases as well as on other selected important degrees of freedom, such as the Mg ions positions, the phosphate positions of the nucleotides, the coordinates of the aspartic acid in the DFG motifs, and the main h-bonds between nucleotides and the respective binding sites. These harmonic restraints gradually decreased in coupling strength, generally going from ~1000 to ~10 kJ $mol^{-1}$ $nm^{-2}$. Lastly, the restraints were turned off for a completely unbiased simulation where the box was further equilibrated for 50 ns in the NpT ensemble before being considered fully equilibrated and moved to production. For each of the six systems (*fully active*, *active* and *inactive* states for both GRA and WT KIMs) a total of 8 independent replicas were run, each for 2 µs (Table S7). To enhance the independence of the replicas and ensure that the production runs did not spawn from the same identical state, for at least four replicas of each system set the full equilibration pipeline was restarted. For the others in the same set, the velocities of the last equilibration frame were randomly resampled, the unbiased equilibration was separately elongated for 100 ns, and that final frame was used to start the corresponding production runs.

The initial velocity distributions were sampled from the Boltzmann distribution at 310 K. The temperature was kept constant at 310 K by a velocity-rescale thermostat (*102*) with a coupling constant of 1 ps, while the pressure was maintained at 1 bar through an isotropic cell-rescale barostat (*103*) with a coupling constant of 5 ps and a compressibility of $4.5x10^{-5}$ $bar^{-1}$. Periodic boundary conditions were employed in all three box dimensions. The long-range electrostatics were calculated by the particle mesh Ewald algorithm (*104*), with a Fourier spacing of 0.12 nm, combined with a potential-shift switching function between 0.0 and 1.0 nm for the electrostatic and vdW interactions, while short-ranged vdW interactions were cut-off at 1.0 nm. H-bonds were constrained with LINCS (*105*) and thus Newton's second equation of motion was integrated with a 2 fs timestep with the GROMACS' *md* leap-frog algorithm.

Molecular dynamics simulations analysis and visualization

The trajectories were PBC corrected and cleaned for analysis with GROMACS native tools. Snapshots for analysis were collected every 0.1 ns (50000 steps) for a total of 20000 frames per trajectory. Distance and flexibility analysis were run with GROMACS *distance* and *rms* native tools. Specifically, RMSF calculations were run over the Ca atoms with respect to the corresponding starting configurations. Contact map analysis was run in MDAnalysis (*106*) on all the trajectories after concatenating them. Contacts were computed via proximity matrices with a cut-off of 4.5 Å on the heavy atoms of the side chains. Synthetic HDX-MS profiles were computed on the concatenated trajectories in python3 with HDXer (*107*).
Post processing of data and further analysis was run with NumPy (*108*) in JupyterLab.
Figures were generated in Matplotlib (*109*), ChimeraX (*110*), and Inkscape (https://inkscape.org).

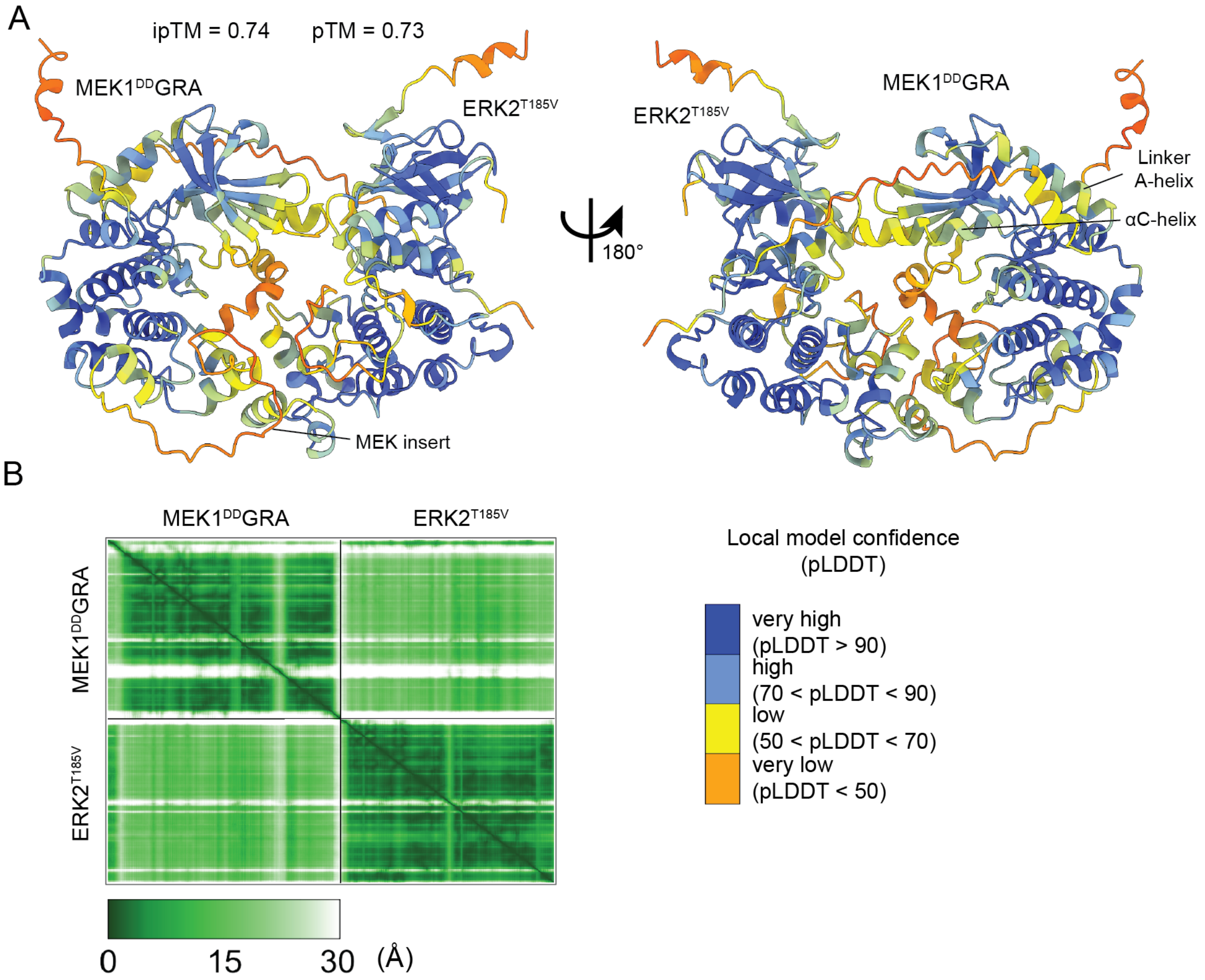


**Fig. S1. AlphaFold3 (AF3) prediction of the MEK1$^{DD}$GRA-ERK2$^{T185V}$ complex. A** Cartoon model of the complex coloured by pLDDT, value legend under the models, ipTM and pTM values of prediction above the model. **B** Predicted alignment error (PAE) plot for the AF3 MEK1$^{DD}$GRA-ERK2$^{WT}$ prediction. Dark green regions indicate confident global placement of residues, light green areas display high error values and indicate flexibility.

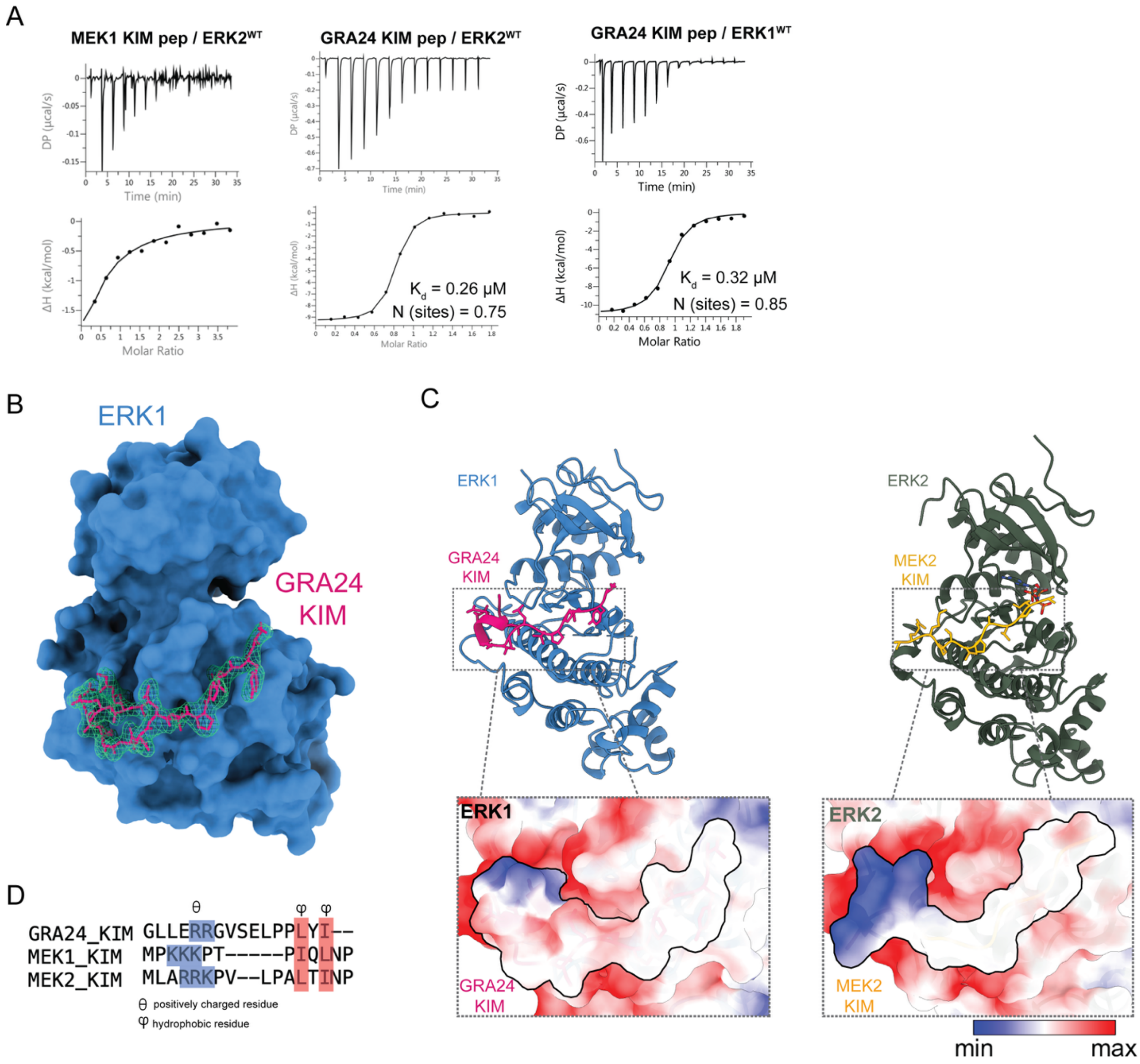


**Fig. S2. The GRA24 KIM1 peptide binds ERK1 and 2 with high affinity and induces the same conformation of ERK1 as the MEK2 KIM peptide. A** ITC binding curves of KIM peptides. Representative graphs with the full experimental list in Table S5. The left panel with native MEK1 KIM peptide shows too weak binding for reliable $K_d$ extraction. $K_d$ values of the experiment for GRA24 KIM1 peptide towards ERK1 and ERK2 are shown (calculated mean from three independent experiments yields a $K_d$=0.31 µM for both). **B** Crystal structure of the ERK1-GRA24KIM1 complex, omit density (green mesh, 3σ) is shown for the GRA24 KIM. **C** Comparison of the ERK1-GRA24KIM1 complex and the crystal structure of ERK2 bound to the MEK2 KIM peptide (PDB 4H3Q), the RMSD between the kinase Cα atoms = 0.79 Å. Zooms show the similarity on binding modes, surface charge and hydrophobic interactions. **D** Alignment of the KIM peptides for GRA24, MEK1 and MEK2.

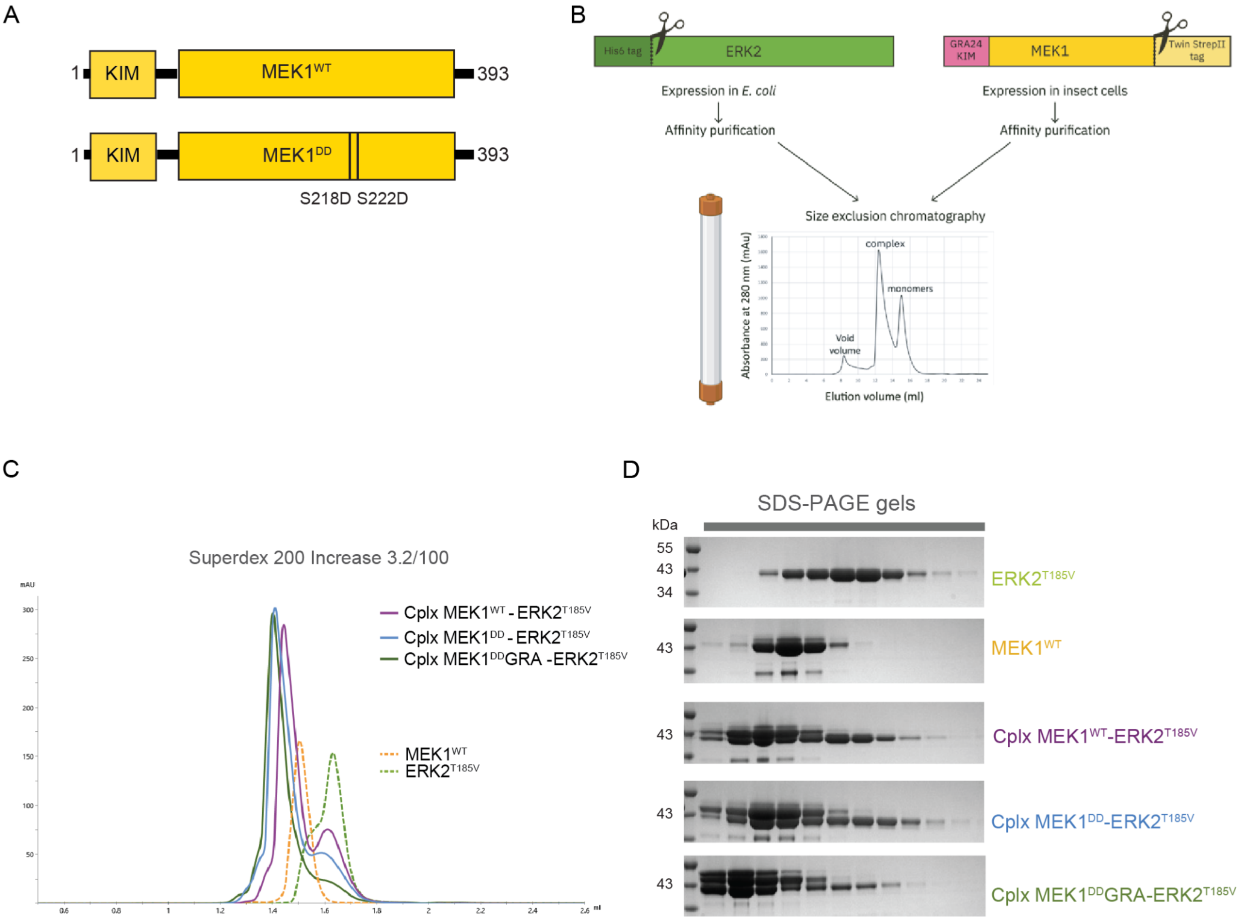


**Fig. S3. Purification of MEK1-ERK2 complexes. A** Schematic representation of further MEK1 protein constructs used for solution studies. **B** Exemplary 2D classes of a cryo-EM dataset collected on the MEK1$^{DD}$-ERK2$^{T185V}$ complex. **C** Schematic of purification strategy for individual kinase expression and complex reconstitution before SEC. The single kinases elute later than the complex. **D** Size-exclusion chromatograms of individual runs overlaid. The same column, and the same amount of protein was injected per run. Runs injected with only single kinases are shown with a dashed line, runs with both kinases mixed pre-injection are shown as a solid line. The grey bar under the elution volume indicates loaded fractions on SDS gel, shown in C. **E** Coomassie-stained SDS PAGE gels of SEC purification, as shown in D.

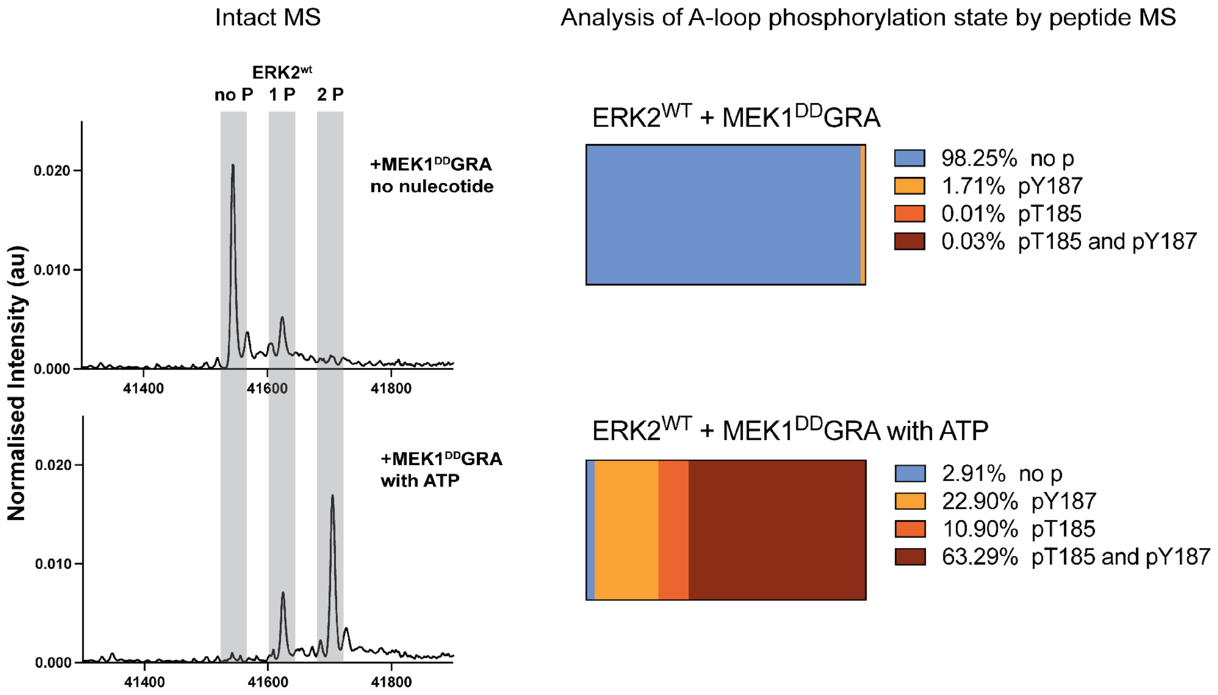


**Fig. S4. The reconstituted complex is active, as shown by mass spectrometry (MS) analyses.** MS analysis of ERK2$^{WT}$ after addition of MEK1$^{DD}$GRA and incubation for 2 h before flash-freezing until analysis, with no nucleotide (top row) and with 1 mM ATP (bottom row). Intact MS analysis (left panel) shows phosphorylation states of ERK2$^{WT}$ by a shift of 80 Da (1P) or 160 Da (2P) in mass compared to the unphosphorylated peak (no P). Peptide analysis by LC-MS/MS (right panel) shows the occupancy of phosphorylated residues of the A-loop TxY motif on ERK2$^{WT}$. Without nucleotide, the majority of ERK2$^{WT}$ is unphosphorylated. In the presence of ATP, the majority of ERK2$^{WT}$ is either mono- or double-phosphorylated on the A-loop. Interestingly, both T185$^{P}$ and Y187$^{P}$ mono-phosphorylated forms are present.

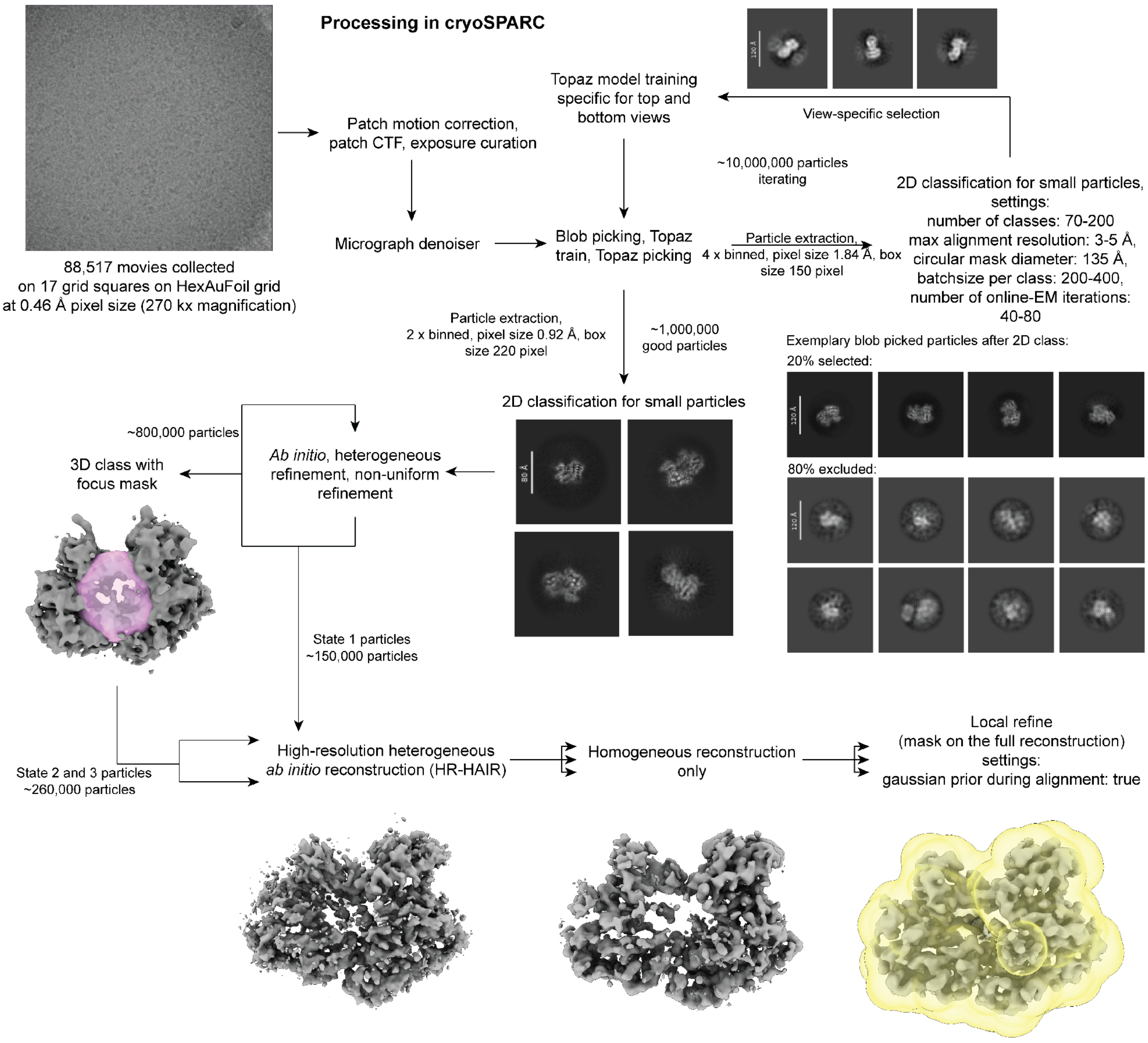


**Fig. S5. Processing pipeline used to determine the 3 states of the MEK1$^{DD}$GRA-ERK2$^{T185V}$ complex.** Details in materials and methods.

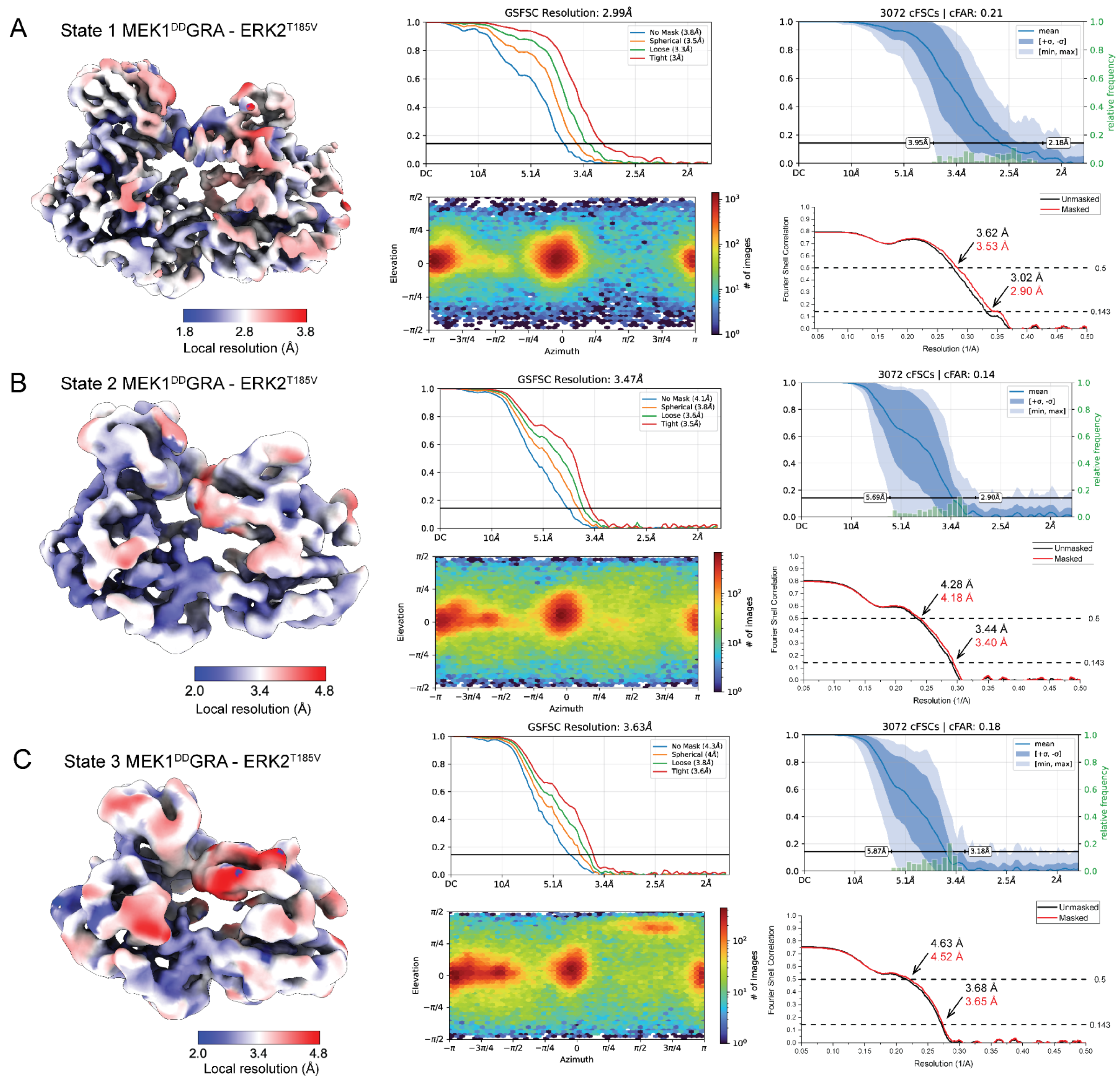


**Fig. S6. Cryo-EM processing and data assessment of the three cryo-EM reconstructions.** From left to right: reconstructions of MEK1$^{DD}$GRA-ERK2$^{T185V}$ coloured by local resolution, FSC half map curves with 0.143 cutoff, viewing direction distribution, cFSC graph, and model vs map (masked in red and unmasked in black) FSC curves (Nyquist = 0.54 Å$^{-1}$) are shown for the MEK1$^{DD}$GRA-ERK2$^{T185V}$ state 1 (**A**), state 2 (**B**) and state 3 (**C**) structures.

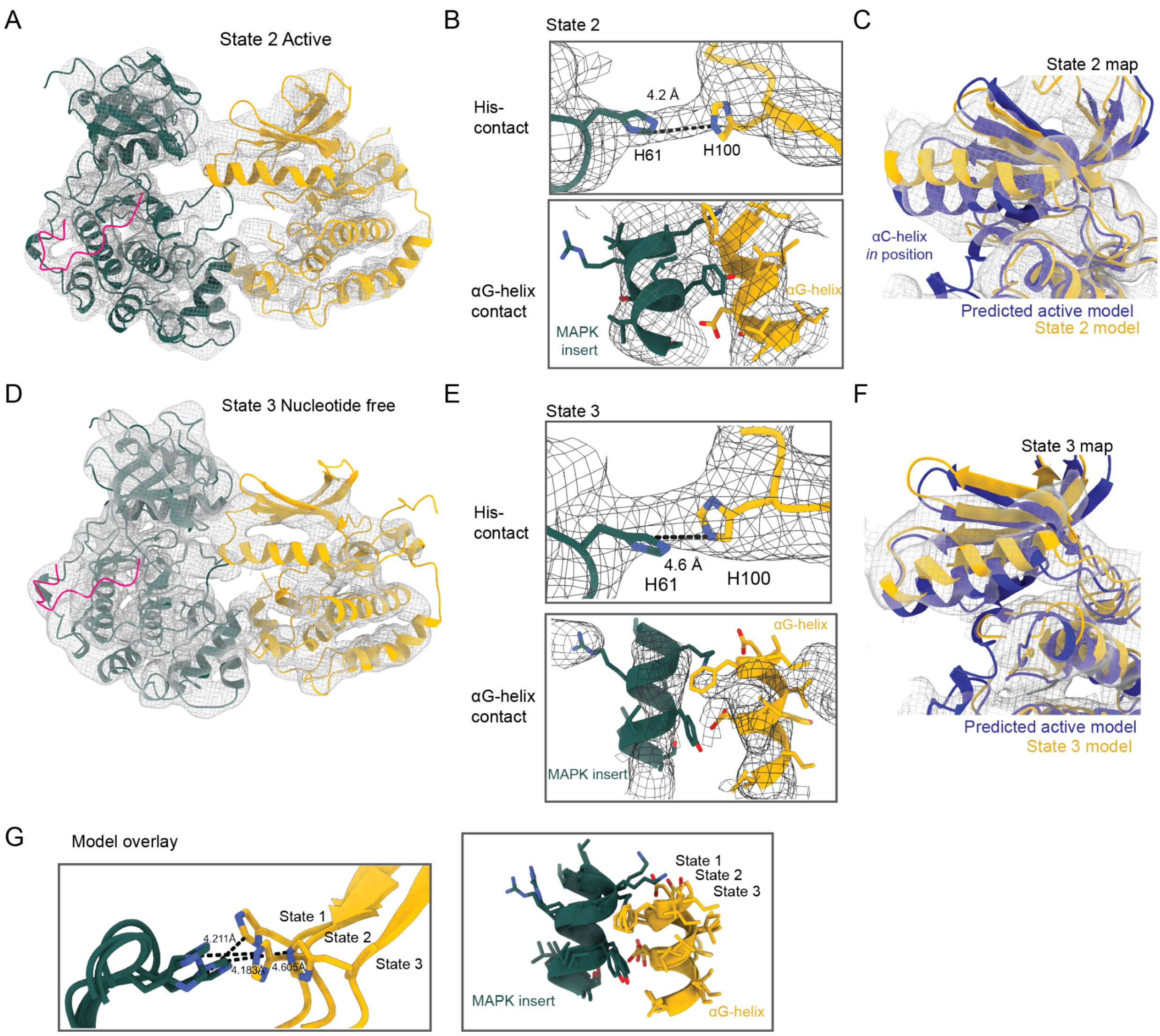


**Fig. S7. Detailed views of MEK1$^{DD}$GRA-ERK2$^{T185V}$ complex models with Coulomb potential maps.** **A** Model to Coulomb potential fit for state 2. **B** Details of His100-His61 interaction and αG-helix contact site between MEK1$^{DD}$GRA and ERK2$^{T185V}$ for state 2 MEK1 and ERK2 are colored as before with Coulomb potential in mesh. **C** Fit of predicted active MEK1 model (downloaded from KinCore, (*38*)) model with, αC-helix 'in', overlaid with the state 2 model in the unsharpened Coulomb potential displayed as mesh. **D, E, F** Same as A, B, C for state 3 map and model. **G** Overlay of the His and αG-helix site for the models from the three states.

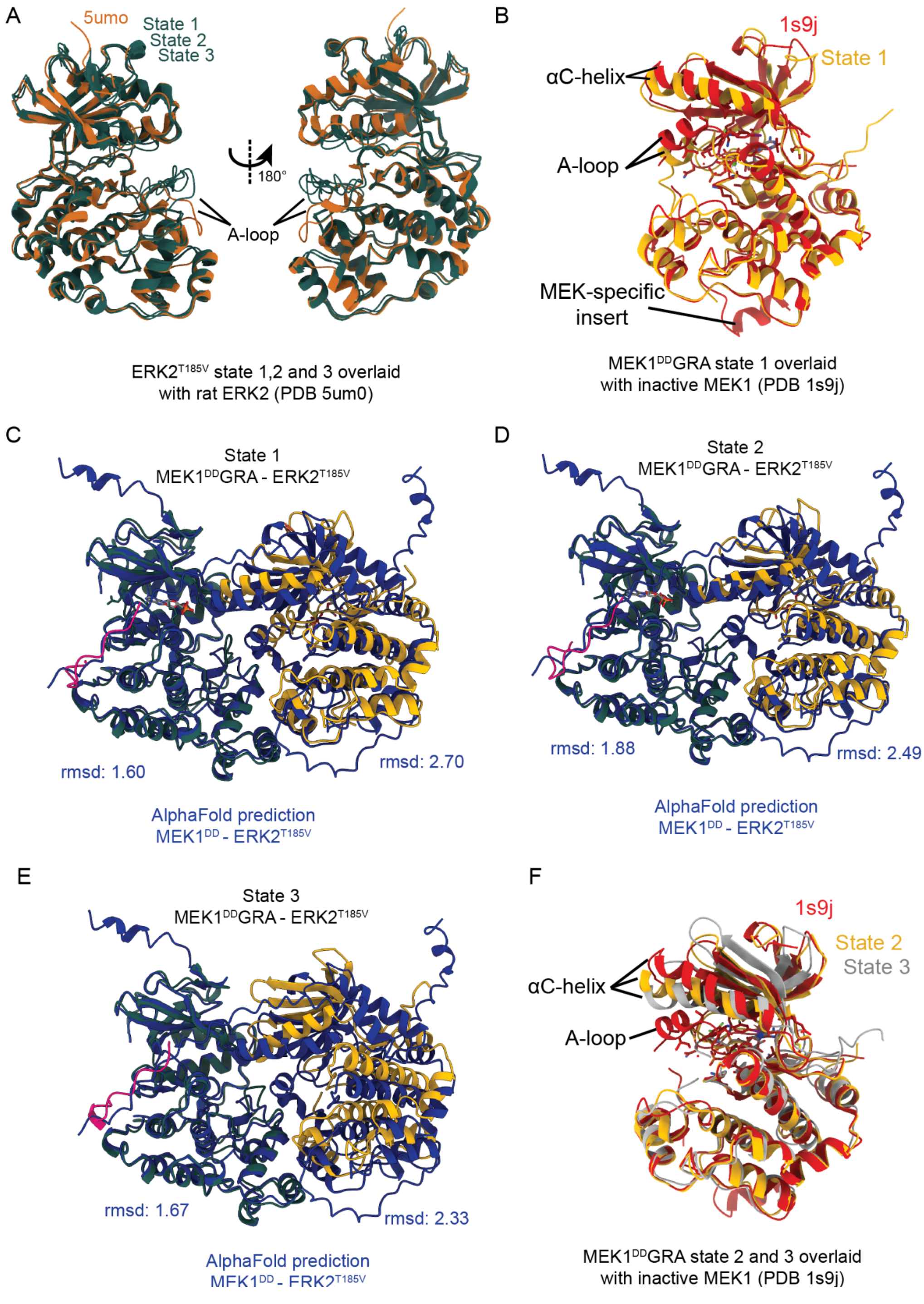


**Fig. S8. AlphaFold3 (AF3) prediction and crystal structure models compared to cryo-EM models. A** Comparison of ERK2 crystal structure (PDB 5umo, orange) to State 1, 2, and 3 model (green) from this study. A-loop is labelled to highlight minor differences in placement. **B** Comparison of state 1 model with inactive human MEK1 crystal structure bound to an inhibitor (PDB 1s9j, coloured red). Sites with differences are highlighted with labels. **C, D, E** Comparison of the three states cryo-EM models with the AF3 prediction aligned on the substrate MAPK ERK2. RMSDs are shown for the individual kinase-kinase main chains. **F** Same as B but for state 2 and 3.

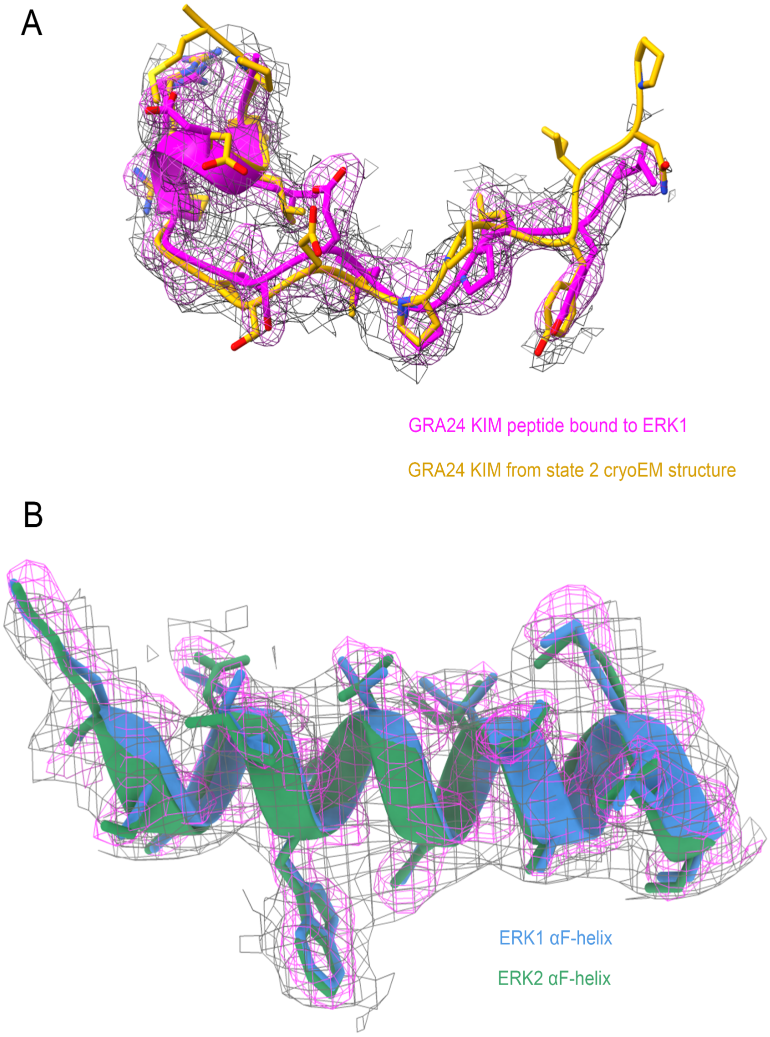


**Fig. S9. Comparison between the cryo-EM model and crystal structure of the GRA24 KIM binding to ERK1/2.** The sharpened Coulomb potential (grey mesh contoured at 0.0552 (2.1σ)) of the cryo-EM structure of State 1 compared to the electron density (magenta mesh contoured at 1.5σ) from the ERK1-GRA24KIM1 crystal structure. **A** The GRA24 KIM1 motif (cryo-EM model in yellow, crystal structure in magenta) and **B** the ERK kinase (cryo-EM model of ERK2 in green, crystal structure of ERK1 in blue).

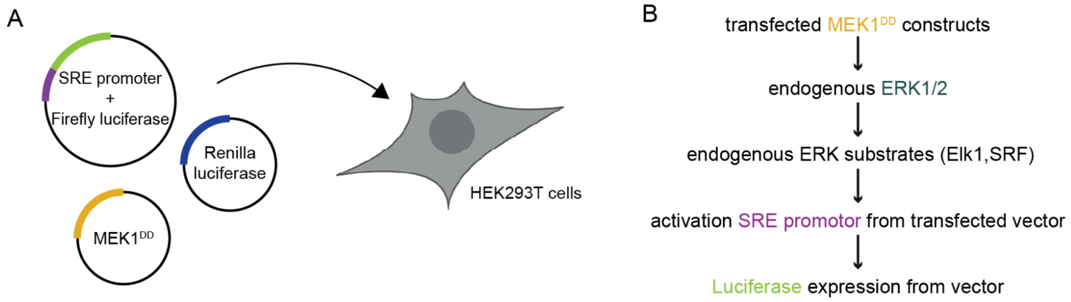


C

MEK1$^{DD}$ mutants:

| | |
|---|---|
| GRA | Used for structural studies, schematic displayed in Fig. 1B |
| H100A / H239A | Histidine to alanine point mutant at residue 100 or 239 |
| dblH-mutant | Residue 100 and 239 mutated to alanine |
| F311A | Phenylalanine to alanine point mutant at residue 311 in αG-helix |
| K57N, ΔEI | MEK1 mutants found in cancer patients: lysine 57 on A-helix and deletion of E102 and I103 at ß3-αC |
| Ascreen | A-helix mutant with residues 49-60 replaced by alanines |
| T55L | Threonine to leucine point mutant on the A-helix, increasing helical propensity |
| Δinsert | Deleted residues 267-306, removing MEK-specific insert |
| ΔKIM | Deleted residues 1-13, removing KIM-motif |

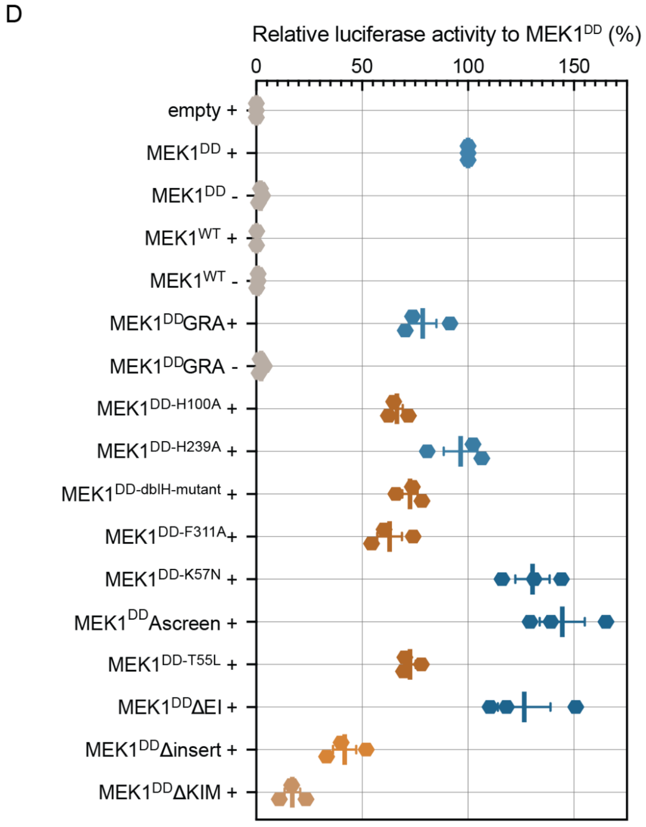


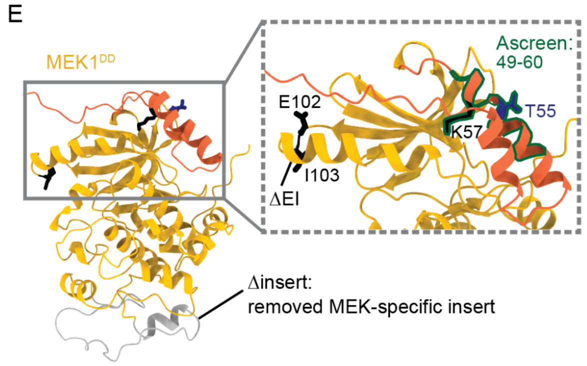


**Fig. S10. Reporter cell assay with MEK1$^{DD}$ and mutants. A** Diagram of experimental set-up: HEK293T cells were transiently transfected with a plasmid encoding firefly luciferase under the control of the SRE promoter, and a control renilla luciferase plasmid. Additionally, a plasmid encoding for MEK1$^{DD}$ or mutants were added to trigger an ERK-specific response. **B** When ERK is activated through MEK1$^{DD}$, it phosphorylates multiple substrates, including Elk1, which in turn will form a complex with Signal Response Factor (SRF) and stimulate the Signal Response Element (SRE) promoter to trigger the firefly luciferase expression. The activity of ERK2 can be directly monitored through luminescence measurements. **C** Table presenting the tested MEK1$^{DD}$ mutants. **D** Plotted luciferase activity as percentage of activity relative to MEK1$^{DD}$ for mutants and control experiments. The '+' indicates the use of the inducible SRE promoter vector, while the '-' indicates use of non-inducible control vector. For each condition the mean and standard error are shown as bars, with the individual data points from three different experiments plotted as hexagons. The relative luminescence (RL) used to calculate relative activity are shown in Table S3. **E** Cartoon representation of MEK1$^{DD}$ highlighting the Δinsert mutant and zoom of linker with A-helix specific mutants.

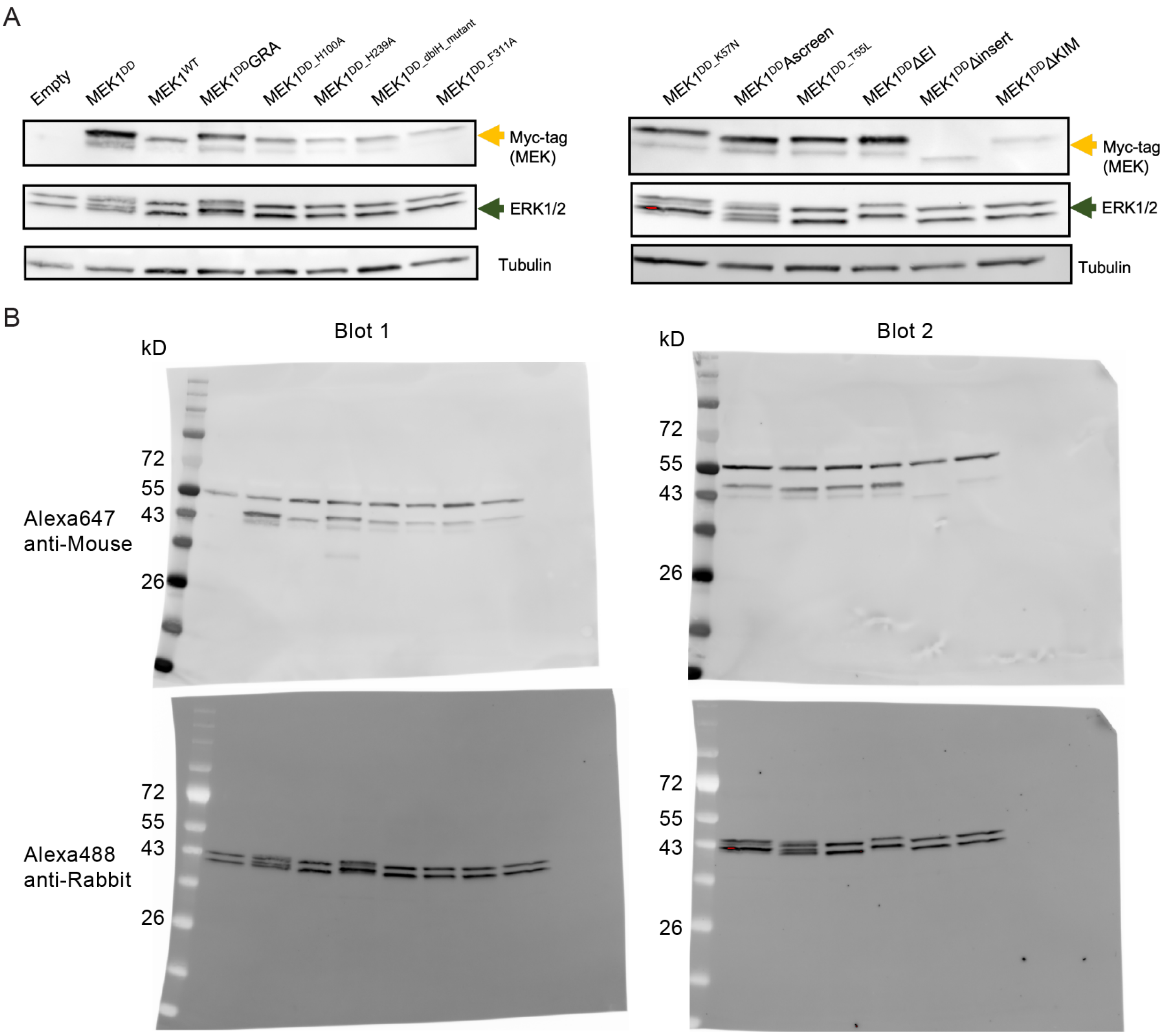


**Fig. S11. Protein expression monitored by Western blots. A** Expression of MEK mutants validated through anti-Myc antibody of transiently transfected tagged proteins. There are differences in expression levels of MEK constructs but those do not correlate to changes in signalling observed. The presence of endogenous ERK1 and 2 was confirmed by blots. **B** The uncropped blots presented in A.

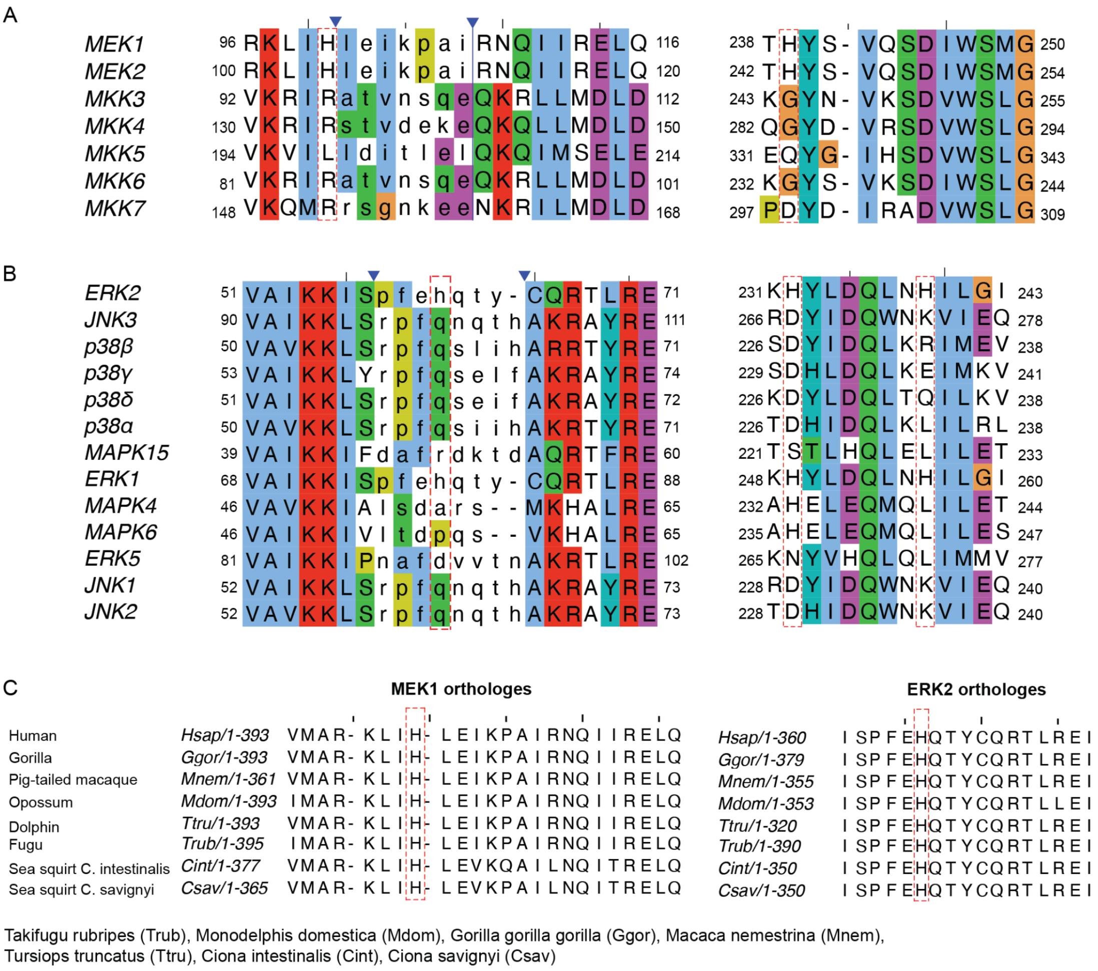


**Fig. S12. Histidine conservation in human MAPKs, and MAP2Ks and across metazoan MEK1 and ERK2 orthologs. A** Alignment of human MAP2Ks from KinCore (*111*) in Clustal coloring. Blue triangle indicates collapsed columns from alignment gaps. For the first block, red dashed box indicates H100 in MEK1 (MAP2K1), the second H239. Both residues interact with ERK2 substrate and are present only in MEK1 and MEK2 (MAP2K2). **B** As for A, alignment from KinCore of human MAPKs in Clustal coloring showing conservation for H61 (left block), and H232 and 239 (right block) for interaction sites. Sites conserved only in ERK2 (MAPK1) and ERK1 (MAPK3). **C** Alignment of MEK1 and ERK2 orthologues showing the conservation of the histidines in metazoans.

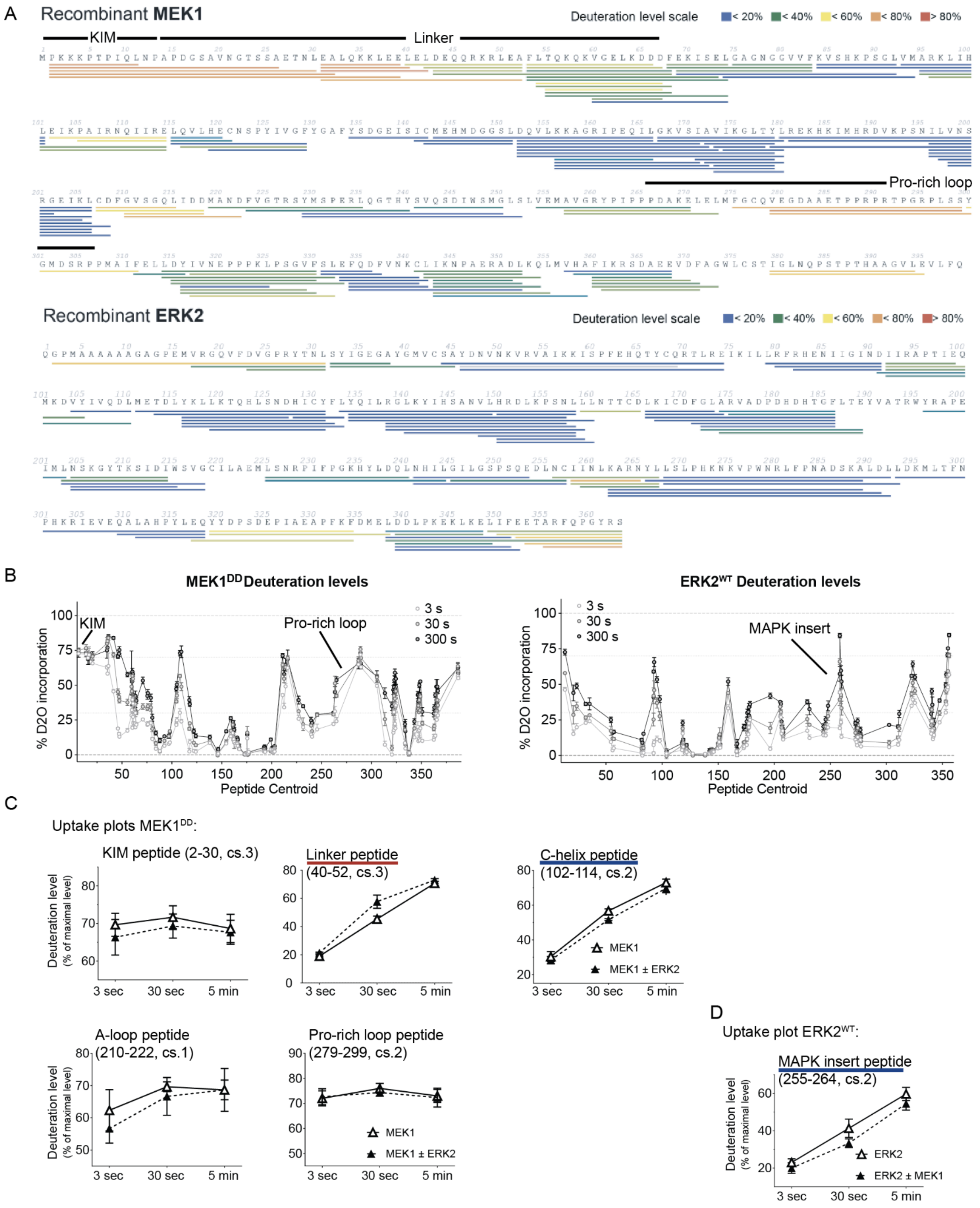


**Fig. S13. HDX-MS analysis of MEK1 upon addition of ERK2 (left) and ERK2 upon addition of MEK1 (right) A** Peptide map: Each line represents a high-quality peptide that was analysed to measure its H/D exchange rate. Peptides are colored according to their dynamics following the legend, with highly dynamic peptides shown in red and protected peptides colored in blue. Protein sequence is from MEK1$^{DD}$ (top) or ERK2$^{WT}$ (bottom) respectively. High exchange rates are indication for flexible regions, relevant flexible MEK1 regions are labelled. **B** Deuteration levels for MEK1$^{DD}$ (left) and ERK2$^{WT}$ (right): Overall deuteration graph showing percentage deuterium incorporation for each peptide analysed. Each dot represents a peptide, plotted according to its central residue (peptide centroid). Percentages are calculated based on a theoretical maximal level. Residue numbering refers to endogenous protein and for clarity regions of high exchange are labelled for each protein. **C** Uptake plots of peptides at three time points for MEK1$^{DD}$ alone and in presence of ERK2$^{WT}$. Data shown from three independent experiments and mean presented with standard deviation. The cs indicates charged state. Peptides with significant changes are marked red or blue in case of increased exposure or shielding respectively. **D** Uptake plot for shielded peptide of ERK2$^{WT}$ in the presence of MEK1$^{DD}$. Data shown from three independent experiments and mean presented with standard deviation. The cs indicates charged state.

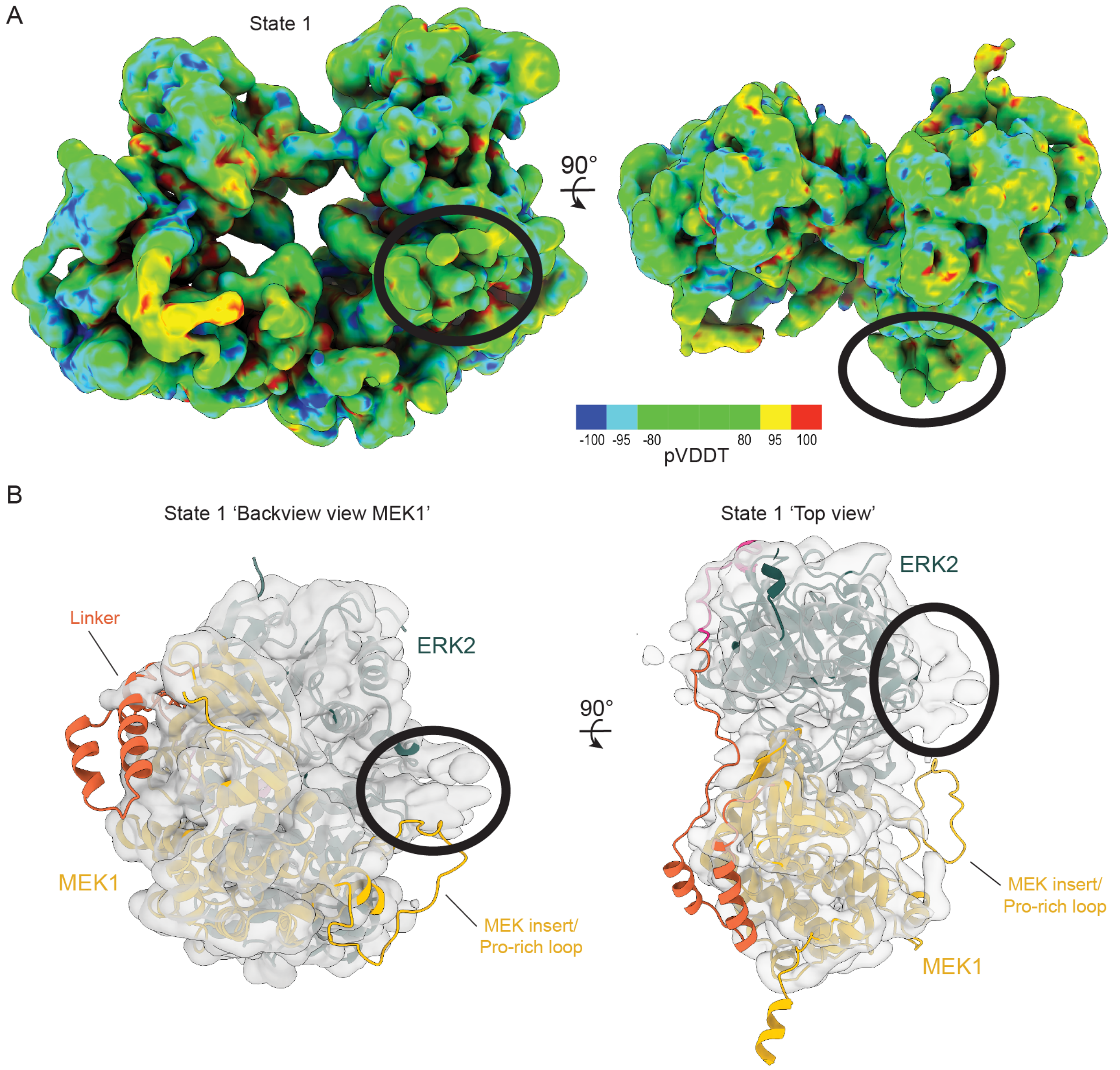


**Fig. S14. Proline-rich loop additional signal. A** Coulomb potential map of state 1 after LocScale run in feature enhance mode. Colored by pVDDT, where regions colored in green indicate regions of high confidence, while those shaded blue and red have significant phase shifts. Black circle indicates additional signal in the map. **B** Same map as A, but transparent with composite model fit to highlight additional signal in proximity of MEK1s proline-rich loop.

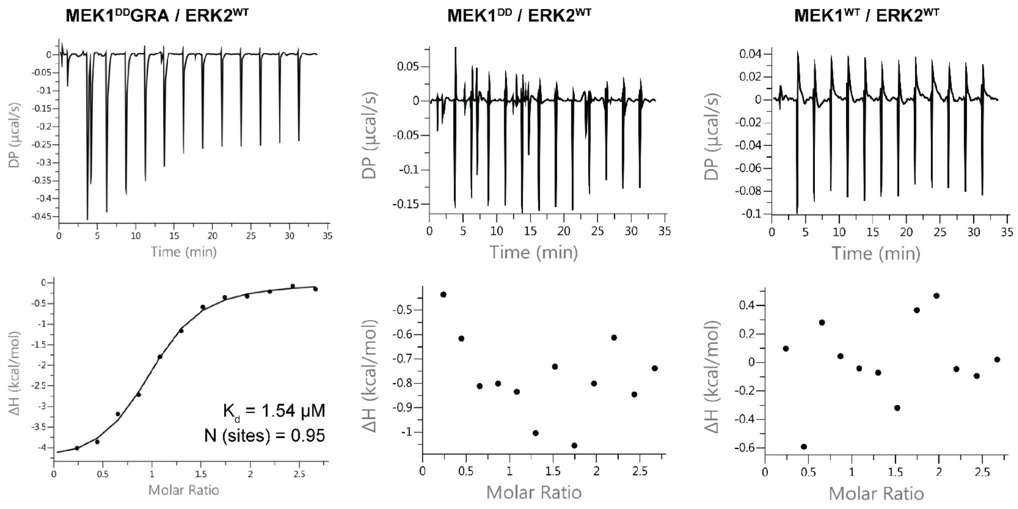


**Fig. S15. ITC data for MEK1 binding to ERK2$^{WT}$.** The left panel shows MEK1$^{DD}$GRA titrated against ERK2$^{WT}$ with measured $K_d$ and N (sites). The calculated mean $K_d$ of four independent experiments was 1.94 μM. The middle and right panels show titrations of MEK1$^{DD}$ and MEK1$^{WT}$ against ERK2$^{WT}$, respectively. These are representative binding curves for experiments. The full list of experiments with calculated thermodynamic parameters is presented in Table S5. All experiments were performed at 20 °C.

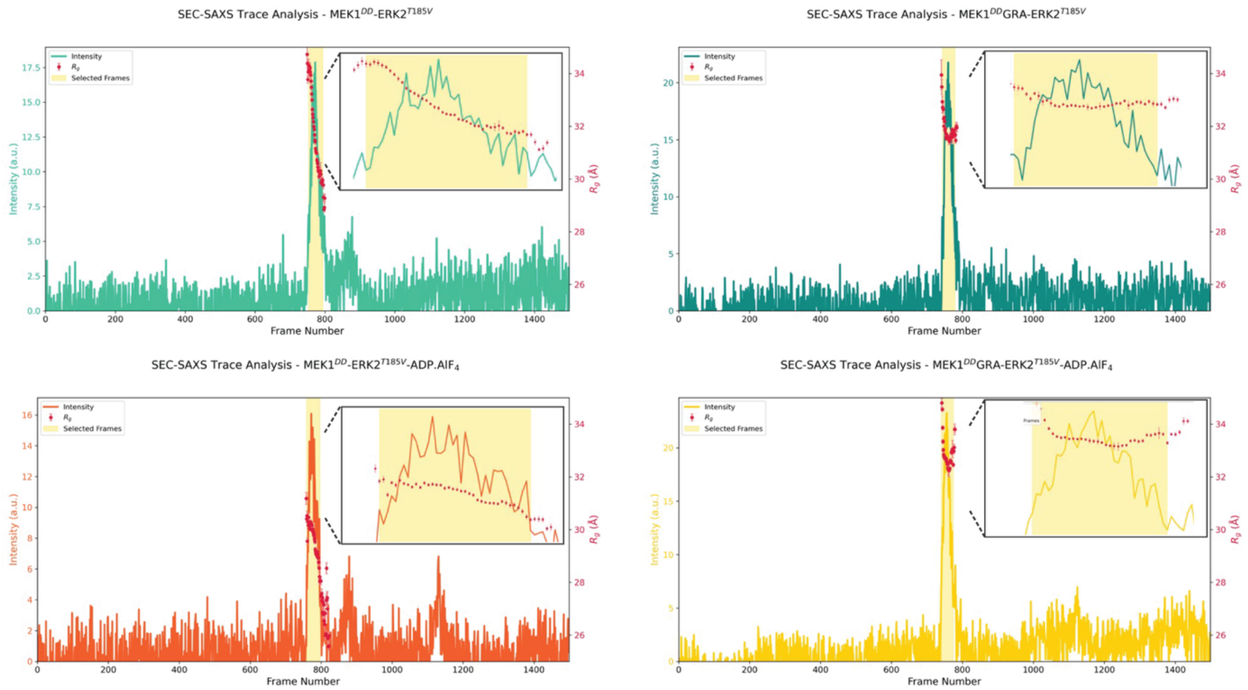


**Fig. S16. SAXS results on complexes without addition of nucleotide.** SEC-SAXS traces are shown for MEK1-ERK2 complex samples run over a Superdex 200 Increase 10/300: MEK1$^{DD}$-ERK2$^{T185V}$ (light green), MEK1$^{DD}$GRA-ERK2$^{T185V}$ (dark green), MEK1$^{DD}$-ERK2$^{T185V}$ with ADPAlF$_4^-$ (orange) and MEK1$^{DD}$GRA-ERK2$^{T185V}$ with ADPAlF$_4^-$ (yellow). Red dots indicate $R_g$ for given frame and yellow shading highlights merged frames for scattering curves presented in Extended Data Fig. 8. Zoom inlets show peak frames and change of $R_g$.

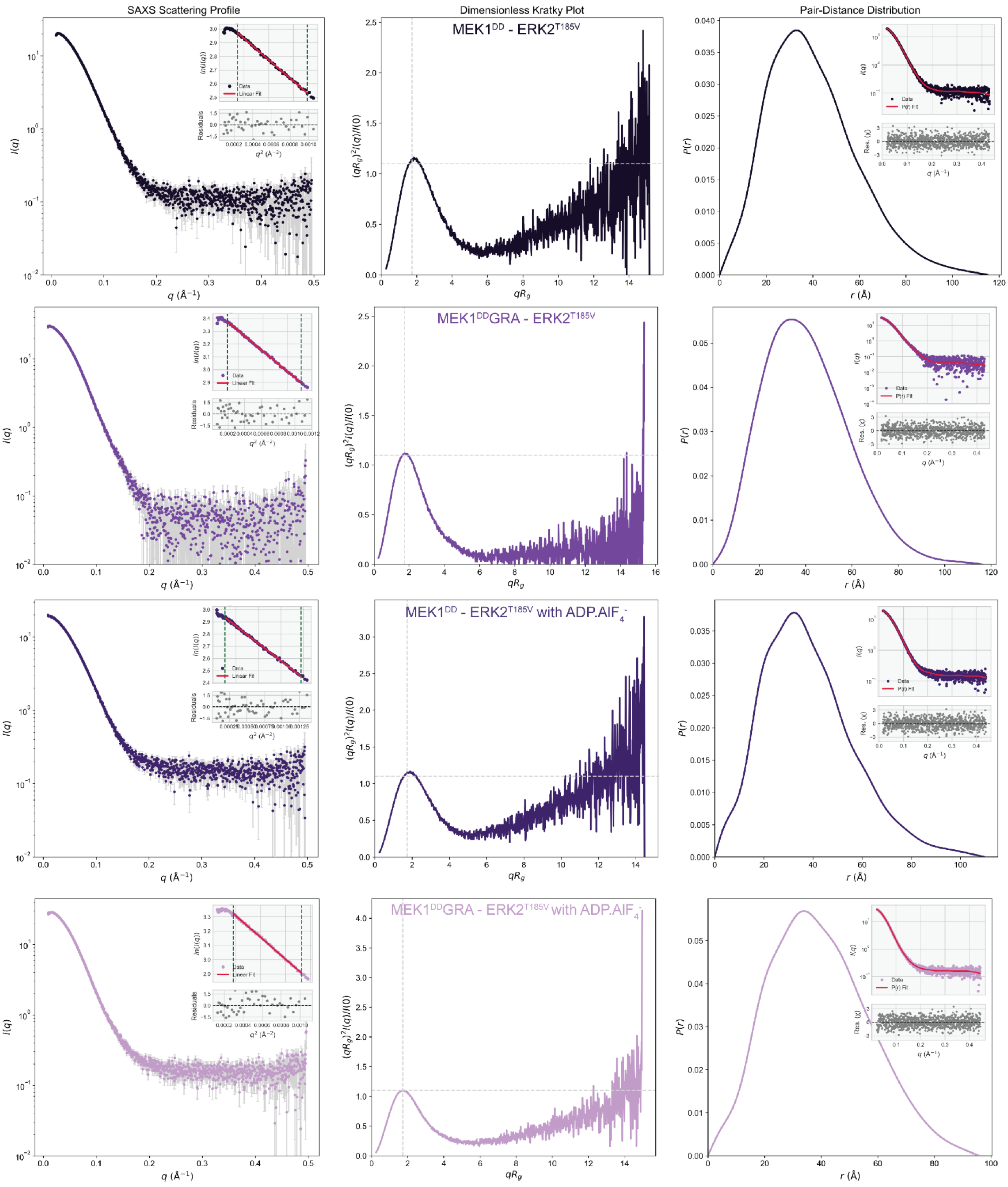


**Fig. S17. SAXS results on complexes with ADP.AlF$_4^-$.** From left to right: Scatter profile, dimensionless Kratky plot, and P(r) distribution for MEK1$^{DD}$-ERK2$^{T185V}$, MEK1$^{DD}$GRA-ERK2$^{T185V}$, MEK1$^{DD}$-ERK2$^{T185V}$ with ADP.AlF$_4^-$ and MEK1$^{DD}$GRA-ERK2$^{T185V}$ with ADP.AlF$_4^-$ (darkest to lightest purple).

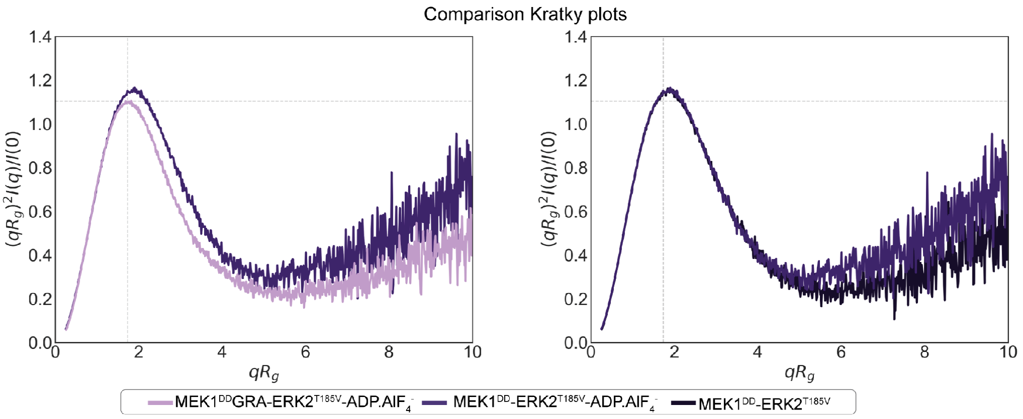


**Fig. S18. SAXS result comparison of complexes.** Bottom row shows dimensionless Kratky plot comparison of two samples not shown in Fig 3d: MEK1$^{DD}$GRA-ERK2$^{T185V}$ with ADPAlF$_4^-$ to MEK1$^{DD}$-ERK2$^{T185V}$ with ADPAlF4$^-$ (left), and MEK1$^{DD}$-ERK2$^{T185V}$ to MEK1$^{DD}$-ERK2$^{T185V}$ with ADPAlF$_4^-$ .

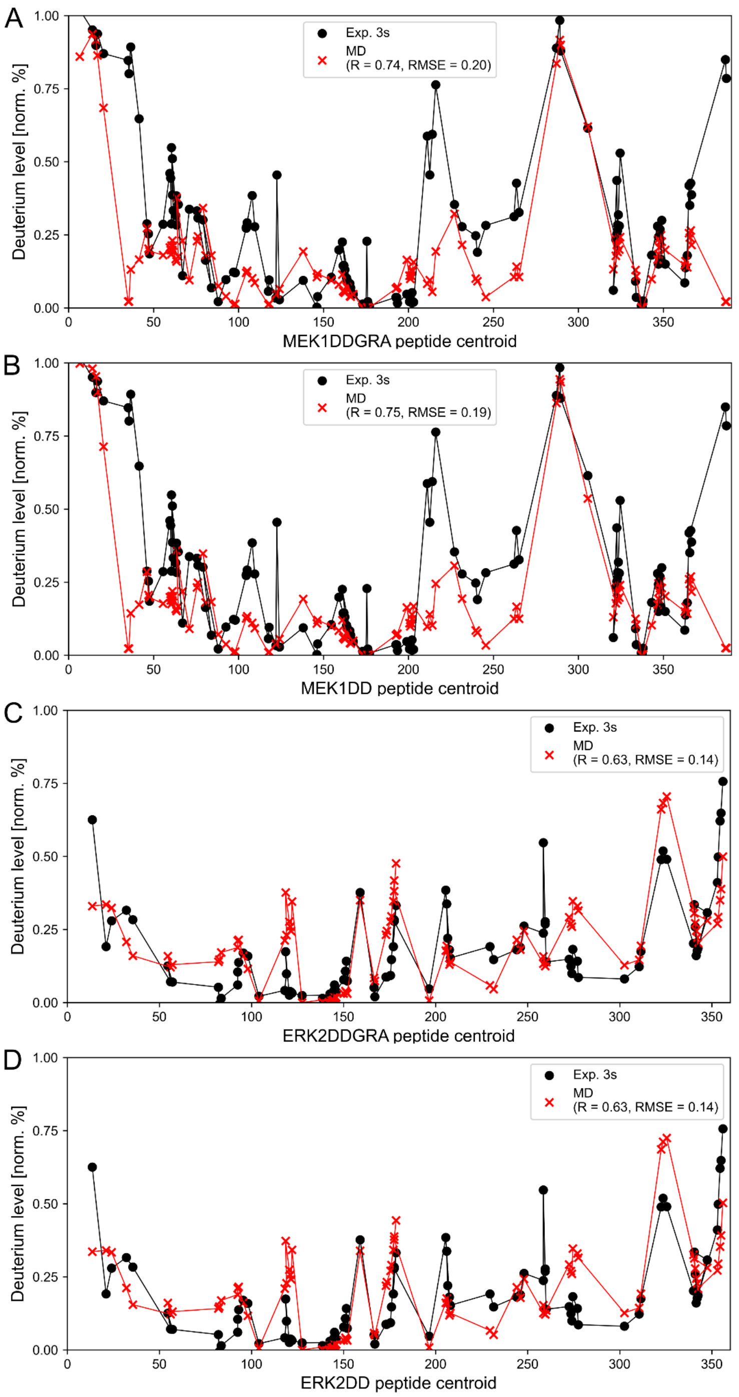


**Fig. S19. HDX-MS profiles comparison.** Experimental (black dots and lines) vs. MD predicted (red crosses and lines) HDX-MS profiles for **A** MEK1$^{DD}$GRA, **B** MEK1$^{DD}$, **C** ERK2$^{WT}$ from simulations with MEK1$^{DD}$GRA and **D** ERK2$^{WT}$ from simulations with MEK1$^{DD}$. The profiles are predicted from the full trajectories (48 µs, 48000 frames). The deuteration level is normalized and the experimental data is rescaled to approximate a 100% maximum theoretical coverage. The last two points of the MEK1 datasets are reported but not used in the fit as the C-terminal domain of MEK1 was not simulated.

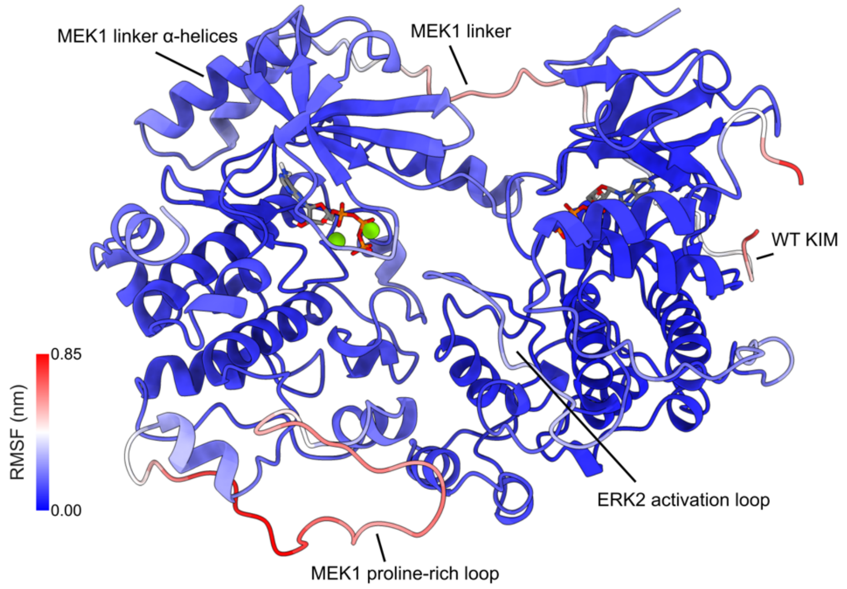


**Fig. S20. Average root mean square fluctuation (RMSF) reprojected as a heatmap on the MEK1$^{DD}$-ERK2$^{WT}$ complex for all models simulated.** The average RMSF correlates well with HDX-MS data (Fig. S19) and with pLDDT scores from the AF3 prediction (Fig. S1).

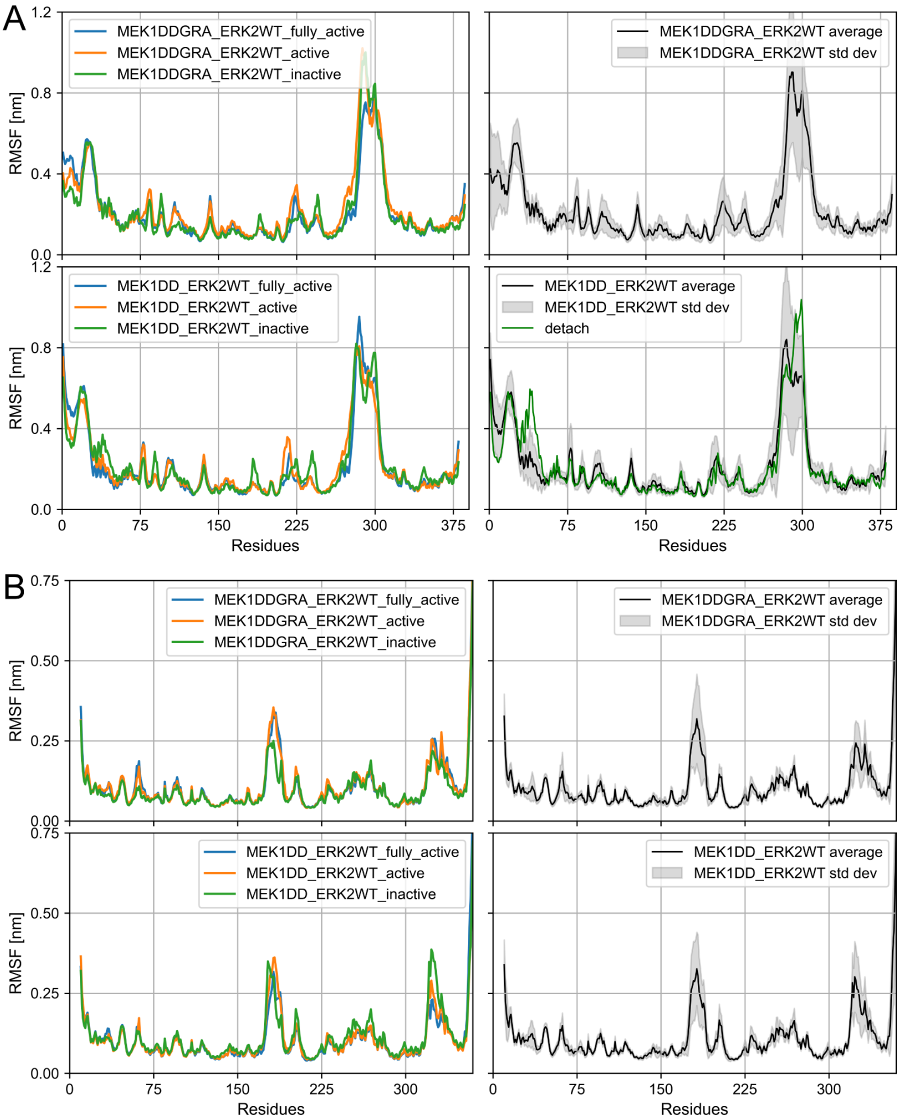


**Fig. S21. RMSF for all the MD simulations of the complex with respect to their corresponding starting configurations. A** RMSF for MEK1$^{DD}$GRA (top row) and MEK1$^{DD}$ (bottom row) for the three individual starting states (left, 16 µs each) and for all the states together (right, 48 µs total). **B** Same as A but for ERK2$^{WT}$. In the last panel of A the green line *detach* is one of the simulations where the linker shows large rearrangements (as plotted in Fig. 4B of the main text).

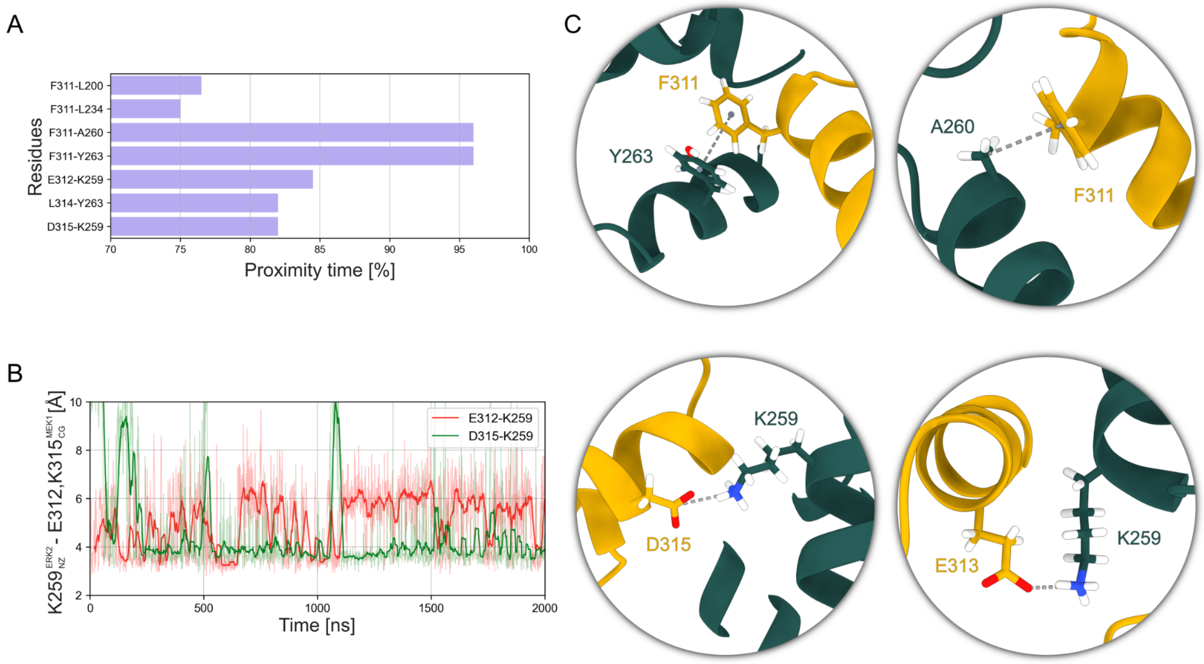


**Fig. S22. Analysis of the C-lobe domains interface. A** Contact map between MEK1 and ERK2 at the C-lobe domains interface. The bar plot shows the most common residues that are in close proximity (< 4.5 Å) for most of the simulation time (> 75%, computed on a total of 96 µs). **B** Distance between the nitrogen atom in ERK2$^{WT}$ K259 ammonium group and the β-carbon of MEK1$^{DD}$ E312 and K315. A salt bridge is transiently formed throughout the simulation. **C** Snapshots from MD simulations showing key main interactions from panel A.

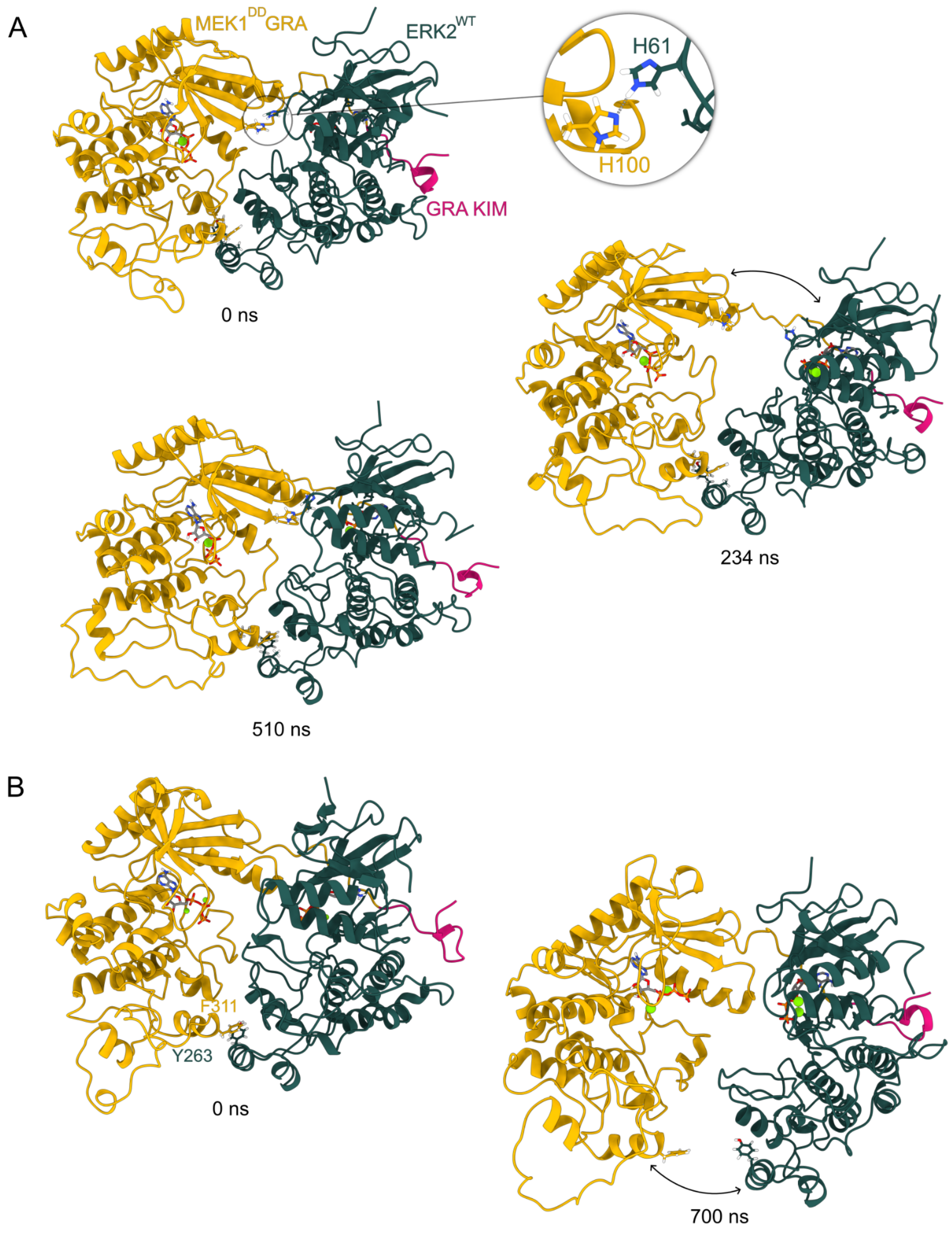


**Fig. S23. N- and C-lobe interfaces behaviour. A** Breaking and reforming of the interface at the N-lobe domains over time. **B** Breaking of the interface at the C-lobe domains over time. In both cases the KIM (GRA here) helps in keeping the complex together.

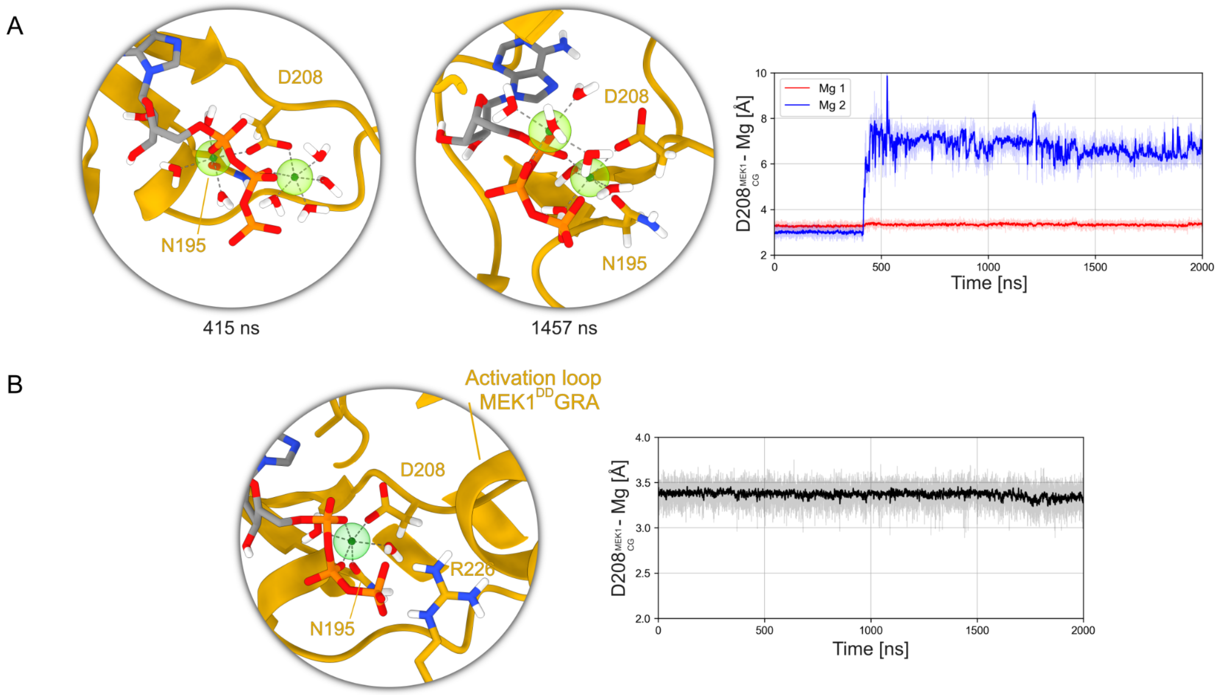


**Fig. S24. Mobility of magnesium ions in MEK1 binding site. A** Magnesium ions in the binding site of a fully active structure of MEK1$^{DD}$GRA. The ions stay coordinated to the ATP, but one of the two leaves the coordinating oxygen in D208 of MEK1 DFG domain as shown in the time series plot. **B** Magnesium ion in the binding site of an inactive structure of MEK1$^{DD}$GRA. This configuration is extremely stable and the ion does not leave in any replicate (totalling 32 μs). Ions are displayed as semi-transparent green spheres and dashed lines indicate the six coordinating oxygen atoms of nearby residues or water molecules forming the typical octahedral geometry.

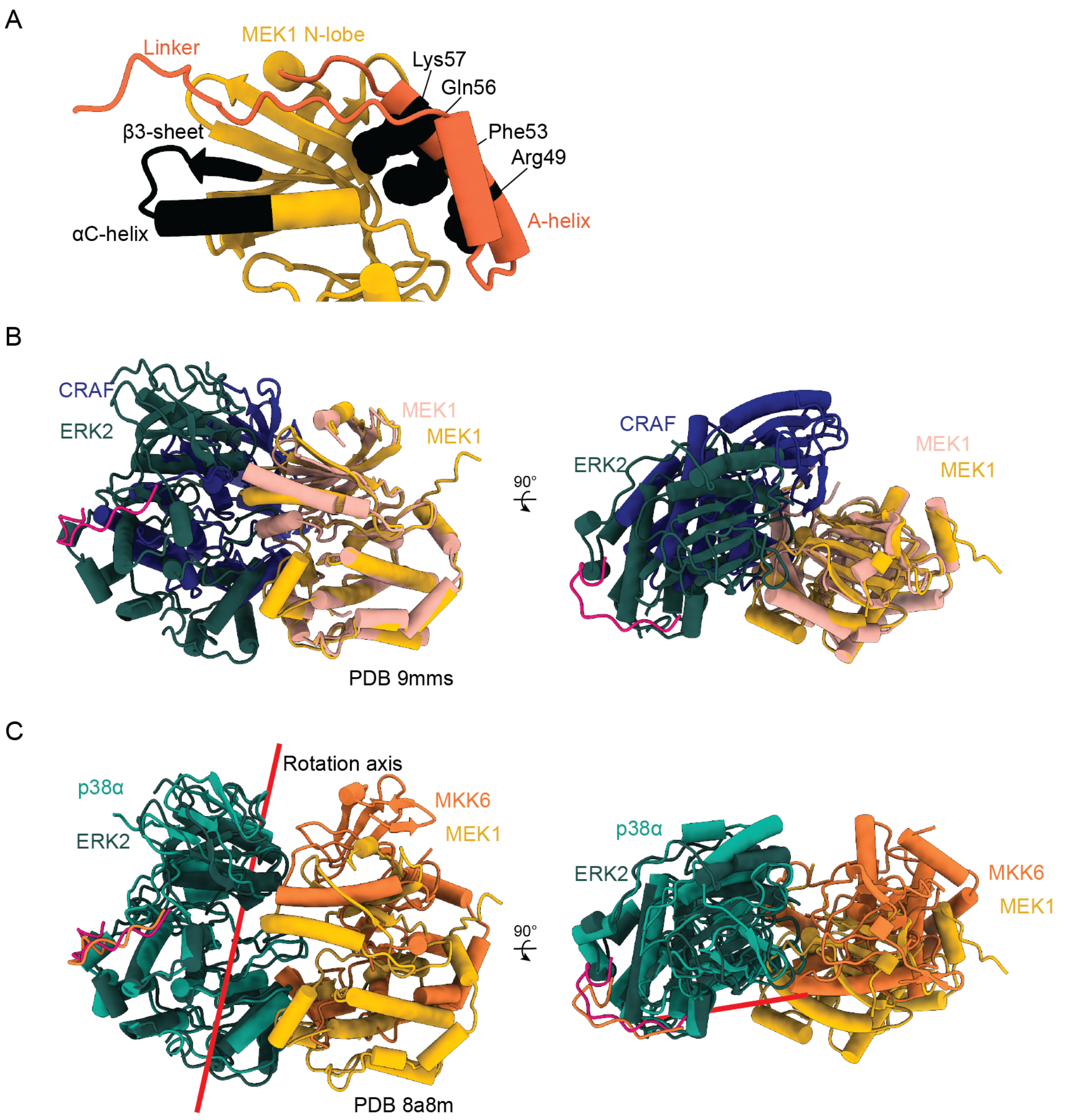


**Fig. S25. MEK1 cancer mutations and comparison of MAPK complexes. A,** Sites of frequent mutations isolated from patient data (*26*) mapped on MEK1 state 2 model. Affected residues are colored black. For the β3-αC loop, mostly in-frame deletions occur whereas for the A-helix missense mutations are recorded. Affected residues on the A-helix are displayed as spheres. **B** Overlay of the MEK1$^{DD}$GRA-ERK2$^{T185V}$ complex with the CRAF-MEK1 complex aligned on MEK1. The interface between MEK1's activating kinase and its substrate is completely different. **C** Overlay of the MAP2K-MAPK complexes on the substrate MAPK. The axis of rotation about which MEK1 must be moved in order to be in the same position as MKK6 is shown as a red line.

**Table S1. X-ray crystallographic data processing and refinement statistics.**

| **Structure** | **ERK1-GRA24KIM1 (PDB-9TU0)** |
|---|---|
| Raw Data DOI | https://doi.org/10.15151/esrf-dc-2310714581 |
| Beamline | MASSIF-1 (ESRF) |
| Space group | *P*6522 |
| Wavelength | 0.966 Å |
| Unit cell dimensions (Å) a, b, c | 59.82, 59.82, 388.67 |
| α, β, γ (°) | 90.0, 90.0, 120.0 |
| Resolution range (Å) | 20-2.17 |
| Reflections | 609516 (32486) |
| Number of unique reflections | 17953 (899) |
| Multiplicity[1] | 34 (36) |
| Completeness[1] (spherical, %) | 77.0 (16.2) |
| Completeness[1] (ellipsoidal, %) | 91.6 (49.0) |
| Rmerge[1] | 0.18 (2.6) |
| <I/s(I)>[1] | 18.5 (1.9) |
| CC(1/2) | 0.99 (0.91) |
| Wilson B factor | 42.2 $Å^2$ |
| No. of atoms | |
| Protein | 2891 |
| Ligands/ion | 15 |
| Water | 70 |
| R-factor (%) | 23.2 |
| Free R-factor (%) | 28.7 |
| RMS deviations: | |
| Bonds (Å) | 0.004 |
| Angles (°) | 0.657 |
| Ramachandran | |
| Favoured (%) | 94.58 |
| Allowed (%) | 5.42 |
| Outliers (%) | 0 |

[1] Statistics for the highest resolution bin (2.38-2.17) are shown in parenthesis

**Table S2. Cryo-EM data collection, processing and refinement.**

| Data collection and processing | MEK1$^{DD}$GRA-ERK2$^{T185V}$ state 1 (PDB-9TYG) (EMDB-56418) | MEK1$^{DD}$GRA-ERK2$^{T185V}$ state 2 (PDB-9TYH) (EMDB-56419) | MEK1$^{DD}$GRA-ERK2$^{T185V}$ state 3 (PDB-9TYI) (EMDB-56420) |
|---|---|---|---|
| Microscope | Titan Krios G4 | | |
| Camera | Falcon 4i | | |
| Voltage (kV) | 300 | | |
| Magnification | 270,000 | | |
| Pixel size (Å) | 0.46 | | |
| Electron exposure ($e^-$/Å$^2$) | 50.0 | | |
| Exposure rate ($e^-$/pixel/sec) | 6.1 | | |
| Number of frames per exposure | 58 | | |
| Defocus range (µm) | -1.0 to -2.2 | | |
| Automation software | EPU | | |
| Micrographs collected (no.) | 88517 | | |
| Micrographs used | 74698 | | |
| Total extracted particles (no.) | 10,714,101 | | |
| Refined particles (no.) | 977,550 | 977,550 | 977,550 |
| Final particle images (no.) | 142,124 | 113,564 | 93,925 |
| Map resolution (Å) | | | |
| FSC 0.5 (Å unmasked/masked) | 4.7/3.5 | 5.6 /4.1 | 6.0/4.4 |
| FSC 0.143 (Å unmasked/masked) | 3.8/2.99 | 3.9/ 3.5 | 4.3/3.6 |
| Map resolution range (Å) | 2.6-3.4 | 2.8-3.8 | 3.0-4.1 |
| Map sharpening factor (Å$^2$) | -25 | -135 | -120 |
| Map sharpening method | cryoSPARC | cryoSPARC | cryoSPARC |
| **Model composition** | | | |
| Initial models used | 9TU0, AF3 | State 1, AF-Q02750-K1A | State 2, AF-Q02750-K1A |
| Non-hydrogen atoms | 5235 | 5111 | 4971 |
| Protein residues | 645 | 628 | 617 |
| Ligands | 3 | 2 | 0 |
| **Refinement** | | | |
| Refinement package | Servalcat, phenix | phenix | phenix |
| Real or reciprocal space | Real and reciprocal | real | real |
| Resolution cutoff (Å) | 3.0 | 3.5 | 3.6 |
| Model-Map scores | | | |
| CC | 0.73 | 0.75 | 0.72 |
| FSC 0.5 (Å) | 3.53 | 4.2 | 4.5 |
| B factors (Å$^2$) | | | |
| Protein | 70 | 135 | 150 |
| Ligands | 174 | 243 | - |
| R.M.S deviations | | | |
| Bond length (Å) | 0.003 | 0.003 | 0.003 |
| Bond angles (°) | 0.718 | 0.733 | 0.665 |
| Validation | | | |

| | | | |
|---|---|---|---|
| MolProbity score | 1.57 | 1.77 | 1.83 |
| Clash score | 6.30 | 7.53 | 7.41 |
| Poor rotamers (%) | 0.52 | 1.60 | 1.08 |
| Ramachandran plot | | | |
| Favoured (%) | 96.55 | 96.76 | 94.07 |
| Allowed (%) | 3.45 | 3.24 | 5.93 |
| Outliers (%) | 0 | 0 | 0 |

**Table S3. Reporter assay relative luminescence (RL) for all measured samples and controls.**

| | Experiment 1 | Experiment 2 | Experiment 3 | Passed normality test? (Shapiro-Wilk statistics) | Paired t test for significant difference to MEK1$^{DD}$ (p-value) |
|---|---|---|---|---|---|
| **Condition** | **RL**[1] | **RL** | **RL** | | |
| **Empty +**[2] | 0.052 | 0.147 | 0.154 | Yes (0.999) | Yes (0.0209) |
| **Empty -** | 0.043 | 0.111 | 0.119 | - | - |
| **MEK1$^{DD}$ +**[3] | 9.578 | 5.813 | 7.711 | Yes (1.000) | - |
| **MEK1$^{DD}$ -** | 0.158 | 0.32 | 0.302 | - | - |
| **MEK1$^{WT}$ +** | 0.067 | 0.157 | 0.149 | Yes (1.000) | Yes (0.0209) |
| **MEK1$^{WT}$ -** | 0.074 | 0.198 | 0.223 | - | - |
| **MEK1$^{DD}$GRA +** | 6.757 | 5.330 | 5.732 | Yes (0.9746) | No (0.1234) |
| **MEK1$^{DD}$GRA -** | 0.179 | 0.362 | 0.289 | - | - |
| **MEK1$^{DD-H100A}$ +** | 6.238 | 4.214 | 4.869 | Yes (0.9425) | Yes (0.0376) |
| **MEK1$^{DD-H239A}$ +** | 7.743 | 5.943 | 8.203 | Yes (0.8634) | No (0.6322) |
| **MEK1$^{DD}$dblH-mutant +** | 7.073 | 4.576 | 5.131 | Yes (0.7917) | Yes (0.0402) |
| **MEK1$^{DD-F311A}$ +** | 5.804 | 4.348 | 4.265 | Yes (0.8541) | No (0.0568) |
| **MEK1$^{DD-K57N}$ +** | 12.555 | 6.72 | 11.042 | Yes (0.8569) | No (0.0862) |
| **MEK1$^{DD}$Ascreen +** | 13.299 | 9.503 | 9.925 | No (0.7654) | |
| **MEK1$^{DD-T55L}$ +** | 7.482 | 4.084 | 5.456 | Yes (0.9505) | Yes (0.0059) |
| **MEK1$^{DD}$Δinsert +** | 5.002 | 2.418 | 2.663 | Yes (0.9422) | Yes (0.0126) |
| **MEK1$^{DD}$ΔEI +** | 11.307 | 8.699 | 8.497 | Yes (0.9966) | No (0.0974) |
| **MEK1$^{DD}$ΔKIM +** | 2.291 | 1.096 | 0.975 | Yes (0.9019) | Yes (0.0153) |

[1]Ratio calculated from firefly/renilla luminescence, averaged from technical triplicates.

[2]empty pcDNA3.1 vector transfected with non-inducible vector, used as negative control

[3]set as positive control to calculate relative activity

**Table S4. Calculated relative activity to $MEK1^{DD}$ with mean, SD, and SE.**

| | **Experiment 1** | **Experiment 2** | **Experiment 3** | **Descriptive statistics** | | |
|---|---|---|---|---|---|---|
| **Condition** | Relative activity[1] | Relative activity | Relative activity | Mean | SD | SEM |
| **Empty +** [2] | 0 | 0 | 0 | 0.0 | 0.0 | 0.0 |
| **$MEK1^{DD}$ +** [3] | 100 | 100 | 100 | 100 | 0 | 0 |
| **$MEK1^{DD}$ -** | 1.1 | 3.0 | 2 | 2.0 | 0.95 | 0.55 |
| **$MEK1^{WT}$ +** | 0.2 | 0.2 | -0.1 | 0.085 | 0.16 | 0.093 |
| **$MEK1^{WT}$ -** | 0.2 | 0.9 | 0.9 | 0.67 | 0.39 | 0.23 |
| **$MEK1^{DD}$GRA +** | 70.4 | 91.5 | 73.8 | 79 | 11 | 6.5 |
| **$MEK1^{DD}$GRA -** | 1.3 | 3.8 | 1.8 | 2.3 | 1.3 | 0.76 |
| **$MEK1^{DD\text{-}H100A}$ +** | 64.9 | 71.8 | 62.4 | 66 | 4.9 | 2.8 |
| **$MEK1^{DD\text{-}H239A}$ +** | 80.7 | 102.3 | 106.5 | 97 | 14 | 8.0 |
| **$MEK1^{DD}$dblH-mutant +** | 73.7 | 78.2 | 65.9 | 73 | 6.2 | 3.6 |
| **$MEK1^{DD\text{-}F311A}$ +** | 60.4 | 74.1 | 54.4 | 63 | 10 | 5.8 |
| **$MEK1^{DD\text{-}K57N}$ +** | 131.3 | 116.0 | 144.1 | 130 | 14 | 8.1 |
| **$MEK1^{DD}$Ascreen +** | 139.1 | 165.1 | 129.3 | 144 | 19 | 11 |
| **$MEK1^{DD\text{-}T55L}$ +** | 78.0 | 69.5 | 70.2 | 73 | 4.7 | 2.7 |
| **$MEK1^{DD}$ΔEI +** | 118.2 | 150.9 | 110.4 | 126 | 21 | 12 |
| **$MEK1^{DD}$Δinsert +** | 52.0 | 40.1 | 33.2 | 42 | 9.5 | 5.5 |
| **$MEK1^{DD}$ΔKIM +** | 23.5 | 16.7 | 10.9 | 17 | 6.3 | 3.6 |

[1]Calculated by (condition ratio - negative control ratio)/(positive control ratio - negative control ratio).
[2]empty pcDNA3.1 vector transfected with active reporter, used as negative control
[3]set as positive control to calculate relative activity

**Table S5. ITC measurements of MEK1-ERK2 and ERK1/2-GRA24 KIM1 peptide affinities.** Summary of ITC parameters obtained for binding of $ERK2^{WT}$ to different MEK1 constructs and the KIM peptide of MEK1 or effector protein GRA24. (n.d for not determined, in cases of no binding or very weak binding.)

| **Rep** | **Interaction** | **[Syringe] (µM)** | **[Cell] (µM)** | **N (sites)** | **$K_d$ (µM)** | **dH (kcal/mol)** | **dG (kcal/mol)** | **-TdS (kcal/mol)** |
|---|---|---|---|---|---|---|---|---|
| 1 | $MEK1^{DD}GRA/ERK2^{WT}$ | 280 | 30 | 0.79 | 3.21 | -5.81 | -7.37 | -1.56 |
| 2 | $MEK1^{DD}GRA/ERK2^{WT}$ | 280 | 20 | 0.97 | 2.30 | -5.40 | -7.56 | -2.16 |
| 3 | $MEK1^{DD}GRA/ERK2^{WT}$ | 280 | 20 | 0.95 | 1.54 | -4.47 | -7.80 | -3.33 |
| 4 | $MEK1^{DD}GRA/ERK2^{WT}$ | 280 | 20 | 0.90 | 0.718 | -4.78 | -8.25 | -3.47 |
| 1 | $MEK1^{DD}/ERK2^{WT}$ | 280 | 20 | n.d | n.d | n.d | n.d | n.d |
| 1 | $MEK1^{WT}/ERK2^{WT}$ | 280 | 20 | n.d | n.d | n.d | n.d | n.d |
| 1 | $ERK2^{WT}$/GRA24 KIM | 280 | 30 | 0.75 | 0.260 | -9.36 | -8.84 | 0.52 |
| 2 | $ERK2^{WT}$/GRA24 KIM | 280 | 30 | 0.78 | 0.305 | -9.49 | -8.74 | 0.75 |
| 3 | $ERK2^{WT}$/GRA24 KIM | 280 | 30 | 0.76 | 0.376 | -9.59 | -8.62 | 0.97 |
| 1 | $ERK2^{WT}$/MEK1 KIM | 280 | 30 | n.d | n.d | n.d | n.d | n.d |
| 2 | $ERK2^{WT}$/MEK1 KIM | 600 | 30 | n.d | n.d | n.d | n.d | n.d |
| 1 | $ERK1^{WT}$/GRA24 KIM | 200 | 20 | 0.774 | 0.338 | -11.2 | -8.68 | 2.5 |
| 2 | $ERK1^{WT}$/GRA24 KIM | 200 | 20 | 0.856 | 0.324 | -10.9 | -8.71 | 2.18 |
| 3 | $ERK1^{WT}$/GRA24 KIM | 250 | 25 | 0.671 | 0.272 | -12.8 | -8.81 | 3.97 |

**Table S6 - SAXS data collection and processing**

Sample details

| Sample name | $MEK1^{DD}$-$ERK2^{T185V}$ | $MEK1^{DD}$GRA-$ERK2^{T185V}$ | $MEK1^{DD}$-$ERK2^{T185V}$-ADP.$AlF_4$ | $MEK1^{DD}$GRA-$ERK2^{T185V}$-ADP.$AlF_4$ |
|---|---|---|---|---|
| Source | | | | |
| Scattering particle composition | $C_{3848}H_{6074}N_{1036}O_{1119}S_{35}$ | $C_{3880}H_{6120}N_{1044}O_{1129}S_{35}$ | $C_{3848}H_{6074}N_{1036}O_{1119}S_{35}$ | $C_{3880}H_{6120}N_{1044}O_{1129}S_{35}$ |
| Mw from chem comp (kDa) | 85876.93 | 86579.69 | 85876.93 | 86579.69 |
| Concentration (mg/ml) | 10 | 10 | 10 | 10 |
| Buffer composition | 50 mM HEPES pH 7.5, 200 mM NaCl, 10 mM $MgCl_2$, 2.5% glycerol, 0.5 mM TCEP | | | |

SAXS data collection parameters

| - | $MEK1^{DD}$-$ERK2^{T185V}$ | $MEK1^{DD}$GRA-$ERK2^{T185V}$ | $MEK1^{DD}$-$ERK2^{T185V}$-ADP.$AlF_4$ | $MEK1^{DD}$GRA-$ERK2^{T185V}$-ADP.$AlF_4$ |
|---|---|---|---|---|
| Date of collection | 25/06/2025 | | | |
| Instrument | ESRF BioSAXS beamline BM29 with Dectris PILATUS3 X 2M detector | | | |
| Wavelength (Å) | 0.99 | | | |
| Beamsize (µm) | 150 x 150 | | | |
| Detector distance (m) | 2.827 | | | |
| q range ($Å^{-1}$) | 0.0085 – 0.4959 | | | |
| Absolute scaling method | Comparison with scattering from $H_2O$ | | | |
| Normalisation | To transmitted intensity by beamstop diode | | | |
| Monitoring for rad. dam. | Data frame-by-frame comparison | | | |
| Exposure time | 2 second continuous exposures | | | |
| Sample configuration | SEC-SAXS with samples exposed in 1mm quartz capillary | | | |
| Sample temperature (°C) | 20 | | | |
| Flow rate (ml/min) | 0.5 | | | |
| Column used | GE Superdex 200 Increase 10/300 | | | |
| Frames collected | 1500 | | | |

Software employed for data reduction, analysis, and interpretation

| | |
|---|---|
| SAXS data reduction | FreeSAS (Kieffer et al, JSR, 2022) |
| Basic analysis (Guinier) | BioXTAS RAW (Hopkins., JAC, 2024) |
| Mw calculation | Bayes (Hajizadeh., Sci. Rep., 2018), Vc (Rambo., Nature, 2013), Vp (Piiadov, Prot. Sci. 2019), |
| P(r) | BIFT (Hansen., JAC, 2000) |
| Atomistic modelling | PEPSI-SAXS (Grudinin et al, Acta D, 2017), NOLB (Hoffman et al, JCTC, 2017) |

Primary analysis

| | MEK1$^{DD}$-ERK2$^{T185V}$ | MEK1$^{DD}$GRA-ERK2$^{T185V}$ | MEK1$^{DD}$-ERK2$^{T185V}$-ADP.AlF$_4$ | MEK1$^{DD}$GRA-ERK2$^{T185V}$-ADP.AlF$_4$ |
|---|---|---|---|---|
| Frames averaged | 754 - 794 | 744 - 779 | 759 - 796 | 741 - 774 |
| Guinier analysis | | | | |
| I(0) ($cm^{-1}$) | 21.90 ± 0.03 | 31.96 ± 0.04 | 20.24 ± 0.03 | 31.79 ± 0.05 |
| $R_g$ (Å) | 30.58 ± 0.07 | 30.96 ± 0.05 | 29.22 ± 0.07 | 30.21 ± 0.07 |
| $R^2$ | 0.9987 | 0.9992 | 0.9984 | 0.9991 |
| $q_{min}$ (n) | 0.01925 (22) | 0.01681 (17) | 0.01583 (15) | 0.0212 (26) |
| $q_{max}$ (n) | 0.04218 (69) | 0.04218 (69) | 0.04413 (73) | 0.04266 (70) |
| $qR_g$ min | 0.5887 | 0.5205 | 0.4626 | 0.6405 |
| $qR_g$ max | 1.2898 | 1.3060 | 1.2894 | 1.2890 |
| SHANUM | | | | |
| q max | 0.4959 | 0.4959 | 0.4959 | 0.4959 |
| N sh | 16.23 | 16.15 | 14.66 | 15.61 |
| N opt | 15 | 14 | 13 | 14 |
| q opt | 0.4583 | 0.4299 | 0.4397 | 0.4447 |
| Last q | 923 | 865 | 885 | 895 |
| P(r) analysis (BIFT) | | | | |
| $q_{min}$ - $q_{max}$ | 0.0192 – 0.4588 | 0.0168 – 0.4305 | 0.0158 – 0.4402 | 0.0212 – 0.4451 |
| I(0) ($cm^{-1}$) | 22.05 ± 0.03 | 32.10 ± 0.03 | 20.35 ± 0.02 | 31.97 ± 0.02 |
| $R_g$ (Å) | 31.28 ± 0.11 | 31.32 ± 0.06 | 29.87 ± 0.07 | 30.50 ± 0.03 |
| $d_{max}$ (Å) | 114.62 ± 6.85 | 116.47 ± 6.12 | 109.74 ± 5.46 | 95.60 ± 2.81 |
| $\chi^2$ | 1.20 | 1.10 | 1.18 | 1.14 |
| MW calculation | | | | |
| Bayes MW | 67.1 | 91.2 | 55.6 | 74.3 |
| Vc MW | 62.7 | 84.3 | 52.1 | 69.0 |
| Vp MW | 73.0 | 99.6 | 58.4 | 77.8 |

Model fitting

| Sample | MEK1$^{DD}$GRA-ERK2$^{T185V}$ | MEK1$^{DD}$GRA-ERK2$^{T185V}$-ADP.AlF$_4$ |
|---|---|---|
| PEPSI-SAXS - NOLB | | |
| q range for fitting | 0.0168 – 0.4305 | 0.0212 – 0.4451 |
| Initial $\chi^2$ | 4.81 | 5.13 |
| Refined $\chi^2$ | 2.70 | 3.61 |

**Table S7. Molecular dynamics simulations systems.** Summary of all the MD simulation runs for all the systems.

| System | State | # replicas | Time per replica (µs) | Total time per system (µs) |
|---|---|---|---|---|
| MEK1$^{DD}$GRA/ERK2$^{WT}$ | Fully active | 8 | 2 | 16 |
| | Active | 8 | 2 | 16 |
| | Inactive | 8 | 2 | 16 |
| MEK1$^{DD}$/ERK2$^{WT}$ | Fully active | 8 | 2 | 16 |
| | Active | 8 | 2 | 16 |
| | Inactive | 8 | 2 | 16 |
| | | | **Total time (µs)** | **96** |

**Data S1 and S2. (separate file)**
HDX-MS data tables for MEK1$^{DD}$ and ERK2

**Movie S1** - MD simulation trajectory of MEK1$^{DD}$GRA-ERK2$^{WT}$ fully active complex as Y187 on the activation loop of ERK2 forms a catalytical competent state with ATP in the MEK1 binding site. The simulation frames are extracted from the time line reported in Figure 4A.
